# A computationally designed panel of diverse and selective peroxygenases for terpene oxyfunctionalization

**DOI:** 10.1101/2024.09.27.615329

**Authors:** Judith Münch, Jordi Soler, Ofir Gildor-Cristal, Sarel J. Fleishman, Marc Garcia-Borràs, Martin J. Weissenborn

**Author notes:** **Corresponding Author** Martin J. Weissenborn − Leibniz Institute of Plant Biochemistry, 06120 Halle (Saale), Germany; Department of Chemistry, Martin Luther-University Halle-Wittenberg, Weinbergweg 22, 06120, Halle/Saale, Germany. **Corresponding Author** Sarel J. Fleishman − Department of Biomolecular Sciences, Weizmann Institute of Science, Rehovot 7600001, Israel. **Corresponding Author** Marc Garcia-Borràs − Institut de Química Computacional i Catàlisi and Departament de Química, Universitat de Girona, Carrer Maria Aurèlia Capmany 69, Girona 17003, Catalonia, Spain.

## Abstract

Enzyme engineering has a critical role in the transition to economical, low-energy and environmentally friendly chemical production. Current approaches relying on costly iterations of mutation and selection are limited to reactions with a straightforward experimental readout and struggle to address mutational epistasis. We focus on unspecific peroxygenases (UPOs), prized engineering targets due to their ability to oxyfunctionalize diverse organic molecules of industrial and environmental value. To address the lack of scalable screening for UPO functions, we applied enzyme-design calculations to focus experiments. Starting from an AlphaFold2 model of *Mth*UPO, the automated FuncLib algorithm generated 50 diverse active-site multipoint designs—all of which were functional. Screening against nine diverse terpenes revealed large improvements and new oxyfunctionalization products, resulting in molecules of high pharmaceutical and industrial value. We rationalized observed specificity changes using AI-based docking and molecular dynamics simulations, providing molecular insights that could generalize to engineering other UPOs. Thus, computational design and modeling can dramatically accelerate the urgently needed green transition of the chemical industry.

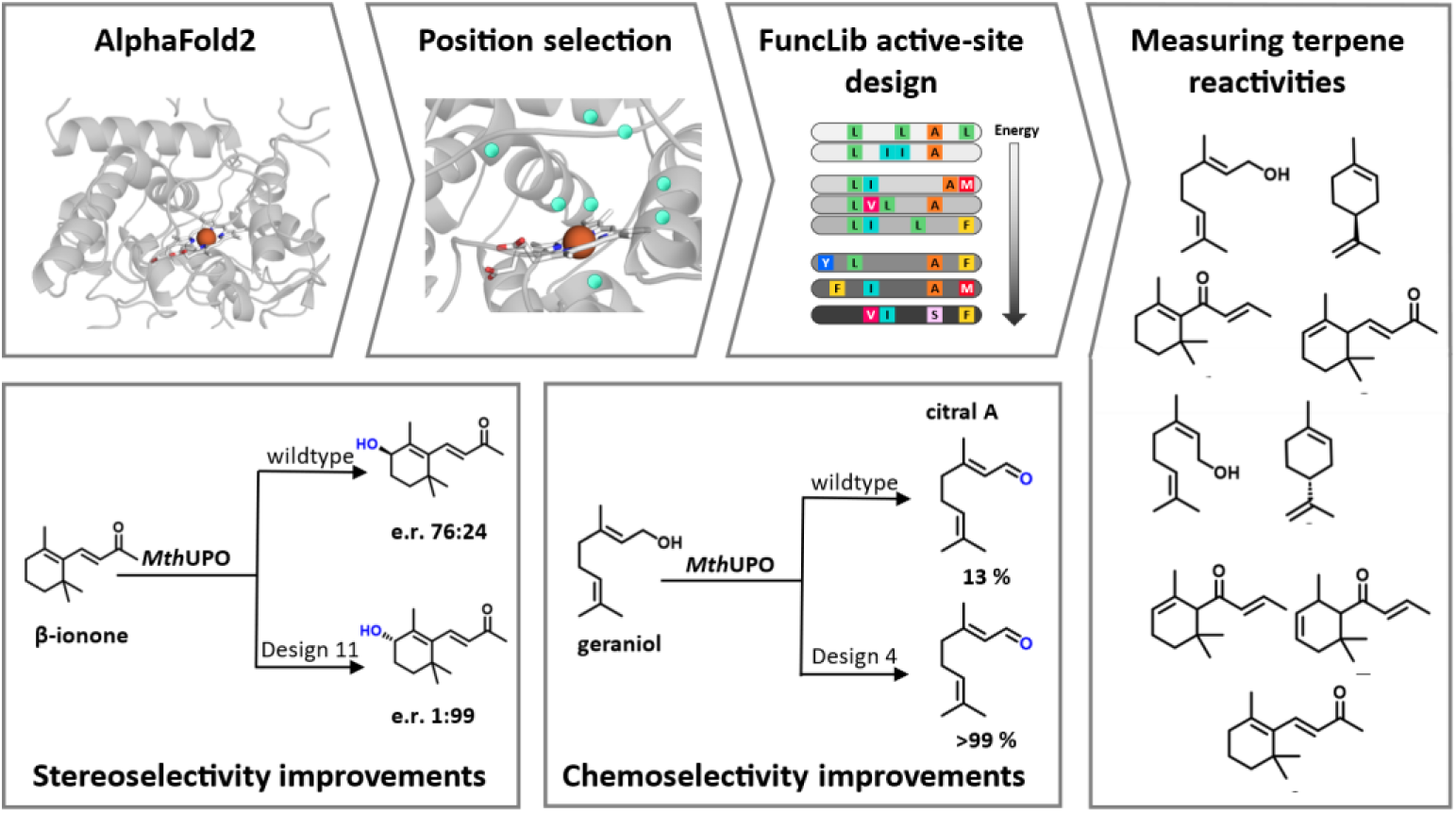

## Introduction

Selective oxyfunctionalization ranks among the most challenging and desirable reactions in synthetic chemistry. These reactions are often critical for selective C-H activation and alkene epoxidation of complex organic molecules.^1–4^ Since their discovery in 2004, fungal unspecific peroxygenases (UPOs) have attracted great interest for their ability to perform versatile oxyfunctionalization reactions on a broad scope of substrates.^5–7^ These enzymes have several advantages relative to other versatile oxidases, such as P450s^8,9^: they are typically stable,^10^ use pre-reduced hydrogen peroxide as co-substrate instead of molecular oxygen and expensive reductants such as NAD(P)H, exhibit a broad substrate scope^11^ and high turnover numbers (TON) up to 900,000^12^. UPOs can be engineered for overexpression in rapidly proliferating host organisms, such as yeast, through the implementation of protein^13^ and signal-peptide^14^ engineering and promoter-shuffling techniques.^15^ Recent studies yielded enhanced variants characterized by increased activity,^16^ augmented thermo-,^17^ pH-,^18^ and solvent-stabilities,^17^ and large shifts in chemo-, regio-, and stereoselectivities.^19–24^ These studies frequently employed directed evolution,^20,23,25,26^ a prominent protein-engineering approach that emulates the engineering prowess of natural evolution through iterative rounds of random or semi-rational mutation and selection of variants that exhibit desirable properties.^27^ But despite yielding exceptional results, directed evolution is labor intensive and time consuming. It is especially impractical when seeking enzymes that exhibit improvements across multiple substrates, as mutations that are favorable for one substrate rarely benefit others.^28^ Furthermore, measuring oxyfunctionalization products is not amenable to chromogenic or fluorogenic measurement, limiting options for medium to high throughput screening which is often essential to successful *in vitro* evolution campaigns. These limitations are currently addressed through small, “smart” libraries, using insights from molecular structures and mechanistic information, reducing the screening effort from thousands of variants to several hundred per round of evolution.^29–31^ As a pertinent example, we recently performed an engineering campaign starting from the UPO from *Myceliophthora thermophila* (*Mth*UPO) resulting in two UPOs that selectively produced (*S*)- and (*R*)-4-hydroxy-β-ionone, with enantiomeric ratios of 97:3 (*R*) and 0.3:99.7 (*S*), respectively.^23^ This study establishes a benchmark for rational engineering of a challenging UPO: we screened in total 2,500 variants in three rounds of evolution, 75 % of which displayed lower activity than the parental variant.

Here, we ask whether recent developments in computational protein design and engineering^32^ can address the limitations of rational engineering of enzyme active sites through a limited experimental effort.^33^ Such protein-design methods can be applied to crystallographic structures and can be generalized to AI-based model structures.^23,34,35^ For example, FuncLib is an automated method for designing diverse combinations of multipoint mutations within the active site of an enzyme.^35^ This method uses phylogenetic analysis and Rosetta atomistic design calculations to generate variants that exhibit large diversity in active-site geometry and electrostatics without impairing the stability, foldability and primary activity of the enzyme. The resulting designs may exhibit strong epistatic dependencies among the mutations^35–37^ that are rarely observed in natural and lab-evolved variants; yet these designs exhibit diverse activities suggesting that FuncLib may uncover sequences and activities that are difficult for evolutionary processes to reach.

We focus on oxyfunctionalization of small terpenes which constitute the largest class of secondary plant metabolites.^38^ Terpenes and their oxyfunctionalized derivatives, terpenoids, frequently showcase pharmacological activity^39^ and find applications in the flavor and fragrance industries.^40,41^ Chemo-, regio- and stereoselective oxygenation presents a challenge for chemical catalysis, given that many terpenes feature multiple sites where C-H hydroxylation and C=C epoxidation are chemically and energetically almost indistinguishable. Enzymes are naturally stereoselective and can position substrates to promote reactions at desired positions that are not kinetically favored or chemically activated; thus, accurate control of the UPO active-site pocket may enable oxyfunctionalization reactions that overcome the dictates of intrinsic chemical reactivity of the substrate.

Starting from an AlphaFold2 model of *Mth*UPO, we generated 50 *Mth*UPO FuncLib designs. All proved to be functionally secreted from yeast, allowing us to compare their influence on various substrates and to find large activity improvements and substantial shifts in chemo-, regio- and stereoselectivity through a limited experimental screening effort (**Fig. 1**). AI-based modeling and molecular dynamics simulations of substrate-enzyme pairs provided insights into some of the dramatic specificity changes that may extend to other UPOs.

**Figure 1.**
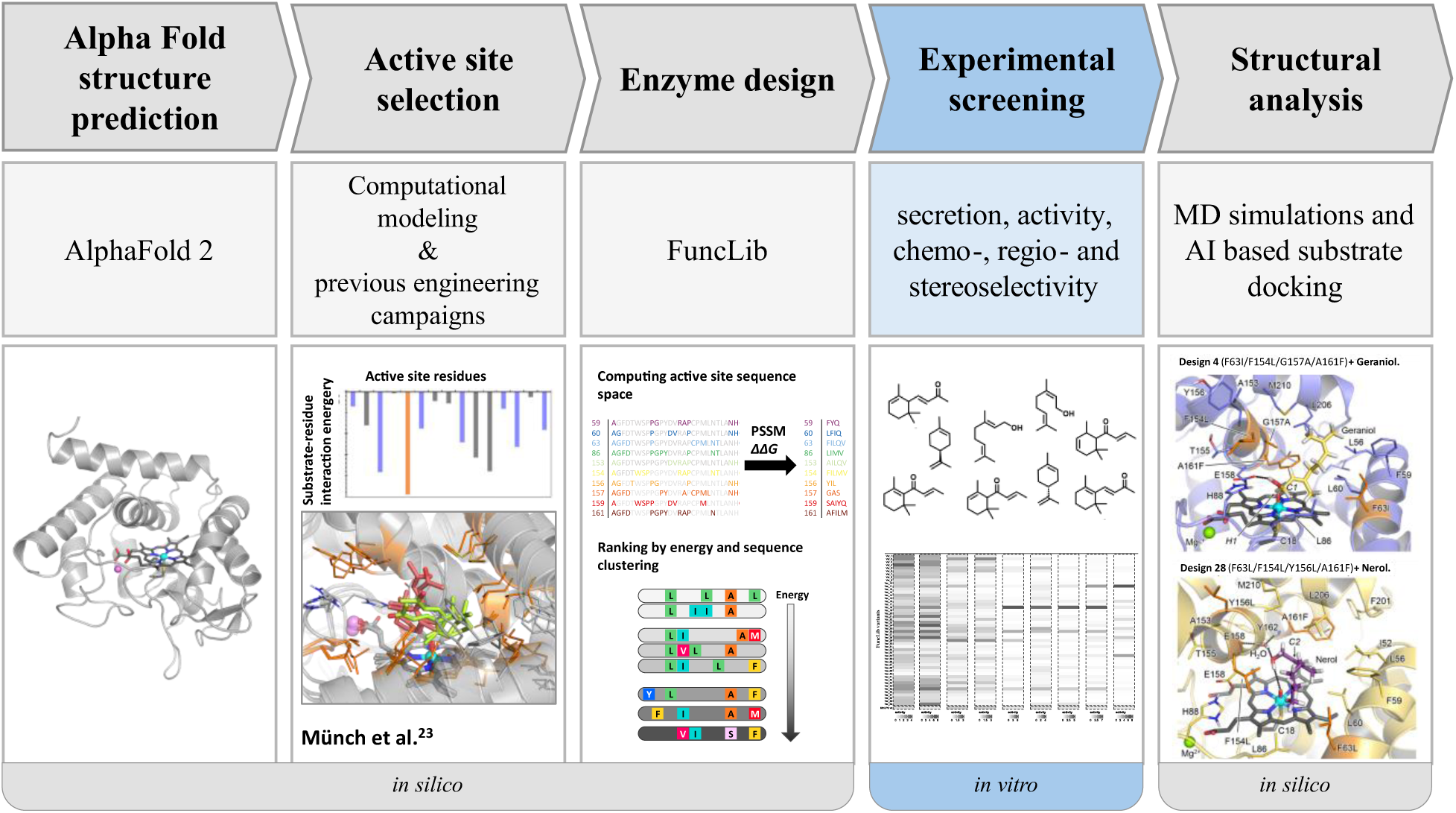
Overview of the different steps of the work protocol.

## Results

### Modelling and design calculations

Because the molecular structure of *Mth*UPO has not been determined experimentally, we started from an AlphaFold2 model (Uniprot entry: G2QID2) of the core enzymatic domain (amino acid positions 1-227).^42^ Because AlphaFold2 does not model cofactors, we compared the model to the structure of an artificial UPO (artUPO, PDB entry: 7ZNV), a homolog from *Marasmius rotula* (*Mro*UPO).^43^ Visual inspection verified that the side chain conformations of amino acids that are in direct contact with heme in other UPOs were aligned with those observed in experimentally determined UPOs.^23,43^ The proximal axial ligand Cys18 and the catalytic dyad Glu158/His88 were kept fixed in their modelled conformations in all design calculations to maintain the core catalytic activity. Active-site positions were chosen for design to alter the active-site cavity based on our previous work and modelling (**Table S1**).^23^ The designs were selected to encode 2-4 mutations and at least two mutations from one another. 50 lowest-energy FuncLib designs (**Table S2**) were selected for experimental screening (see section *FuncLib design* in the SI for further details). Among the 50 designs, some positions exhibited no mutation (Leu206 and Met210) or only low sequence diversity, reflecting the high sequence conservation and energetic sensitivity of active-site positions, whereas others showed radical mutations with position Gly157 mutated in 88 % of the designs.

### Designs generate chemically challenging oxyfunctionalized terpenes

We selected ten small cyclic and non-cyclic terpenes and terpenoids (**Scheme 1**) to assess the catalytic capabilities of the designed *Mth*UPO variants. Additionally, we chose three standard colorimetric substrates that serve as indicators for peroxygenase or peroxidase activity (ABTS (2,2’-azino-bis(3-ethylbenzothiazoline-6-sulfonic acid), DMP (2,6-dimethoxyphenol), and NBD (5-nitro-1,3-benzodioxole) and performed a splitGFP assay to determine the protein secretion levels independently of activity.^44^

**Scheme 1.**
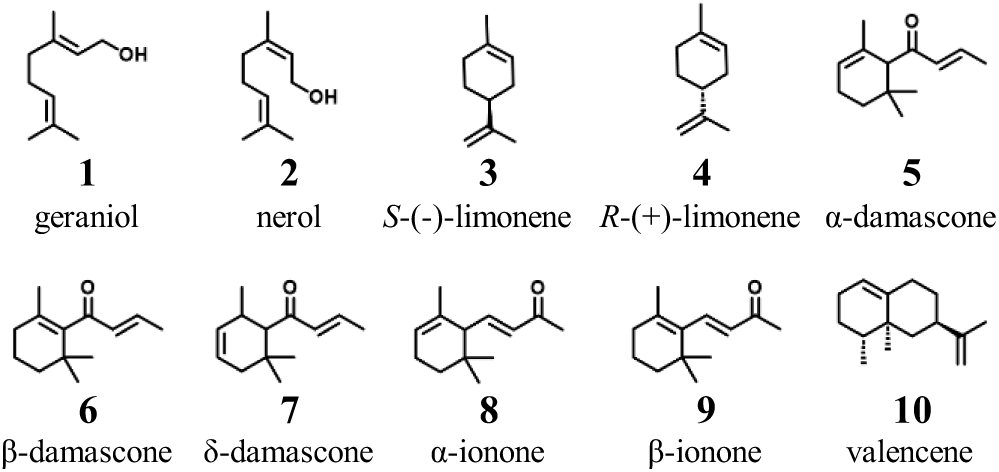
Terpene substrates investigated in this study.

Amongst the substrates, the oxyfunctionalized products of geraniol and nerol are of outstanding interest, as partial oxidation of their terminal alcohol to aldehyde produces citral A (geranial, **33**) and citral B (neral, **31**), which are commonly used aroma compounds in perfumery due to their strong lemon (citrus) scents.^45^ Moreover, citral is a key building block for the synthesis of vitamin A^46^ and shows anti-inflammatory, anti-tumor, and anti-bacterial activity.^47–49^ Limonene and its oxyfunctionalized products are widely used in the fragrance^50,51^ and polymer^52,53^ industry, and have important pharmacological effects^54–57^. Wild type *Mth*UPO, however, yields a diverse array of products from limonene (**Fig. S3**), including multiple oxyfunctionalizations, similar to the results observed with PaDa-I and artUPO.^43^ Additionally, the volatility of limonene renders upscaling reactions for product isolation and identification challenging.

Wild type *Mth*UPO is readily secreted in functional form from yeast, and remarkably, all 50 designs were successfully expressed and secreted. Secretion levels were between 0.6-fold and 1.9-fold of wild type levels (**Fig. 2**), indicating that active-site design using FuncLib preserves the expressibility of the parental enzyme. Furthermore, among the designed enzymes, we measured improvements in activity relative to the wild type for nine of the ten terpenes, with only valencene (**10**), the only substrate comprising two conjugated rings in our set, not showing detectable activity in the wild type or any of the designs. The greatest improvement was seen in designs 3 and 47 relative to ABTS (1,880-fold and 1,950-fold increase in absorbance related to product formation, respectively). Design 34 demonstrated a 200-fold activity improvement on DMP (**Fig. 2**). Both design 3 and 47 performed excellently as well for DMP, but underperformed with all other substrates, indicating that those designs improved the peroxidase rather than peroxygenase activity. By contrast, designs 22 and 25 showed twofold improvement on the peroxygenase model substrate NBD.

**Figure 2.**
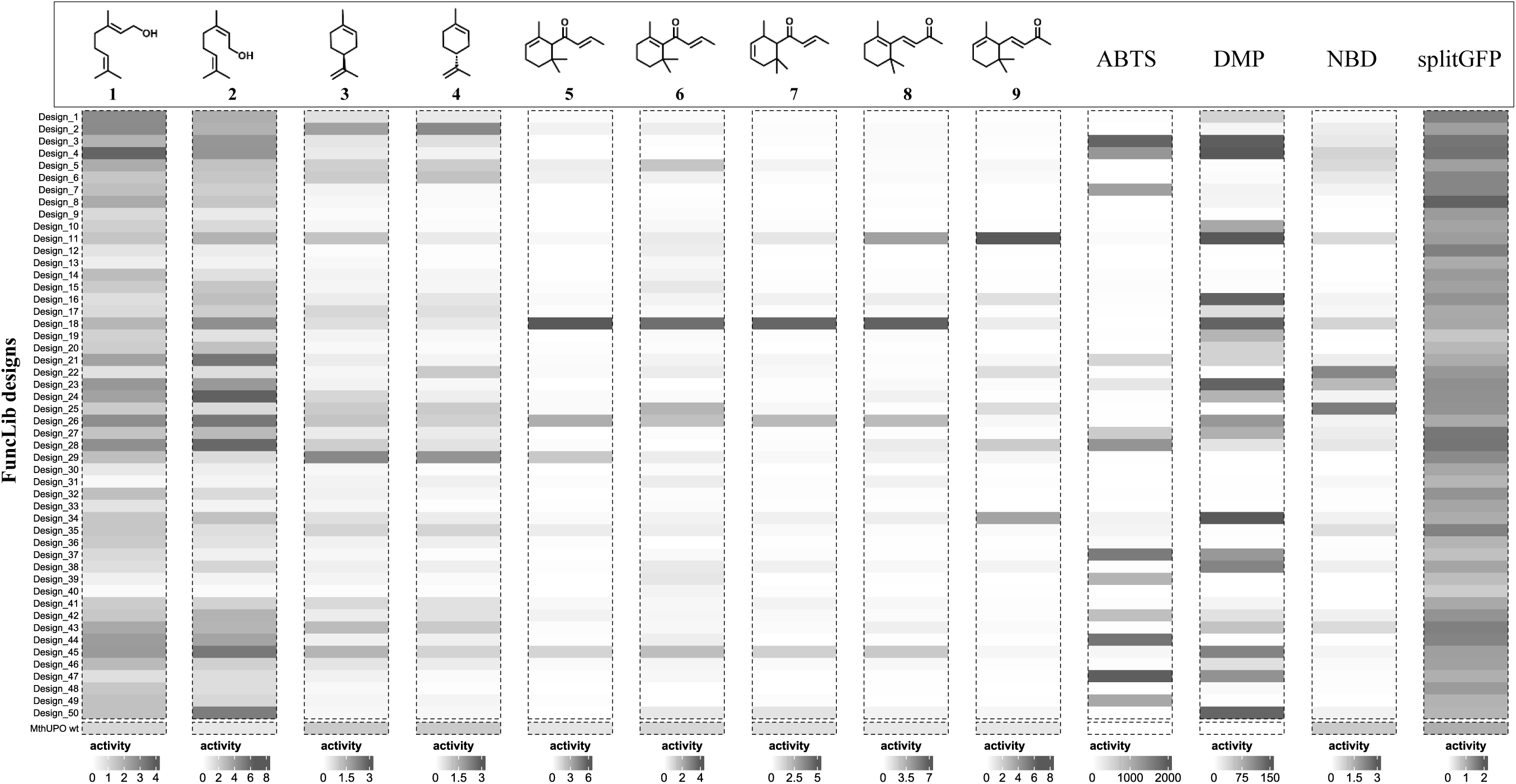
Total activity (sum of all products when multiple products were formed) relative to the activity of wild type *Mth*UPO. The activity of wild type *Mth*UPO is set to 1 separately for each substrate. All measurements were performed in a single replicate in microtiter plates. ABTS, DMP, NBD were analyzed photometrically, while product formation for all other substrates was determined by GC-MS.

All FuncLib designs showed at least baseline activity toward geraniol (**1**) and nerol (**2**), confirming that all designs were functionally secreted. The largest improvement toward geraniol was seen with design 4 exhibiting 3.8-fold increase compared to the wild type. The highest activity for nerol (**2**) is seen by design 24 with 6.2-fold improvement. The highest overall activity toward all damascones and α-ionone (**8**) is seen in design 18 with 3.1 to 6.7-fold increases. The highest activity on β-ionone (**9**) was revealed by design 11 with a sevenfold increase relative to the wild type. For limonene, the activity gain reached 1.7 and 2.2-fold with design 2 and 29, respectively, for (*R*)-(+)-limonene (**4**) and (*S*)-(-)-limonene (**3**). Thus, a variety of different designs exhibited excellent levels of oxyfunctionalization of different substrates.

### Novel products and outstanding improvements in regioselectivity

Aside from the increase in activity, we also observed dramatic shifts in regioselectivity among the designs (**Fig. 3**). Design 45 displays a notable shift in the formation of 3-hydroxy-β-damascone (**15**), which increased from 3 % for wild type *Mth*UPO to 46 %. This regioselectivity shift is remarkable as the aliphatic C3-position is substantially less activated compared to the allylic C4-position of the main product 4-hydroxy-β-damascone (**14**). Thus, the designs can substantially change the profile of the resulting products towards ones that are less favored for chemical reactivity.

**Figure 3.**
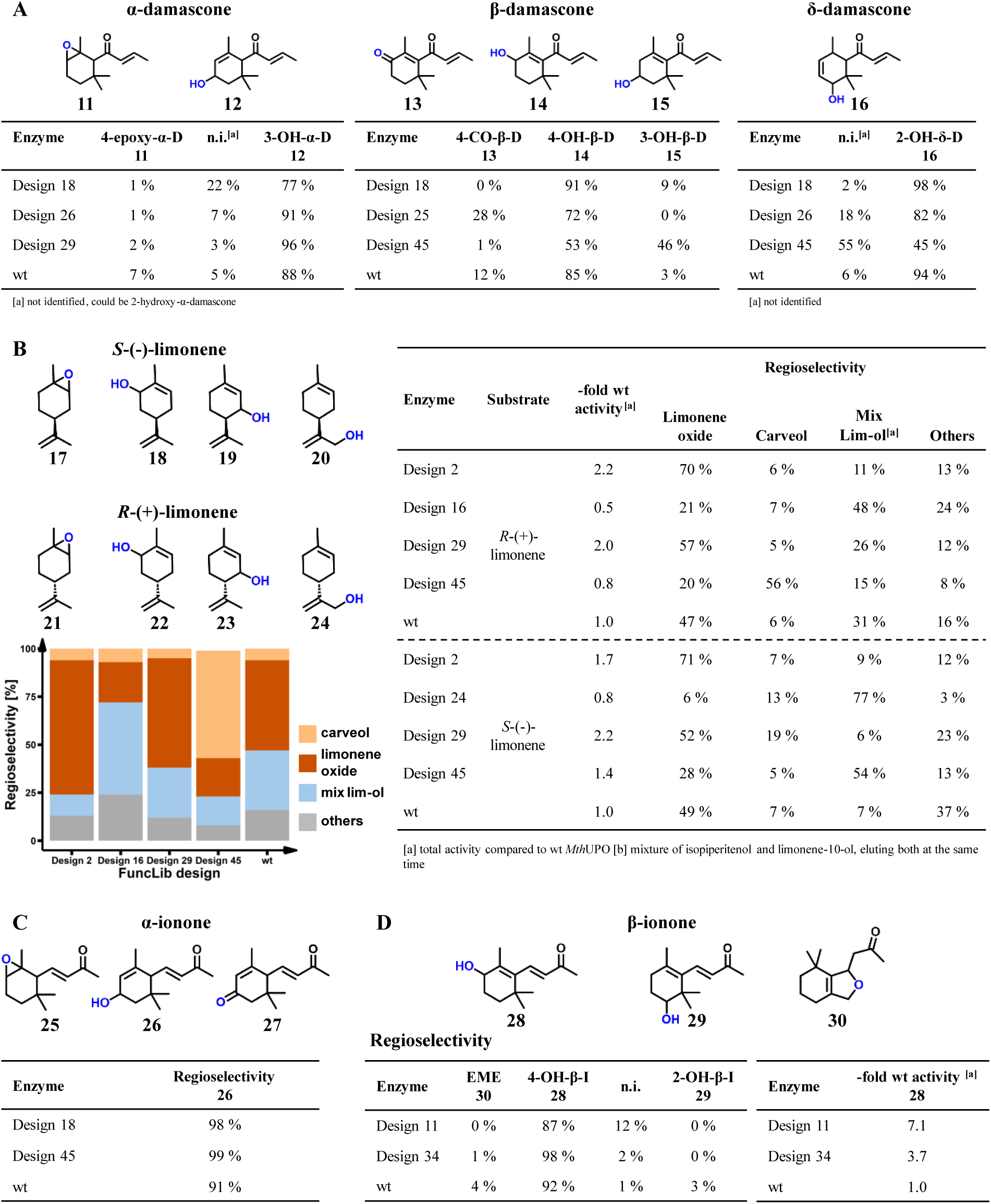
Relative abundance of selected products after conversion with different FuncLib designs. All data are analyzed by GC-MS. **A** Conversion of α-, β- and δ-damascone. **B** Conversion of (*S*)-limonene and (*R*)-limonene stacked bar chart displays the relative abundance of products after conversion of (*R*)-limonene leading to carveol (**22**) (orange), limonene oxide (**21**) (red), mix lim- ol (**23/24**) (blue) and others (grey). Isopiperitenol elutes on the GC-MS simultaneously with a second limonene alcohol (presumably limonene-10-ol), elution peaks are not separable, but mass spectra indicate the presence of two different moieties, the sum of both products is referred to as *mix lim-ol*. **C** Conversion of α-ionone. **D** Conversion of β-ionone reveals differing relative product abundance, and increased enzyme activity compared to wildtype *Mth*UPO.

We also saw shifts in chemo- and regioselectivity for limonene. For instance, wild type *Mth*UPO mainly generates the epoxide limonene oxide (**21**), whereas design 45 mainly produces (+)-carveol (**22**) (Fig. 5D) from (*R*)-(+)-limonene, alongside a 5.6-fold increase in turnover number (TON). This change in chemoselectivity is noteworthy as hydroxylation reactions are energetically less favored than epoxidation reactions. Moreover, this reaction provides access to a valuable fragrance and pharmacologically active compound.^54,55^ The main product of (*S*)-(-)-limonene with design 45 was identified as isopiperitenol (**19**), which could only be found in trace amounts after conversion by wild type *Mth*UPO. This activity is boosted 25-fold, reaching up to 310 TONs.

α- and β-ionone are mainly converted by *Mth*UPO to 3-hydroxy-α-ionone (**26**) and 4-hydroxy-β-ionone (**28**).^23^ With the FuncLib designs the main products remained **26** and **28**, respectively, but the regioselectivity increased strongly. For the formation of **26** it increased from 91 % (wild type *Mth*UPO) to 98 % (design 18) and 99 % (design 45), respectively. For **28** the improvement was from 92 % (wild type *Mth*UPO) to 98 % (design 34).

Overall, our experiments revealed large shifts in the main substrates, the formation of novel products, and an increase from satisfactory to outstanding regioselectivities, thus underscoring the efficacy of FuncLib in generating useful functional diversity. In addition, some FuncLib designs significantly enhanced reactivities and produced regioisomers that are energetically disfavored, demonstrating how changes to the active-site geometry and electrostatics can dramatically impact the outcome of the reactions (see below).

All measurements were performed in triplicates utilizing enzymes in supernatant from a microtiter plate expression.

### Large chemoselectivity shifts among FuncLib designs

FuncLib also leads to large shifts in chemoselectivity as demonstrated by geraniol (**1**) and nerol (**2**). Design 4 displays excellent selectivity (>99 %) for citral A (geranial, **33**, **Fig. 4**) and a 4.5-fold improvement in TON compared to wild type *Mth*UPO, which only showed 40 % chemoselectivity for converting the terminal alcohol group. The most pronounced chemoselectivity shift toward citral B (neral, **31**) was achieved with design 26 with 89 % relative abundance reaching 11,170 TONs, which corresponds to a 4.5-fold activity gain, but only a minor chemoselectivity shift compared to the 72 % of wild type *Mth*UPO. Further products are 2,3-epoxides, with a relative abundance of up to 58 % for 2,3-epoxy geraniol (**34**) with design 2 and 69 % for 2,3-epoxy nerol (**32**) with design 24. Some designs further exhibited overoxidation from nerol to neric acid, a reaction that was not observed with geraniol (**Fig. 4B**). The selectively oxidized aldehydes citral A and citral B are of great importance for industrial application, which renders design 4 with its outstanding chemoselectivity an exciting biocatalyst. Upscaling reactions with design 28 were successfully performed and yielded more than 150 mg of 2,3-epoxy nerol, which was subsequently used to determine TONs of both nerol and geraniol 2,3 oxides. Design 2 displayed a 1.8-fold increase in TON for **34** when using geraniol. Design 28 displayed a 14.3-fold boost for **32** compared to the wild type *Mth*UPO enzyme, resulting in 9,470 TONs for product **32 (Fig. 4C)**.

**Figure 4.**
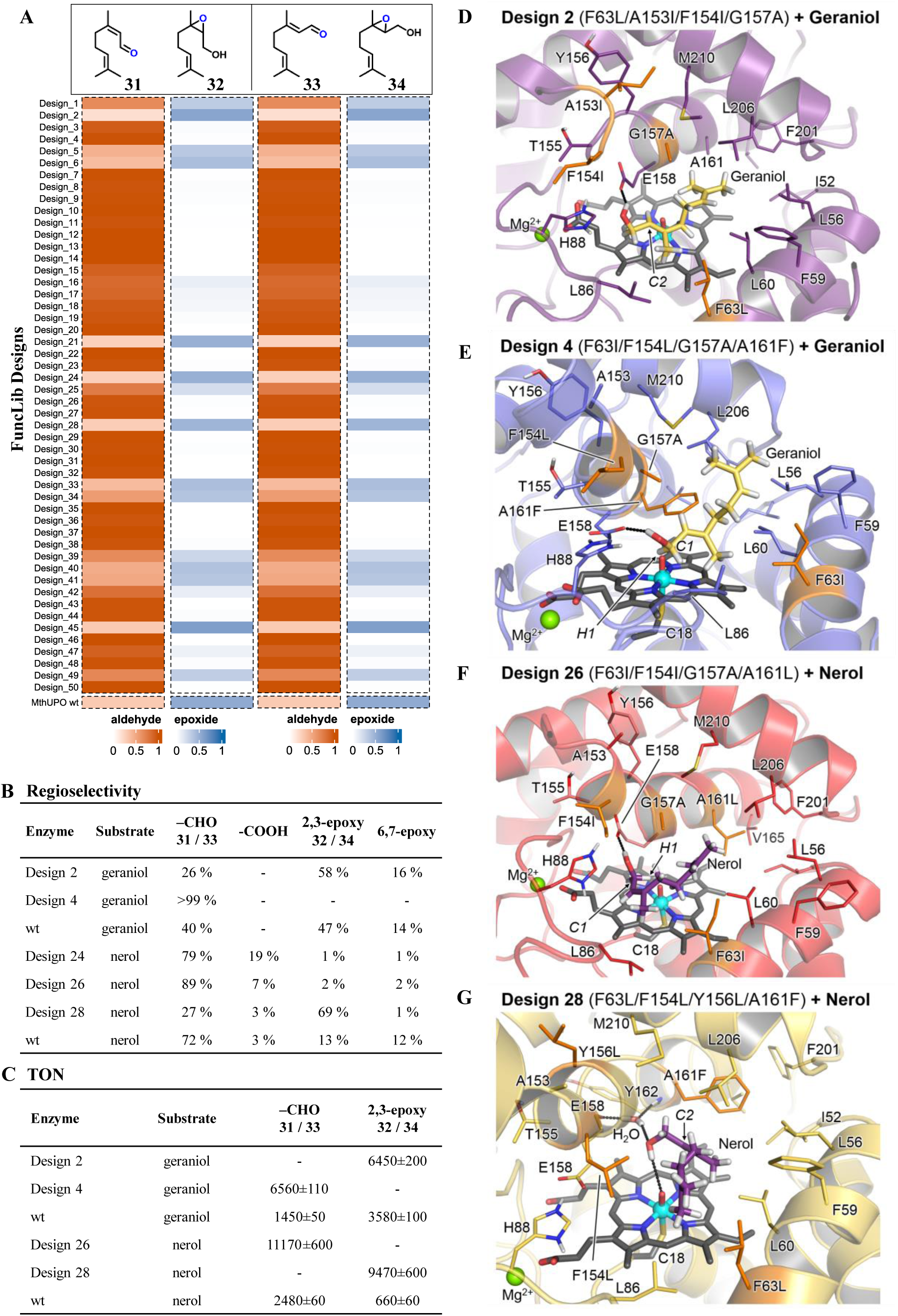
Oxyfunctionalization of geraniol (**1**) and nerol (**2**) with FuncLib designs of *Mth*UPO. **A** Proportion of 2,3-epoxide (**32/34**) (blue) and aldehyde (**31/33**) (orange) formation in the overall reaction. Measurements were performed in single replicates with enzyme from microtiter plate supernatant. **B** Regioselectivity of selected enzyme variants toward citral A (**33**)/ citral B (**31**), neric acid and 2,3-epoxy (**32/34**) and 6,7-epoxy formation measurements were performed in triplicates. **C** Turnover number of selected products, data are mean ± s.d. of measurements in triplicates. **D-G** Most representative structure of the binding mode of geraniol / nerol in different FuncLib deisngs as determined by clustering analysis.

### Molecular basis of chemoselectivity changes

To shed light on the molecular basis for the chemoselectivity changes observed on the non-cyclic terpene substrates, we conducted computational modeling based on DFT calculations using truncated models and molecular dynamics (MD) simulations. For both geraniol and nerol, DFT calculations in the absence of enzyme show that allylic hydroxylation on the terminal carbon atom (C1) and epoxidation at the 2,3 double bond are the energetically most favored oxyfunctionalizations, with the terminal allylic C-H activation (via Hydrogen Atom Transfer, HAT) slightly preferred (**Figs. S6** and **S7** in the SI). This is in line with the major products detected experimentally for these substrates with all variants.

We chose to conduct Molecular Dynamics (MD) simulations (see SI for details) to analyze the large selectivity shifts seen in design 4 vs. design 2 for geraniol oxidation and design 28 vs. 26 for nerol. According to these simulations, geraniol establishes a persistent polar interaction with the Glu158 sidechain of design 4, thus positioning both C1-H and C2-C3 bonds near the catalytically active oxo-ferryl cation radical complex species (Compound I, see **Fig. 4E** and **Fig. S8** in the SI). The presence of the bulky Phe161 (mutated from the wild type Ala) side chain prevents the substrate from approaching the catalytic species in alternative near-attack conformations. Considering that terminal allylic hydroxylation is energetically slightly preferred, the citral A product is preferentially formed from this binding mode. On the other hand, design 2 led mainly to 2,3-epoxy geraniol with high activity and moderate chemoselectivity (1.8-fold increase and 58% selectivity). The models show that design 2 has a less dense active site and the C2-C3 double bond of geraniol may approach the catalytic iron-oxo group. At the same time, C1-H bonds are not geometrically well oriented for effective HAT (**Fig. 4D** and **Fig. S9** in the SI), leading to effective epoxidation by subtle substrate repositioning in the active site as compared to design 4.

A similar scenario is found for nerol and designs 26 and 28. In design 26, nerol is preferentially bound by placing both the allylic terminal position and the C2-C3 double bond geometrically preorganized for oxyfunctionalization (**Fig. 4F** and **Fig. S10** in the SI). Nevertheless, mutations included in design 28, particularly Tyr156Leu, induce structural changes in the active-site cavity that favor the positioning of a water molecule that persistently interacts with the Glu158 and Tyr162 backbones. These structural changes, stemming from cooperation among several mutations, reposition nerol in the active site placing the C2-C3 double bond in a near-attack conformation for epoxidation by Compound I while geometrically disfavoring terminal C-H activation (**Fig. 4G** and **Fig. S11** in the SI).

Recently developed AI-based structure predictors, such as AlphaFold3,^58^ can accurately predict biomolecular interactions between proteins, cofactors, and ligands. We asked whether the open-access predictor Chai-1^59^ could be used to shed light on the promiscuity and specificity of some of the designed enzymes modelling selected design/substrate pairs (**Fig. 5**). We focused on design 18, which exhibits high activity toward substrates 5-8 (**Fig. 2**), and generated *ab initio* models for the design, substrates, heme and Compound I oxygen. The models show that the substrates may all orient similarly, with the Compound I oxygen atom placed close to the specific carbon atom that undergoes oxidation (**Fig. 5A**; **Fig S14**) In addition, we asked why design 18 oxidized substrate 7 much more efficiently than design 11. Modeling suggests that three mutated positions close to the Compound I oxygen determine this specificity profile: in design 18, Phe60, Ile154 and Leu161 provide sufficient room for the ring to come close to the reactive oxygen, whereas Leu60, Leu154 and Phe161 in design 11 prevent this close approach (**Fig. 5B**).

**Figure 5.**
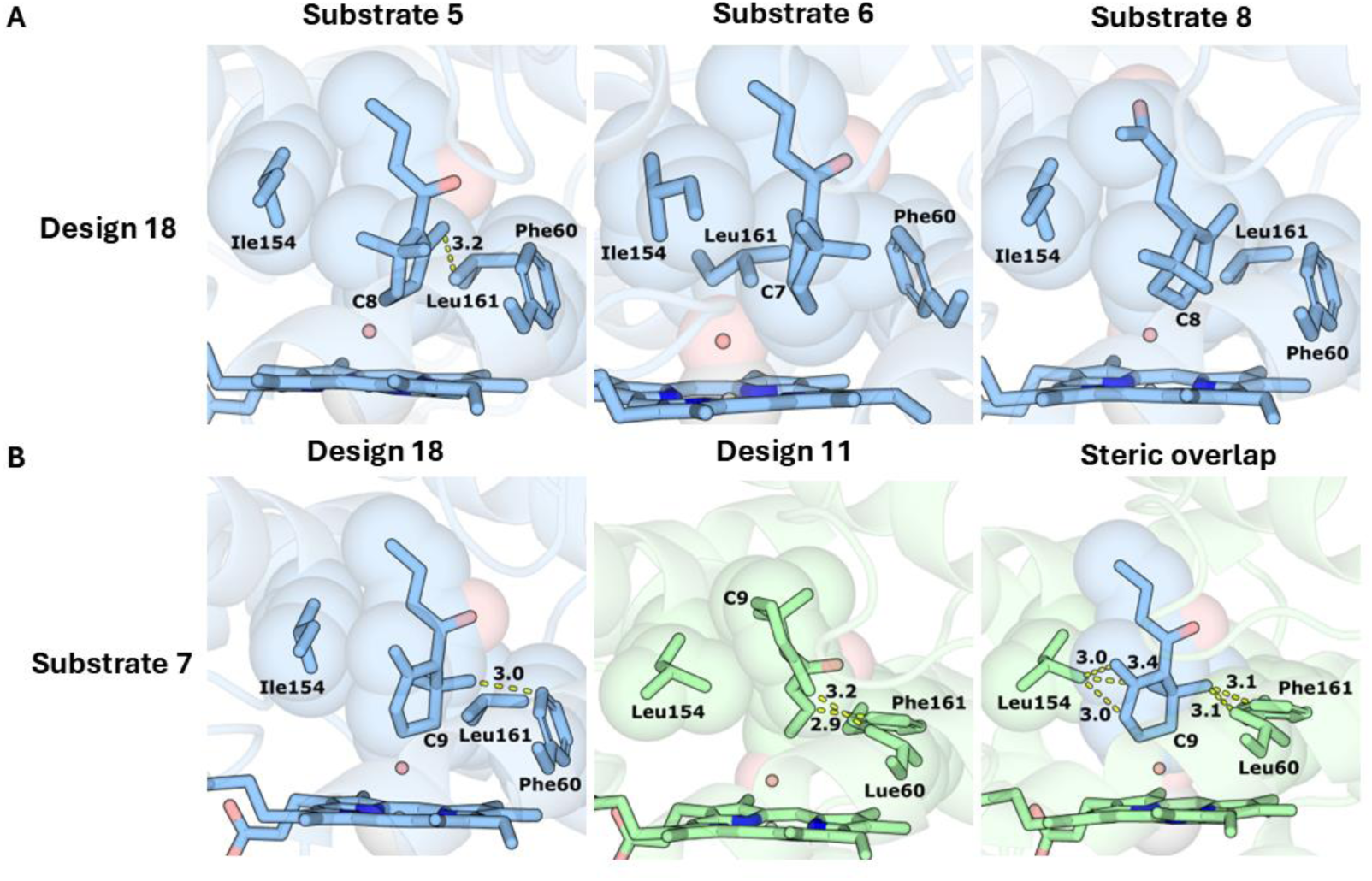
AI-based docking provides a molecular basis for the observed reactivities in design/substrate pairs. (A) Design 18, substrates 5, 6, and 8, and Compound I were modeled using Chai-1, generating up to 50 models for each. For each pair, we selected a model that formed a near-attack conformation (NAC) defined as exhibiting distance < 4.5 Å between the oxidized carbon and the Compound I oxygen. (B) Substrate 7 exhibits much higher activity with design 18 than with design 11. In both cases, 10 models were generated with Chai-1. *(left)* The modeled interaction places the oxidized C9 atom of substrate 7 in a NAC. (*center*) Design 11 does not accommodate the NAC with this substrate, and (*right*) superimposing the substrate conformation modeled in design 18 in the active-site pocket of design 11 reveals significant steric overlap with designed amino acid residues that are close to the heme co-factor. Carbons in blue or green; Compound I oxygen in red sphere; heme in sticks; steric overlap and interatomic distances (Å) indicated with dashed yellow lines.

**Figure 6.**
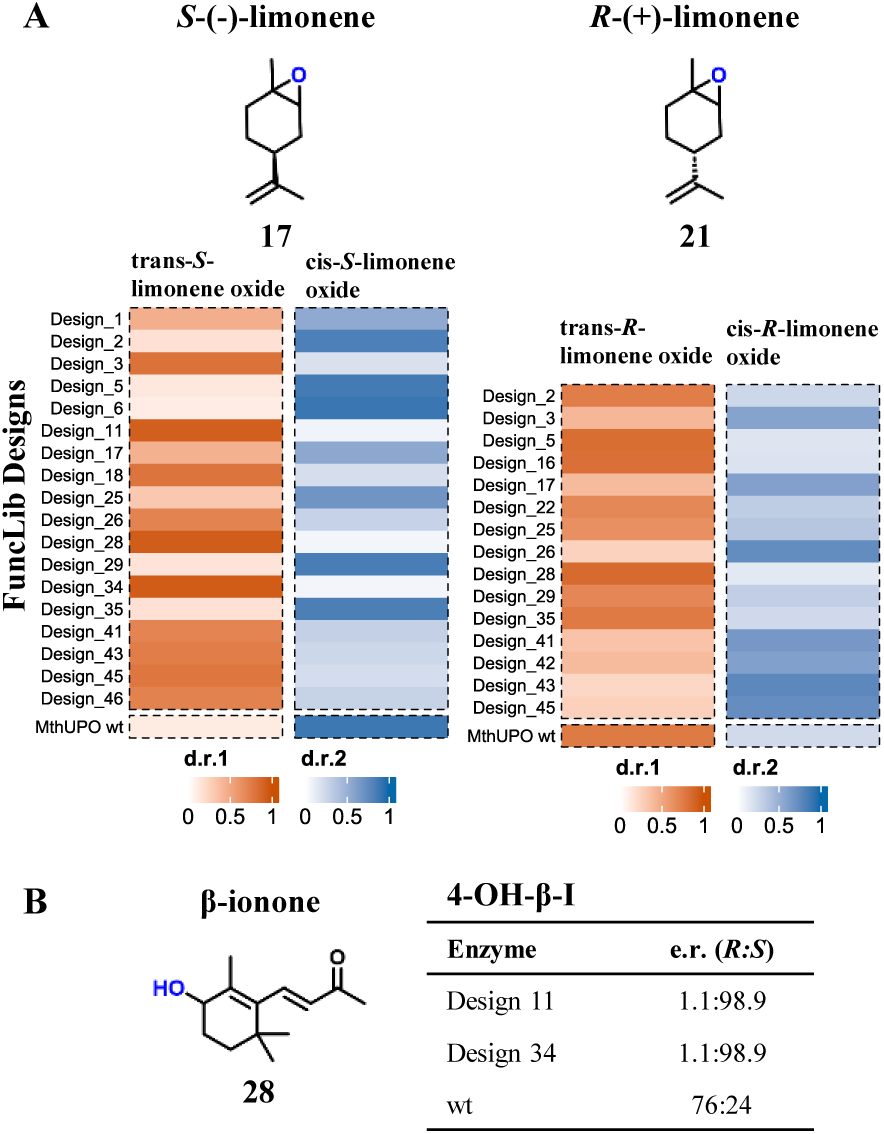
Influence of FuncLib designs on stereoselectivity. **A** Shifts in the diastereomeric ratio of *trans*-limonene oxide (orange) and *cis*-limonene oxide (blue). Measurements were performed as singe replicate with enzyme from supernatant in microtiter plate scale. **B** Shifts in enantiomeric ratio of 4-hydroxy-β-ionone (**28**) giving access to *S*-hydroxy-β-ionone with Design 11 and Design 34. Measurements are performed as single replicates.

### FuncLib designs impact stereoselectivity of limonene oxide and 4-hydroxy-β-ionone

The major influence of the FuncLib library on stereoselectivity is shown by shifts in the diastereomeric ratio of 1,2-epoxy limonene (**Fig. 5 A**) ranging from 86:14 (design 28) for *trans*-(*R*)-limonene oxide to 23:77 (design 43) for *cis*-(*R*)-limonene oxide and 94:6 (design 28 and 34) for *trans*-(*S*)-limonene oxide to 10:90 (design 6) for *cis*-(*S*)-limonene oxide. We further see outstanding changes in the enantioselectivity of 4-hydroxy-β-ionone (**28**) from an enantiomeric ratio of 76:24 for wild type *Mth*UPO to 1:99 for both design 11 and 34 producing (*S*)-4-hydroxy-β-ionone (**Fig. 5B**). Lack of enantioselectivity of the FuncLib designs towards the (*R*)-4-hydroxyl-β-ionone product is not surprising as the key position for controlling *R*-selectivity in β-ionone hydroxylation (Leu206)^23^ was not diversified in the FuncLib library (**Table S2**). These shifts confirm that FuncLib not only influences regio- and chemoselectivity, but can further exert major influences on stereoselectivity in accordance with previous work on small aromatic substrates.^36^

## Discussion

In both industrial and academic contexts, the ability to produce a small set of highly active enzymes with diverse selectivities is extremely valuable to determine the feasibility of desired reactions. Previously, such diversity was generated, including in UPOs, by using natural or engineered enzymes from a variety of sources. We demonstrated here that substantial diversity can also be provided by the FuncLib algorithm starting from a single enzyme that exhibits favorable stability and activity profiles.

All FuncLib designs we tested experimentally were functional and exhibited large and potentially useful changes in activity and selectivity profiles. This is remarkable, especially given that the designs were based on an AlphaFold model and that the UPO family is challenging for protein engineering due to difficulties in heterologous expression, a limited number of solved crystal structures and its high glycosylation. For example, in a previous engineering effort in our lab we screened 2,500 *Mth*UPO active-site variants, of which 75 % displayed lower activity than the parental variant. By contrast, screening 50 designed variants in the current study led to substantial improvements in activity, chemo-, regio- and stereoselectivity such as >99 % chemo- and regioselectivity for the fragrant and pharmacological active citral A (starting from 40 % for wild type *Mth*UPO), more than 1,900-fold activity increase for ABTS, variants with at least doubled activity for all substrates, 98 % regioselectivity and an inversion of enantioselectivity to an e.r. of 1:99 (starting from 76:24) for (*S*)-4-hydroxy-β-ionone, increased regioselectivity for the formation of the energetically less favored 3-hydroxy-β-damascone (from 3 % to 46 %) as well as novel products and significant regioselectivity shifts with limonene. While achieving excellent selectivity may require further improvements in design methodology, FuncLib can identify substantially enhanced starting sequences for subsequent engineering campaigns, including using traditional engineering approaches, presenting a fascinating opportunity to accelerate enzyme engineering efforts towards new oxyfunctionalization selectivities.

In addition to high predictive value, the computational chemistry modelling retrospectively suggested mechanistic explanations and molecular bases for the high reactivities observed in several design variants. These calculations are highly sensitive to structural details and crucial for describing critical active site remodeling and substrate positioning, although they require significant computational effort. We are encouraged that the high reliability of AlphaFold and FuncLib calculations enables such sophisticated simulations yielding valuable insights into the structural rearrangements that enable efficient catalysis. Such insights may be useful in the design of efficient and selective enzymes in other UPO targeting different substrates. The high conservation of UPO structures around the heme binding site suggests that mutations in these positions may impact substrate specificity in other members of this family. We also found that the recent AI-based biomolecular modeling tools^58^ produce model structures that may help rationalize the observed specificity profiles. This is an encouraging sign that in the near future we may be able to focus design calculations on specific desired substrates using a combination of atomistic design and AI-based substrate docking calculations.

Notwithstanding the high success rate of the FuncLib design library, for some substrates, we found limited improvement in catalytic efficiency. We note that FuncLib usually designs thousands of active-site variants, but cost and time considerations preclude screening such large sets. We recently demonstrated an economical approach to design large combinatorial libraries for screening (htFuncLib).^60^ We envision that such an approach, in combination with mechanistic insights from computational modeling, may enable finding many more high-efficiency and specificity designs, including against substrates for which we did not find high-efficiency designs. Recent progress in AI-based modeling and evolution-guided atomistic design is providing unanticipated opportunities to address protein-engineering challenges at the forefront of biocatalysis that have frustrated conventional *in vitro* evolution approaches. The new approaches could be further refined by considering additional mechanistic insights from atomistic models obtained from computational chemistry calculations. In this study, we demonstrated that this combination effectively samples the functional space of an enzyme active site toward diverse regio- and enantioselectivity outcomes while accounting for high activities. Using this approach, in principle, any natural enzyme can be engineered quickly and effectively to tune its activity profile for basic or applied needs. The implications for green and medicinal chemistry are enormous.

## AUTHOR INFORMATION

### Author Contributions

Conceptualization, J.M., M.J.W, S.J.F. and M.G.B.; Methodology, J.M, S.J.F., J.S.; Validation, J.M., J.S., S.J.F. and M.G.B.; Formal Analysis, J.M., J.S., OGC; Investigation, J.M., OGC and J.S.; Resources, M.J.W, M.G.B. and S.J.F; Data Curation, J.M. and J.S.; Writing – Original Draft, J.M and J.S.; Writing – Review & Editing, J.M., J.S., S.J.F., M.G.B. and M.J.W; Visualization, J.M., OGC and J.S.; Supervision, S.J.F., M.G.B., M.J.W.; Project Administration, J.M.;

### Declaration of generative AI and AI-assisted technologies in the writing process

During the preparation of this work the authors used ChatGPT in order to improve readability and language of the manuscript. After using this tool, the authors reviewed and edited the content as needed and take full responsibility for the content of the published article.

### ASSOCIATED CONTENT

**The following files are available free of charge.**

**Supporting Information (PDF)**

• Material and methods; gene and protein sequence; table containing mutations for all FuncLib designs; GC-MS temperature programs; utilized linker and tags; structural characterization data for selected substrates and variants.

### Data and code availability

Raw data for all activities found in the screening process and the sequences of the 50 FuncLib designs can be found on the following dataverse server:

Muench, Judith, 2024, “UPO FuncLib Paper”, https://doi.org/10.7910/DVN/ZPKEAI, Harvard Dataverse, DRAFT VERSION

## ACKNOWLEDGMENT

J.M. thanks the Friedrich-Naumann-Stiftung für die Freiheit for a PhD scholarship. M.J.W. and J.M. thank the Bundesministerium für Bildung und Forschung (Maßgeschneiderte Inhaltsstoffe 2, 031B0834A) and the German Research Foundation (DFG, project ID 43649874, TP A05, RTG 2670) for generous funding. M.G.-B. thanks the Spanish MICINN (Ministerio de Ciencia e Innovación) for projects PID2022-141676NB-I00 and TED2021-130173B-C42, and the Ramón y Cajal RYC 2020-028628-I fellowship. Part of the computational resources used were funded by FEDER (Fondo Europeo de Desarrollo Regional) and Spanish MINECO (Ministerio de Economía, Comercio y Empresa) through projects CTQ2014-54306-P, CTQ2015-69363-P, and MPCUdG2016/096, as well as by the European Research Council (ERC) under the European Union’s ERC-StG-2015 (Grant agreement No. 679001).

## ABBREVIATIONS

HAT: hydrogen atom transfer
Mix-10-ol: mixture of limonene-10-ol and isopiperitenol
TON: turnover number
UPO: unspecific peroxygenase.

