## Supplemental Information for "A computationally designed panel of diverse and selective peroxygenases for terpene oxyfunctionalization"

**I Material and Methods**

**Chemicals.** Solvents were used as provided without further purification from Carl Roth (Karlsruhe, DE) as GC ultragrade. The commercially available compounds were also used without further purification from the following suppliers:

Sigma-Aldrich, St. Louis, US: ethanol (absolute), (*S*)-(-)-perillaldehyde, *trans*-(+)-limonene, (*S*)-(-)-limonene (>96 %), (-)-carveol (mixture of isomeres) (>97 %), δ-damascone (>90 %), (-)-limonene oxide (mixture of *cis* and *trans*) (99 %), β-damascone (≥90 %), (+)-valencene (>70 %), β-ionone (96 %), hydrogen peroxide solution (30 % (*w/w*) in H_2_O, naphthalene (99 %), NBD (98 %), ABTS, DMP

Carl Roth, Karlsruhe, DE: acetone (Rotisolv ≥99.9 % GC Ultra Grade), acetic acid ethyl ester (Rotisolv ≥99.9 % GC Ultra Grade),

Fluka, Buchs, CH: (-)-Carvone (>99 %), (-)-perillyl alcohol

TCI chemicals, Tokyo, Japan: 4-octanone (>98 %), citral (*cis* and *trans* mixture) (>96 %), (+)-limonene (>95 %), α-ionone (>90 %), nerol (>98 %), geraniol (>96 %)

For cultivation of *S. cerevisiae* cells d-Galactose, Peptone and Synthetic Complete Mixture (Kaiser) Drop-Out (-URA) were purchased from Formedium (Hunstanton, GB). Yeast nitrogen base (without amino acids) and Yeast extract were purchased from Carl Roth (Karlsruhe, DE). BsaI was purchased from New England Biolabs (Ipswich, US). BbsI and FastDigest AscI were purchased from ThermoFisherScientific (Waltham, US) and T4 DNA Ligase from Promega (Madison, US).

**Bacterial and Yeast Strains.** For all cloning purposes and plasmid propagation *E. coli* DH10B cells (ThermoFisherScientific, Waltham, US) were utilized. All work regarding *S. cerevisiae* was performed utilizing the INV*Sc*1 strain (ThermoFisher Scientific, Waltham, US).

**Oligonucleotides and gene parts.** All oligonucleotides were purchased in the lowest purification grade “desalted” and minimal quantity at Eurofins Genomics (Ebersberg, DE). The genes of the *Mth*UPO FuncLib library were purchased as gene parts from Twist Bioscience (San Francisco, US).

**AlphaFold2 model.** The AlphaFold2 model of *Mth*UPO was extracted from the AlphaFold Protein Structure Database (UniProt entry G2QID2). Source: https://alphafold.ebi.ac.uk/entry/G2QID2 (16.02.2023)

**FuncLib design.** We used a C-terminally truncated (-18 residues) AlphaFold2 model of *Mth*UPO with its natural signal peptide automatically refined by the FuncLib algorithm (using Rosetta) as protein structure. The loop at the original C‑terminus entered the substrate channel and active side during the simulations, impeding residue variation wherefore we used the truncated model for further calculations. We subsequently defined positions C18 (axial ligand) as well as E158 and H88 (catalytic cascade) as essential amino acids which must not be subject to any variation. The number of mutations per design was set to two to four with at least two variations between each design. We did not apply an algorithm like PROSS^1^ for introducing stabilizing mutations into the enzyme before running the FuncLib campaign, even though introducing several mutations in the active side may compromise enzyme stability. In our previous work with *Mth*UPO we did not encounter stability issues after introducing mutations. Further prior work on *Aae*UPO variant PADA-I showed, that an aggressive FuncLib campaign on unspecific peroxygenase can be tolerated without introducing further stabilizing mutations beforehand.^2^ We selected positions F59, L60, F63, L86, A153, F154, Y156, G157, S159, A161, L206, and M210 to be diversified. Charged amino acids (Arg, His, Lys, Glu, Asp) were excluded from the allowed sequence space to keep the hydrophobic character of the active site. Thr was also excluded as including all mutations to the initial sequence space file, that were found in previous projects (e.g. F59Q), exceeded the maximum permitted number of variants. The highly polar amino acid Thr was not considered to be of most interest for the project compared to other, more hydrophobic amino acids. Initial FuncLib runs showed sterically overlap of mutations containing aromatic identities at position L86, mutations larger than Ala and Ser at position G157 might obstruct access to the heme and the mutation A161F appeared to clash with the presumed heme site, therefore such mutations were prohibited. In the future the usage of AlphaFill or AlphaFold3 to include heme into the active site could prevent the introduction of mutations such as the forementioned. Finaly, the following sequence space was explored: **Table S1**.

**Expression plasmids.** Expression plasmid for *Mth*UPO expression in *S. cerevisiae* were constructed as previously described – all parts are available at AddGene.^3^ The gene parts of the FuncLib library were first cloned as individual level 0 standard modules into the universal Level 0 acceptor plasmid (pAGM9121) and afterward released upon BsaI restriction digest. Golden Gate reactions were performed to combine the *Mth*UPO FuncLib genes with the *Sce*-α.galactosidase signal peptide and a TwinStrep-GFP11-Tag for purification and secretion detection in a level 1 expression plasmid (pAGT572).

**Microtiter plate cultivation of *S. cerevisiae.*** The enzyme production in *S. cerevisiae* in microtiter plates was performed as described previously.^3^

**Shake flask cultivation of *S. cerevisiae*.** The enzyme production in *S. cerevisiae* in shake flasks was performed as described previously.^3^

**Supernatant ultrafiltration and protein purification.** The supernatant was concentrated approx. 20-fold by means of ultrafiltration as described before.^3^

**Heme CO complex measurements.** Concentration determination of enzymes in concentrated supernatant were performed as described before.^3^

**Colorimetric screening assays.** Four colorimetric assays were performed: ABTS, DMP, NBD and splitGFP-assay. All assay were performed as described before.^4^ For NBD, ABTS and DMP, the absorbance difference between five min (t1) and 0 min (t0) was evaluated. The DMP assay was performed with enzyme secreted in medium without additional heme to avoid background reaction.

**Bioconversion in microtiter plate.** All reactions with non-colorimetric substrates were initially performed in microtiter plate. 100 μL supernanat derived from enzyme expression and secretion in *S. cerevisiae* in a microtiter plate were transferred to a 96-deep well plate (CR1496, EnzyScreen, Heemstede, NL). 400 μL of a reaction mastermix were added to achieve final concentrations of 1 mM substrate, 1 mM H_2_O_2_, 5 % acetone and 100 mM Kpi (pH 7.0). Reactions were performed for 1 h atjh 30 °C under continuous shaking at 300 rpm. The extraction was accomplished through the addition of 500 μL EtOAc (GC Ultra Grade) containing 0.25 mM of an internal standard (**Table S3**) and further shaking for 30 min at 25 °C and 300 rpm. Microtiter plates were centrifuged to separate the phases (3000 rpm, 10 min) and 300 μL of the organic phase were transferred to glass coated microtiter plate utilizing the Platemaster (Gilson, Middelton, US) for subsequent GC-MS analysis.

**Bioconversion in single vials.** All bioconversions for regioselectivity determination were performed in tripliactes in single vials with direct addition of H_2_O_2_. 250 nM enzyme supernantant, derived from enzyme expression and secretion in *S. cerevisiae* in a shake flask after ultrafiltration and concentration determination were transferred to a glass vial. A reaction mastermix was added to achieve final concentrations of 1 mM substrate, 5 % acetone, 1 mM H_2_O_2_ and 100 mM Kpi (pH 7.0) to give a total volume of 500 μL. Reactions were performed for 1 h atjh 30 °C under continuous shaking. The extraction was accomplished through the addition of 500 μL EtOAc (GC Ultra Grade) containing 0.25 mM of an internal standard (**Table S3**), the organic phase was transferred to a new glass vial for subsequent GC-MS analysis.

**Bioconversion in single vials with syringe pump.** All bioconversion for TON determination were performed in triplicates in single vials utilizing a syringe pump system. 250 nM enzyme supernantant, derived from enzyme expression and secretion in *S. cerevisiae* in a shake flask after ultrafiltration and concentration determination were transferred to a glass vial. A reaction mastermix was added to achieve final concentrations of 1 mM substrate, 5 % acetone and 100 mM Kpi (pH 7.0) to give a total volume of 400 μL. 100 μL H_2_O_2_ (stock solution 5 mM, final concentration 1 mM) were added over the period of the reaction via a syringe pump. Reactions were performed for 1 h atjh 30 °C under continuous shaking. The extraction was accomplished through the addition of 500 μL EtOAc (GC Ultra Grade) containing 0.25 mM of an internal standard (**Table S3**), the organic phase was transferred to a new glass vial for subsequent GC-MS analysis.

**GC-MS analysis.** All GC-MS measurements were performed on a Shimadzu GCMS-QP2010 Ultra (Shimadzu, Kyoto, JP) with helium as carrier gas. The detector voltage of the secondary electron multiplier was adjusted in relation to the tuning results with perfluorotributylamine. The GC−MS parameters were controlled with GCMS Real Time Analysis, and for data evaluation, GCMS Postrun Analysis (GCMSsolution Version 4.45, Shimadzu, Kyoto, JP) was used.

**Non-chiral gas chromatography–mass spectrometry (GC–MS).** Measurements were performed on a SH-Rxi-5Sil MS column (30 m x 0.25 mm, 0.25 µm film, Shimadzu, Kyoto, JP). 1 µl of each sample was injected with a split ratio of 1:20 (inlet temperature 200°C). The utilized temperature program can be found in **Table S3**. Ionization was obtained by electron impact with a voltage of 70 V, and the temperature of the ion source was 280 °C. Calibration and quantification were implemented in scan mode. All initial screening measurements were performed in unicats

**Chiral gas chromatography–mass spectrometry (GC–MS).** Measurements were performed on a Lipodex E column (25 m x 0.25 mm, Macherey-Nagel, Düren, DE). 1 µl of each sample was injected with a split ratio of 1:10 (inlet temperature 200°C). The utilized temperature program can be found in **Table S3**. Ionization was obtained by electron impact with a voltage of 70 V, and the temperature of the ion source was 200 °C. Calibration and quantification were implemented in scan mode.

**Quantum Mechanics (QM) calculations.** A truncated computational model has been used to model the C-H activation and epoxidation of geraniol (1) and nerol (2) substrates. The truncated model [Fe=O(Por)(SCH3)(substrate)] includes: the active Fe-oxo species (Fe=O), the porphyrin pyrrole core (Por), a methyl thiolate group (-SCH3) to mimic Cysteine axial ligand, and 1 or 2 as substrate. Density Functional Theory (DFT) calculations were carried out using Gaussian16 software package.^5^ Geometry optimizations and frequency calculations were performed using the unrestricted hybrid (U)B3LYP^6-8^ functional with an ultrafine integration grid,^9^ and including the CPCM polarizable conductor model (dichloromethane, ε = 8.9)^10,11^ to have an estimation of the dielectric permittivity in the enzyme active site.^12^ 6-31G(d) basis set was used for all atoms but Fe, where SDD basis set and related SDD pseudopotential were employed. The optimized geometries were verified as minima by a vibrational frequency analysis and transition states geometries have a single imaginary frequency consistent with the reaction coordinate. Enthalpies and entropies were calculated for 1 atm and 298.15 K. A correction to the harmonic oscillator approximation, as discussed by Truhlar and co-workers,^13,14^ was also applied to the enthalpy calculations by raising all frequencies below 100 cm–1 to 100 cm–1 using Goodvibes v.1.0.1 python script.^15^ Single point energy calculations were performed using the functional (U)B3LYP with the Def2TZVP basis set on all atoms, and within the CPCM polarizable conductor model (dichloromethane, ε = 8.9)^10,11^ and an ultrafine integration grid.^9^ Empirical Grimme D3 dispersion corrections with Becke-Johnson (GD3BJ) damping are also included in single point calculations.^16^ All structures have a total neutral charge and calculations were performed with doublet (d) or quartet (q) multiplicities consistent with the expected electronic states of the Fe. Figs of DFT structures were rendered using CYLview^17^ and MolUP VMD extension^18^ was used for output visualization.

**Homology model and Molecular Dynamics (MD) simulations**. Homology model for *Mth*UPO structure (283 residues) obtained from our previous work has been used as starting point.^19^ Mutations were introduced using the Mutagenesis tool in PyMOL.^20^ Molecular Dynamics (MD) simulations in explicit water were performed using the AMBER18 package.^21,22^ Parameters for the geraniol (**1**) and nerol (**2**) substrates were generated within the antechamber^23^ module in AMBER18 package using the general AMBER force field (gaff2),^24^ with partial charges set to fit the electrostatic potential generated at the B3LYP/6-31G(d) level by the RESP model.^25^ The charges were calculated according to the Merz–Singh–Kollman scheme^26,27^ using the Gaussian16 package. Parameters for the heme compound I (Cpd I) and the axial Cys were taken from reference.^28^ The protein was solvated in a pre-equilibrated cubic box with a 12-Å buffer of TIP3P^29^ water molecules using the AMBER18 leap module, resulting in the addition of ∼12500 solvent molecules. The systems were neutralized by addition of explicit counterions (Na^+^ and Cl^−^). All subsequent calculations were done using the AMBER force field 14 Stony Brook (ff14SB).^30^ A two-stage geometry optimization approach was performed. The first stage minimizes the positions of solvent molecules and ions imposing positional restraints on solute by a harmonic potential with a force constant of 500 kcal·mol^-1^ Å^−2^, and the second stage is an unrestrained minimization of all the atoms in the simulation cell. The systems were gently heated using six 50 ps steps, incrementing the temperature by 50 K for each step (0–300 K) under constant-volume and periodic-boundary conditions. Water molecules were treated with the SHAKE algorithm such that the angle between the hydrogen atoms was kept fixed. Long-range electrostatic effects were modelled using the particle-mesh-Ewald method.^31^ An 8 Å cutoff was applied to Lennard–Jones and electrostatic interactions. Harmonic restraints of 30 kcal·mol^-1^ were applied to the solute, and the Langevin equilibration scheme was used to control and equalize the temperature. The time step was kept at 1 fs during the heating stages, allowing potential inhomogeneities to self-adjust. Each system was then equilibrated for 2 ns with a 2 fs time step at a constant pressure of 1 atm and temperature of 300 K without restraints. Once the systems were equilibrated in the NPT ensemble, production trajectories were then run under the NVT ensemble and periodic-boundary conditions. In particular, a total of 1500 ns from 3 independent replicas (500 ns each) were accumulated for the each of the following systems: Design 2 (F63L/A153I/F154I/G157A), Design 4 (F63I/F154L/G157A/A161F), Design 26 (F63I/F154I/G157A/A161L), and Design 28 (F63L/ F154L/Y156L/A161F). Trajectories were processed and analyzed using the CPPtraj^32^ module from Ambertools utilities. VMD visualization software was used to visualize MD simulations.^33^ Protein structures were rendered using PyMOL.^20^

**Docking and protocol used for substrate-bound MD simulations**. Docking calculations were performed using AutoDock Vina.^34^ The most populated clusters (based on backbone clustering analysis) obtained from MD simulations carried out in the absence of substrate were used, and docking predictions were then utilized as starting points for substrate-bound MD simulations. The following systems were prepared: Design 4 + geraniol, Design 2 + geraniol, Design 26 + nerol, and Design 28 + nerol. Same protocol for MD simulations described above has been employed. 3 replicas of 500 ns were carried out on each system without any external restraints on the substrate, thus accumulating a total of 1500 ns for each system. Trajectories were processed and analyzed using the CPPtraj28 module from Ambertools utilities. VMD visualization software was used to visualize MD simulations.^33^ Protein structures were rendered using PyMOL.^20^

**Further FuncLib designs for comparison with previous work**^35^

The starting structure for FuncLib design is derived from the most populated cluster of the parent F63I holoenzyme, determined by MD simulations. Key substrate-interacting positions (L56, L60, F63, L86, F154, A161, F206, and M210) were selected based on previous work for mutation to reduced alphabet A, V, I, L, and F (M allowed at position 210). The C18 axial ligand remained fixed, with one to two mutations per design. The heme cofactor and the structural Mg ion are kept on the FuncLib model. In total, 509 variants were generated in the first round with 267 variants with lower Rosetta scores than the parent enzyme.

The second round of the FuncLib evolution was set up again using the most populated cluster structure of the previous hit 12 (L60F/F63I/A161V) as a parent holoenzyme and the same parameters as the ones used before. In total 197 variants were generated in the second round with 65 variants with lower Rosetta scores than the parent enzyme.

**AI-based docking for design/substrate modeling.**

Models for each design/substrate pair were generated using Chai-1 (downloaded from <https://github.com/chaidiscovery/chai-lab>), an AlphaFold3-based and open-access predictor of biomolecular interactions. Each docking trajectory produces five docked models, and we launched several runs with different seeds to generate up to 50 models per design pair (Table S11).

We used the Simplified Molecular Input Line Entry System (SMILES) notation for heme-oxo species (Compound I) for modeling:

C=CC1=C(C)C2=CC(C(C)=C3CCC([O-])=O)=[N]4C3=CC5=[N]6C(C(C)=C5CCC([O-])=O)=CC7=C(C=C)C(C)=C(C=C1[N-]28)[N-]7[Fe]468=O

In addition, we used SMILES notations extracted from PubChem for each of the terpenes (Table S11).

Models that exhibited ≤4.5 Å between the oxidized carbon and the Compound I oxygen were defined as forming a near-attack conformation (NAC). Models that exhibited ≤ 3.4 Å distance between any atoms on the ligand and the enzyme were considered as forming steric overlaps.

**Preparative work.**

*2,3-Epoxy nerol*

Nerol (590 mg, 3.8 mmol) was dissolved in acetone (7.5 mL) and poured into a solution of potassium phosphate buffer (100 mM, 111 mL, pH 7.0) and 250 nM *Mth*UPO Var 28 (stock solution 20.5 μM, 1.8 mL). 30 mM hydrogen peroxide (stock solution 150 mM, 30 mL) was added via a syringe pump over the period of the reaction time. The solution (total: 150 mL) was stirred at 30 °C over night. Afterwards the mixture was extracted three times with ethyl acetate. The organic phase was washed with brine, dried with sodium sulfate, filtered and concentrated under reduced pressure. The crude product was purified by column chromatography on silica gel using n-hexane/ethyl acetate (9/1 → 2/1) obtaining 156 mg (70 %) 2,3-epoxy nerol as a pale-yellow oil. GC-MS analysis showed impurities of silicone grease leading to a total purity of 85% which corresponds to a yield of 132 mg (55 %) of 2,3-epoxy nerol.

**^1^H-NMR (400 MHz, CDCl_3_):** δ 5.07 (t, J 8 Hz, 1H), δ 3.37 (m, 1H), δ 3.65 (m, 1H)**,** δ 2.94 (q, J 4 Hz, 1H), δ 2.17-2.01 (m, 2H), δ 1.68-1.63 (m, 4H), δ 1.60 (s, 3H), δ 1.33 (s, 3H)

*Isopiperitenol*

(*S*)-(-)-limonene (360 mg, 2.4 mmol) was dissolved in acetone (7.5 mL) and poured into a solution of potassium phosphate buffer (100 mM, 93 mL, pH 7.0) and 200 nM *Mth*UPO Var 28 (stock solution 1.5 μM, 20 mL). 20 mM hydrogen peroxide (stock solution 100 mM, 30 mL) was added via a syringe pump over the period of the reaction time. The solution (total: 150 mL) was stirred at 30 °C over night. Afterwards the mixture was extracted three times with ethyl acetate. The organic phase was washed with brine, dried with sodium sulfate, filtered and concentrated under reduced pressure. The crude product was purified by column chromatography on silica gel using n-hexane/ethyl acetate (19/1 → 5/1) obtaining 3 mg (1 %) isopiperitenol as a pale-yellow oil. GC-MS analysis showed impurities of silicone grease leading to an even lower yield of isolated product.

**^1^H-NMR (400 MHz, CDCl_3_):**

δ 5.42 (s, 1H), δ 4.86 (d, J 20 Hz, 2H), δ 4.15-4.05 (m, 2H), δ 2.11 (s, 1H), δ 2.05 (m, 1H), δ 1.71 (s, 3H), δ 1.68 (s, 3H), 1.41 (d, J 8 Hz, 2H)

**Calibration curves.**

In every case, each sample was prepared in triplicates. The samples were extracted with 0.5 mL EtOAc containing an internal standard by vortexing for 30 seconds and centrifuged at 3000 rpm for 1 min to separate the aqueous and the organic phase. The organic phase was transferred to a GC vial with glass insert and analyzed via GC-MS.

**Citral A and B**

The product was available as mix of both isomers citral A and B, in a first step the ratio of both products and thus the purity of each isomer was determined. Afterwards, five stock solutions (15 mM, 10 mM, 5 mM, 2 mM, and 1 mM) in acetone were prepared. In each case 0.025 mL were added to 0.475 mL of buffered solution (0.1 M Kpi, pH 7) with final concentrations of 0.75 mM, 0.5 mM, 0.2 mM, 0.1 mM, and 0.05 mM. 0.5 mM β‑ionone was used as internal standard.

**2,3-Epoxy geraniol and nerol**

Four stock solutions (15 mM, 10 mM, 5 mM, and 2 mM) in acetone were prepared. In each case 0.025 mL were added to 0.475 mL of buffered solution (0.1 M Kpi, pH 7) with final concentrations of 0.75 mM, 0.5 mM, 0.2 mM, and 0.1 mM. 0.5 mM β‑ionone was used as internal standard.

**Carveol**

Four stock solutions (10 mM, 5 mM, 2 mM, and 1 mM) in acetone were prepared. In each case 0.025 mL were added to 0.475 mL of buffered solution (0.1 M Kpi, pH 7) with final concentrations of 0.5 mM, 0.2 mM, 0.1 mM, and 0.05 mM. 0.25 mM 4-octanone was used as internal standard. The calibration curve of carveol was used for TON determination of both carveol and mix-10-ol (consisting of isopiperitenol and another product, maybe limonene-10-ol, that elute at the same time).

**Identification of products.**

Identification of the different products was performed through mass spectrum comparison with the literature (**Table S4**), elution time comparison with product standards and NMR for isopiperitenol. To ensure correct epoxide identification the injection temperature was lowered to 200 °C for side product identification runs.

**Gene and amino acid sequence of wild type *Mth*UPO** with signal peptide from α Galactosidase (*S.cerevisiae*, underline) and TwinStrep-GFP11 tag (italic) in pAGT572_Nemo 2.0 used for enzyme expression in *S. cerevisiae*:

ATGTTTGCTTTTTATTTCTTGACTGCTTGTATTTCTTTGAAAGGTGTTTTTGGAGCAGGTTTTGACACTTGGTCACCACCTGGACCCTATGATGTTAGGGCTCCTTGTCCGATGTTGAATACATTGGCTAATCATGGTTTCTTACCACATGATGGCAAAGATATTACTCGTGAACAGACAGAAAACGCCTTGTTCGAAGCATTGCACATCAACAAAACCTTAGCCAGCTTTCTGTTTGACTTTGCATTAACAACGAATCCGAAGAATACCTCGACGTTTTCACTGAACGACTTAGGCAATCACAACATTTTGGAACATGATGCATCACTAAGTAGGGCTGATGCGTACTTTGGGAATGTTCTACAGTTCAATCAAACTGTCTTTGATGAGACTAAAACCTACTGGGAAGGAGATACTATTGATTTGAGAATGGCAGCCAAAGCTAGACTAGGTAGAATCAAGACATCTCAAGCTACTAATCCAACGTATTCCATGTCGGAATTAGGAGATGCTTTCACATATGGGGAATCTGCTGCGTATGTGGTAGTGTTAGGTGACAAAGAGTCTCGTACTGTCAAAAGATCCTGGGTTGAATGGTTCTTCGAACATGAGCAACTTCCTCAACATCTTGGTTGGAAAAGACCAGCAGCATCCTTCGAAGAAGAAGATCTGAACTCATCAATGGAGGAGATTGAGAAGTACACCAAGGAATTGGAAGGTAGCAACTCTACAAGTGGTAGTCAAAAGCATAGAAGGAGACTTCCAAGAAGAAGAGCTCACTTTGGCTTTTCGGGTGGTTCTGCTTGGTCACATCCACAATTTGAAAAAGGTGGAGGTTCAGGTGGAGGT*TCGGGTGGTTCTGCTTGGTCACATCCACAATTTGAAAAAGATGGTGGTTCTGGTGGTGGTTCTACTAGTCGTGATCATATGGTTCTTCATGAATATGTTAATGCTGCT GGTATTACTTGA*

MFAFYFLTACISLKGVFGAGFDTWSPPGPYDVRAPCPMLNTLANHGFLPHDGKDITREQTENALFEALHINKTLASFLFDFALTTNPKNTSTFSLNDLGNHNILEHDASLSRADAYFGNVLQFNQTVFDETKTYWEGDTIDLRMAAKARLGRIKTSQATNPTYSMSELGDAFTYGESAAYVVVLGDKESRTVKRSWVEWFFEHEQLPQHLGWKRPAASFEEEDLNSSMEEIEKYTKELEGSNSTSGSQKHRRRLPRRRAHFGF*SGGSAWSHPQFEKGGGSGGGSGGSAWSHPQFEKDGGSGGGSTSRDHMVLHEYVNAAGIT*

\#

**II Supplementary Tables**

**Table S1** Explored sequence space during FuncLib creation.

| **Position** | **Sequence Space** |
| --- | --- |
| F59 | FYQ |
| L60 | LFIQ |
| F63 | FILQV |
| L86 | LIMV |
| A153 | AILQV |
| F154 | FILMV |
| Y156 | YIL |
| G157 | GAS |
| S159 | SAIYQ |
| A161 | AFILM |
| L206 | LA |
| M210 | MAFI |

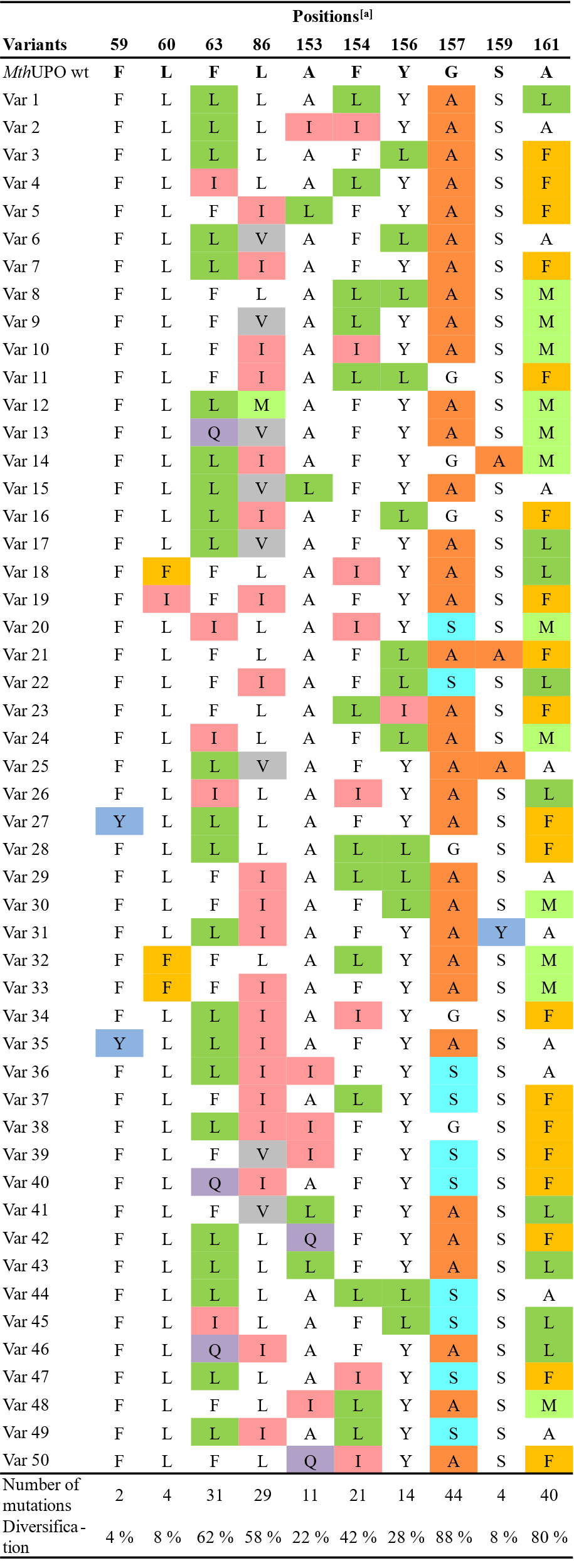
**Table S2** Targeted positions and corresponding amino acid substitutions in each FuncLib variant

^[a]^Positions L206 and M210 were also available for mutation, but no diversification occurred among the first 50 variants. Mutations are highlighted in different colours.

**Table S3** GC-MS parameter.

| **Substrate** | **GC-MS** | **Split** | **Temperature program** | **Internal standard** |
| --- | --- | --- | --- | --- |
| α-damascone  β-damascone  δ-damascone | GCMS-QP2010 Ultra | 20 | 110 °C  4 °C/min to 185 °C  50 °C/min to 300 °C  hold 2 min | β-ionone |
| α-ionone | GCMS-QP2010 Ultra | 10 | 110 °C  4 °C/min to 185 °C  50 °C/min to 300 °C  hold 2 min | β-ionone |
| geraniol  nerol | GCMS-QP2010 Ultra | 10 | 100 °C  4 °C/min to 175 °C  50 °C/min to 300 °C  hold 2 min | β-ionone |
| *R*-(+)-limonene  *S*-(-)-limonene | GCMS-QP2010 Ultra | 20 | 70 °C  4 °C/min to 145 °C  100 °C/min to 300 °C  hold 2 min | 4-octanone |
| β-ionone | GCMS-QP2010 Ultra | 10 | 110 °C  4 °C/min to 185 °C  50 °C/min to 300 °C  hold 2 min | α-ionone |
| valencene | GCMS-QP2010 Ultra | 20 | 110 °C  10 °C/min to 300 °C  hold 2 min | naphthalene |
| β-ionone  (chiral measurement) | GCMS-QP2010 Ultra | 10 | 45 °C  2 °C/min to 80 °C  1 °C/min to 170 °C  20 °C/min to 200 °C  hold 10 min | - |

**Table S4** Chemical structures and mass fragmentation of oxyfunctionalization products from enzymatic reactions with *Mth*UPO.

| **Nr.** | **Name** | **Structure** | **GC-MS (m/z)** | **Identification / Literature** |
| --- | --- | --- | --- | --- |
|  | 4-epoxy-α-ionone  (4-epoxy- α-I) | 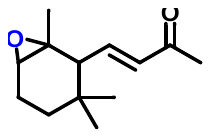 | 208 [M^+^], 193 [M-CH_3_]^+^, 121 (24), 111 (37), 109 (100), 107 (24), 95 (56), 93 (30), 67 (21) | ^36^ |
|  | 3-hydroxy-α- ionone  (3-OH-α-I) | 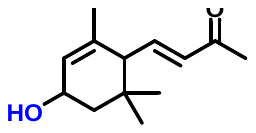 | 208 [M^+^], 193 [M-CH_3_]^+^, 125 (29), 124 (25), 109 (100), 107 (23), 91 (19), 81 (29) | ^36^ |
|  | 3-oxo-α- ionone  (3-CO-α-I) | 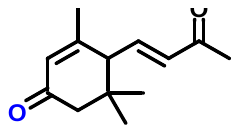 | 206 [M^+^], 193 [M-CH_3_]^+^, 150 (11), 108 (100), 107 (15), 91 (8), 77 (10) | ^36^ |
|  | 4-hydroxy-β-ionone  (4-OH-β-I) | 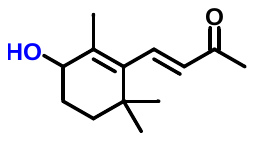 | 208 [M^+^] (28), 193 [M-CH_3_]^+^ (14), 175 (15), 165 (12), 151 (14), 137 (26), 123 (37), 109 (100), 107 (17), 95 (20), 91 (25) | ^35^ |
|  | 7,11-epoxymega­stigma-5(6)-en-9-one  (EME) | 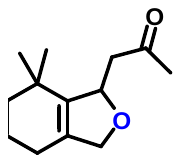 | 208 [M^+^], 193 [M-CH_3_]^+^, 151 (36) 150 (92), 135 (100), 107 (28), 95 (32), 81 (52), 79 (22) | ^36^ |
|  | not identified product |  | 208 [M+], 193 [M-CH_3_]^+^ (98), 175 (33), 131 (21), 109 (20), 105 (33), 91 (28), 79 (17) |  |
|  | 2-hydroxy-β-ionone  (2-OH-β-I) | 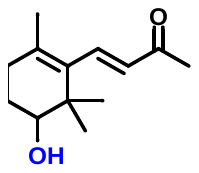 | 208 [M+], 193 [M-CH_3_]^+^ (67), 175 (41), 121 (42), 149 (33), 105 (37), 91 (26), 79 (21) | ^36^ |
|  | 4-epoxy-α-damascone  (4-epoxy-α-I) | 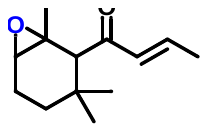 | 208 [M+], 193 [M-CH_3_]^+^, 137 (90), 111 (27), 109 (30), 95 (31), 81 (25), 69 (100) | ^36^ |
|  | 3-hydroxy-α-damascone  (3-OH-α-D) | 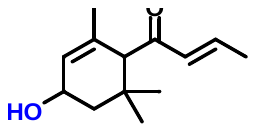 | 208 [M+], 193 [M-CH_3_]^+^,151 (29), 123 (22), 109 (31), 107 (73), 91 (17), 69 (100) | ^36^ |
|  | not identified |  | 208 [M+], 193 [M-CH_3_]^+^, 122 (33), 107 (100), 105 (12), 91 (15), 69 (51) |  |
|  | 4-oxo-β-damascone  (4-CO-β-D) | 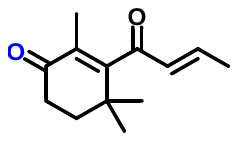 | 206 [M+], 191 [M-CH_3_]^+^ (18), 138 (22), 123 (15), 69 (100) | ^36^ |
|  | 4-hydroxy-β-damascone  (4-OH-β-D) | 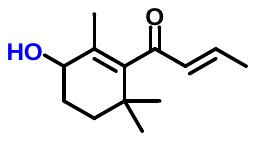 | 208 [M+], 193 [M-CH_3_]^+^ (23), 139 (98), 123 (25), 109 (26), 105 (26), 91 (25), 69 (100) | ^36^ |
|  | 3-hydroxy-β-damascone  (3-OH-β-D) | 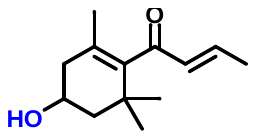 | 208 [M+], 193 [M-CH_3_]^+^ (42), 175 (67), 147 (28), 121 (82), 119 (33), 105 (51), 93 (30), 69 (100) | ^36^ |
|  | 2-hydroxy-δ-damascone  (2-OH-δ-D) | 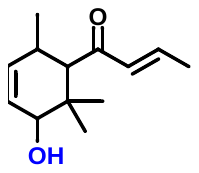 | 208 [M+], 193 [M-CH_3_]^+^, 124 (11), 109 (23), 107 (46), 97 (14), 69 (100) | ^37^, fragmentation pattern (Fig. S2) |
|  | not identified |  | 125 (28), 120 (33), 109 (30), 107 (54), 69 (100) |  |
|  | citral B (neral) | 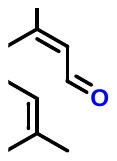 | 152 [M+], 137 [M-CH_3_]^+^ (9),109 (34), 94 (36), 84 (31), 69 (100), 67 (28) | product standard |
|  | neric acid (TMS) | 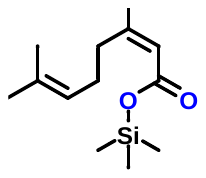 | 240 [M+] (2), 225 [M-CH_3_]^+^(3), 156 (34), 122 (34), 112 (30), 82 (90), 75 (56), 73 (78), 69 (100) | NIST library comparison |
|  | 2,3-epoxynerol | 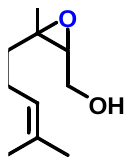 | 170 [M+], 155 [M-CH_3_]^+^, 109 (56), 82 (34), 69 (66), 67 (67), 41 (100) | ^38^ |
|  | 6,7-epoxynerol | 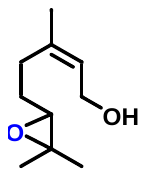 | 170 [M+], 155 [M-CH_3_]^+^, 85 (62), 84 (51), 81 (46), 71 (48), 59 (92), 57 (46), 41 (100) | ^38^ |
|  | citral A (geranial) | 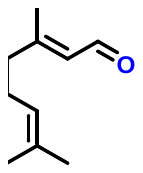 | 152 [M+], 137 [M-CH_3_]^+^ (16),109 (13), 94 (17), 84 (31), 69 (100), 67 (10) | product standard |
|  | 2,3-epoxygeraniol | 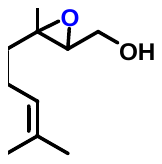 | 170 [M+], 155 [M-CH_3_]^+^, 109 (58), 82 (44), 69 (87), 67 (99), 41 (100) | ^38^ |
|  | 6,7-epoxygeraniol | 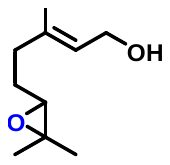 | 170 [M+], 155 [M-CH_3_]^+^, 85 (62), 81 (84), 71 (60), 59 (100), 57 (62), 41 (87) | ^38^ |
|  | (-)-limonene oxide (1) | 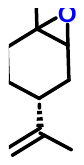 | 152 [M+] (1), 137 [M-CH_3_]^+^ (32), 109 (37), 93 (45), 81 (33), 79 (45), 67 (86), 43 (100) | product standard |
|  | (-)-limonene oxide (2) | 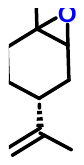 | 152 [M+], 137 [M-CH_3_]^+^ (11), 108 (37), 94 (52), 81 (40), 79 (52), 67 (76), 43 (100) | product standard |
|  | (-)-isopiperitenol | 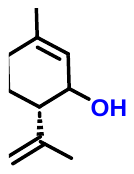 | 152 [M+] (1), 137 [M-CH_3_]^+^ (4), 119 (11), 91 (21), 84 (100), 79 (13), 56 (16) | NMR^39^ |
|  | (-)-carveol | 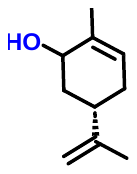 | 152 [M+] (2), 137 [M-CH_3_]^+^ (4), 109 (15), 94 (18), 91 (19), 84 (100), 79 (21), 69 (15), 56 (20) | product standard |
|  | not identified (probably limonene-10-ol) | 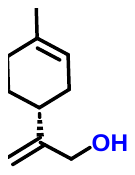 | 152 [M+] (2), 137 [M-CH_3_]^+^ (12), 121 (83), 105 (40), 94 (60), 93 (71), 91 (67), 84 (76), 79 (100), 67 (47) |  |
|  | not identified |  | 152 [M+] (1), 137 [M-CH_3_]^+^ (14), 121 (58), 105 (35), 94 (59), 93 (71), 91 (57), 79 (100), 77 (37), 67 (52) |  |
|  | (+)-limonene oxide (1) | 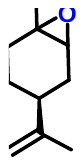 | 152 [M+] (2), 137 [M-CH_3_]^+^ (33), 109 (38), 93 (45), 81 (32), 79 (41), 67 (85), 43 (100) | product standard |
|  | (+)-limonene oxide (2) | 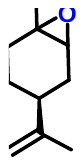 | 152 [M+], 137 [M-CH_3_]^+^ (5), 108 (40), 94 (62), 81 (41), 79 (50), 67 (73), 43 (100) | product standard |
|  | (+)-isopiperitenol | 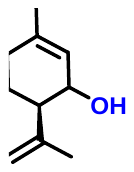 | 152 [M+] (1), 137 [M-CH_3_]^+^ (7), 121 (33), 91 (39), 84 (100), 79 (42), 67 (24) | NMR^39^ |
|  | not identified (probably limonene-10-ol) | 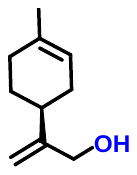 | 152 [M+] (1), 137 [M-CH_3_]^+^ (15), 121 (100), 105 (44), 93 (81), 91 (72), 84 (45), 79 (97), 67 (48) |  |
|  | (+)-carveol | 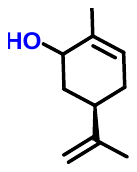 | 152 [M+] (2), 137 [M-CH_3_]^+^ (7), 109 (70), 91 (32), 84 (100), 79 (21), 69 (23), 56 (23) | product standard |
|  | not identified |  | 152 [M+] (1), 137 [M-CH_3_]^+^ (15), 121 (56), 105 (41), 94 (64), 93 (73), 91 (67), 79 (100), 77 (36), 67 (43) |  |

**Table S5** Turnover numbers of (*R*)-(+)-limonene and (*S*)-(-)-limonene products for selected FuncLib variants.

| **Enzyme** | **Substrate** | **Turnover Numbers (TON)** | |
| --- | --- | --- | --- |
|  |  | **Mix Lim-ol^[a]^** | **Carveol** |
| Var 45 | (*R*)-(+)-limonene | 70±3 | 340±10 |
| wt | (*R*)-(+)-limonene | 230±11 | 60±5 |
| Var 45 | (*S*)-(-)-limonene | 310±40 | 50±3 |
| wt | (*S*)-(-)-limonene | 12±2 | 12±1 |
| [a] mixture of isopiperitenol and another product, maybe, limonene-10-ol, eluting both at the same time | | | |

**Table S6** Regioselectivity of 3-hydroxy-α-ionone and diastereomric ratio of both 3-hydroxy-α-ionone diastereomeres for selected FuncLib variants.

| **Enzyme** | **Regioselectivity 3-OH-α-I** | **d.r.**  **3-OH-α-I** |
| --- | --- | --- |
| Var 18 | 98 % | 2:98 |
| Var 45 | 99 % | 15:85 |
| wt | 91 % | 12:88 |

**Table S7 Energies and thermochemistry parameters of all DFT structures reported in Fig. S6**.

Energies and thermochemistry parameters (at T = 298.15 K and P = 1 atm) of all DFT optimized stationary points reported in **Fig. S6**. Electronic energies (E), enthalpy (H), free energy (G), quasi harmonic corrected free energy (G-qh), electronic energies from high level single point calculations (E(SP)), and imaginary frequencies for the TS. Energies and frequencies are given in a.u. and cm^-1^.

| **Cpd I + Geraniol** | **Electronic State** | **E** | **H** | **G** | **G-qh** | **E(SP)** | **Imag. Freq** |
| --- | --- | --- | --- | --- | --- | --- | --- |
| **TS C2-O formation** | doublet (d) | -2092.676457 | -2092.059380 | -2092.173424 | -2092.171701 | -3233.342682 | 427.7i |
|  | quartet (q) | -2092.675786 | -2092.058498 | -2092.171871 | -2092.170118 | -3233.342579 | 416.4i |
| **TS C1-HAT** | doublet (d) | -2092.676749 | -2092.064277 | -2092.179030 | -2092.176207 | -3233.339982 | 1990.5i |
|  | quartet (q) | -2092.677116 | -2092.065004 | -2092.177325 | -2092.175477 | -3233.343119 | 1569.2i |
| **TS C4-HAT** | doublet (d) | -2092.664984 | -2092.053853 | -2092.167345 | -2092.165676 | -3233.336364 | 1814.3i |
|  | quartet (q) | -2092.665536 | -2092.054319 | -2092.167296 | -2092.165898 | -3233.336749 | 1917.7i |
| **TS C5-HAT** | doublet (d) | -2092.666019 | -2092.053922 | -2092.166633 | -2092.164939 | -3233.338410 | 910.5i |
|  | quartet (q) | -2092.664891 | -2092.053127 | -2092.166914 | -2092.164988 | -3233.335709 | 1745.1i |
| **TS C8-HAT** | doublet (d) | -2092.668543 | -2092.055992 | -2092.172137 | -2092.168783 | -3233.332926 | 908.6i |
|  | quartet (q) | -2092.667835 | -2092.056061 | -2092.171090 | -2092.168361 | -3233.333114 | 1804.8i |

**Table S8 Energies and thermochemistry parameters of all DFT structures reported in Fig. S7**.

Energies and thermochemistry parameters (at T = 298.15 K and P = 1 atm) of all DFT optimized stationary points reported in **Fig. S7**. Electronic energies (E), enthalpy (H), free energy (G), quasi harmonic corrected free energy (G-qh), electronic energies from high level single point calculations (E(SP)), and imaginary frequencies for the TS. Energies and frequencies are given in a.u. and cm^-1^.

| **Cpd I + Nerol** | **Electronic State** | **E** | **H** | **G** | **G-qh** | **E(SP)** | **Imag. Freq** |
| --- | --- | --- | --- | --- | --- | --- | --- |
| **TS C2-O formation** | doublet (d) | -2092.676947 | -2092.059721 | -2092.173215 | -2092.171300 | -3233.345299 | 447.3i |
| (conformer 1) | quartet (q) | -2092.675772 | -2092.058432 | -2092.170153 | -2092.168889 | -3233.343573 | 425.2i |
| **TS C2-O formation** | doublet (d) | -2092.684172 | -2092.066749 | -2092.176974 | -2092.175914 | -3233.351789 | 330.2i |
| (conformer 2) | quartet (q) | -2092.682501 | -2092.065190 | -2092.175118 | -2092.174047 | -3233.351443 | 423.1i |
| **TS C1-HAT** | doublet (d) | -2092.675913 | -2092.063148 | -2092.176817 | -2092.174418 | -3233.338928 | 1946.5i |
|  | quartet (q) | -2092.676163 | -2092.063771 | -2092.176228 | -2092.174314 | -3233.340942 | 1527.9i |
| **TS C4-HAT** | doublet (d) | -2092.666498 | -2092.054786 | -2092.166546 | -2092.165161 | -3233.339990 | 1044.2i |
|  | quartet (q) | -2092.665399 | -2092.053886 | -2092.164638 | -2092.163646 | -3233.338742 | 1864.3i |
| **TS C5-HAT** | doublet (d) | -2092.666190 | -2092.054013 | -2092.166757 | -2092.164763 | -3233.340396 | 946.8i |
|  | quartet (q) | -2092.665321 | -2092.053676 | -2092.165825 | -2092.164151 | -3233.340209 | 1768.3i |
| **TS C8-HAT** | doublet (d) | -2092.664861 | -2092.052733 | -2092.167188 | -2092.164358 | -3233.329377 | 1480.2i |
|  | quartet (q) | -2092.669458 | -2092.057613 | -2092.172305 | -2092.169503 | -3233.335003 | 1818.7i |

**Table S8** Cartesian coordinates (xyz, in Å) of all DFT optimized structures in **Fig. S6** and **Table S7**.

```
Geraniol (1)
 O  -3.939366 -1.810334  0.258374
 H  -4.888995 -2.006158  0.295011
 C  -3.802368 -0.486664 -0.279572
 H  -4.206891  0.259801  0.417015
 H  -4.381041 -0.401889 -1.214941
 C  -2.349563 -0.253304 -0.569924
 H  -1.895475 -1.046633 -1.165415
 C  -1.580691  0.787080 -0.210249
 C  -2.059529  1.976331  0.588633
 H  -1.394960  2.174071  1.438861
 H  -2.046611  2.883050 -0.032123
 H  -3.072938  1.855592  0.978295
 C  -0.121420  0.826834 -0.624843
 H   0.058440  0.098124 -1.423739
 H   0.115756  1.817136 -1.041603
 C   0.864158  0.532749  0.536618
 H   0.673888  1.242313  1.354798
 H   0.648164 -0.463421  0.935671
 C   2.301162  0.661512  0.106190
 H   2.583685  1.667481 -0.212557
 C   3.249733 -0.287497  0.049112
 C   3.053168 -1.733030  0.438382
 H   3.739828 -2.011887  1.249945
 H   2.036428 -1.960265  0.766733
 H   3.290329 -2.394814 -0.406237
 C   4.645323  0.044524 -0.426712
 H   4.745293  1.100419 -0.697992
 H   5.391679 -0.180829  0.348301
 H   4.918514 -0.560630 -1.302816

TS C2-O formation
(d, Cpd I + Geraniol)
 C   2.947069 -0.427962 -3.285195
 S   2.609376  1.004223 -2.198180
 H   3.486491 -1.215905 -2.752634
 H   3.582260 -0.067478 -4.102155
 N   2.659855 -1.104880  0.091343
 N   2.275853  1.657467  0.726731
 N   0.277840 -1.065570 -1.457474
 C   1.702366 -3.035488 -1.077522
 C   4.160076  0.267952  1.470092
 C   0.789951  3.588424  0.433040
 C  -1.541407  0.354347 -2.312505
 C   2.663676 -2.425651 -0.281062
 C   3.450556  1.459762  1.409491
 C  -0.142407  2.989774 -0.404785
 C  -0.855076 -0.848995 -2.209665
 O   0.420124 -0.112194  1.165252
 C   3.823757 -3.092932  0.261487
 C   3.845199  2.676857  2.078815
 C  -1.273334  3.675924 -0.984653
 C  -1.249783 -2.066919 -2.871755
 C   4.519103 -2.158749  0.966407
 C   2.892298  3.607503  1.797885
 C  -1.921174  2.772268 -1.769831
 C  -0.348580 -3.023545 -2.511338
 C   3.782773 -0.921086  0.860719
 C   1.915404  2.962192  0.952731
 C  -1.190112  1.531443 -1.664125
 C   0.604703 -2.391561 -1.634450
 N  -0.113603  1.684943 -0.828382
 Fe  1.209397  0.256806 -0.276425
 H   1.831001 -4.089490 -1.302403
 H   5.076719  0.261867  2.051277
 H   0.636773  4.632531  0.686891
 H  -2.421690  0.374423 -2.947039
 C  -0.282240 -2.402762  2.430832
 O  -1.021049 -3.402633  3.148645
 C  -1.186345 -1.452316  1.681431
 C  -2.026003 -0.544995  2.288218
 C  -1.945411 -0.223798  3.753188
 C  -3.049496  0.201653  1.478896
 C  -4.486675 -0.376400  1.651162
 C  -5.511092  0.446887  0.918189
 C  -6.286955  0.089551 -0.118805
 C  -6.288021 -1.271374 -0.772800
 C  -7.263597  1.071861 -0.723666
 H  -1.542861 -3.902555  2.500230
 H   0.424553 -2.864627  1.731650
 H   0.297644 -1.869238  3.186210
 H  -1.365208 -1.678174  0.633549
 H  -1.516566 -1.037584  4.340895
 H  -1.323119  0.672035  3.900913
 H  -2.935136  0.013227  4.159885
 H  -3.062853  1.256780  1.787010
 H  -2.787930  0.174428  0.416288
 H  -4.489755 -1.419181  1.318890
 H  -4.733738 -0.389082  2.722292
 H  -5.610802  1.469421  1.288153
 H  -5.566296 -1.965121 -0.335456
 H  -7.283338 -1.732085 -0.703521
 H  -6.063864 -1.182599 -1.844870
 H  -8.292994  0.690655 -0.666490
 H  -7.232000  2.043538 -0.220493
 H  -7.052538  1.231870 -1.790505
 H   2.029483 -0.842071 -3.710744
 H   5.444490 -2.276508  1.515606
 H   4.058828 -4.138378  0.107738
 H   4.738348  2.783492  2.681094
 H   2.839133  4.639925  2.119012
 H  -1.516751  4.715708 -0.807314
 H  -2.810372  2.912034 -2.371108
 H  -2.108106 -2.158986 -3.524949
 H  -0.311716 -4.063691 -2.809215

TS C2-O formation
(q, Cpd I + Geraniol)
 C   3.344879 -0.541700 -2.964278
 S   2.297020  0.838353 -2.385487
 H   4.136993 -0.776191 -2.249459
 H   3.791775 -0.238159 -3.916581
 N   2.708654 -1.056682  0.138785
 N   2.273050  1.699413  0.689177
 N   0.288435 -1.154716 -1.374656
 C   1.757825 -3.063919 -0.922884
 C   4.199578  0.385570  1.453347
 C   0.784750  3.610839  0.277200
 C  -1.553874  0.207075 -2.258480
 C   2.719215 -2.397580 -0.174553
 C   3.461039  1.556832  1.364422
 C  -0.138376  2.968172 -0.542247
 C  -0.864328 -0.989580 -2.102551
 O   0.370123 -0.100367  1.132653
 C   3.893695 -3.025608  0.380755
 C   3.843863  2.811759  1.969274
 C  -1.289408  3.605591 -1.136508
 C  -1.264654 -2.248335 -2.687318
 C   4.589134 -2.054364  1.034817
 C   2.877617  3.713909  1.648121
 C  -1.939380  2.649607 -1.856796
 C  -0.338695 -3.171736 -2.305744
 C   3.837418 -0.831474  0.887014
 C   1.903351  3.015185  0.841570
 C  -1.188071  1.426705 -1.701132
 C   0.629852 -2.476173 -1.489822
 N  -0.090988  1.647595 -0.902931
 Fe  1.265019  0.271472 -0.284039
 H   1.906830 -4.125753 -1.092607
 H   5.124089  0.417499  2.021088
 H   0.624018  4.665066  0.479693
 H  -2.453877  0.188068 -2.865146
 C  -0.289587 -2.297728  2.410025
 O  -1.063467 -3.295620  3.091352
 C  -1.160700 -1.328336  1.642396
 C  -2.038308 -0.448475  2.260369
 C  -1.950684 -0.127824  3.725867
 C  -3.080258  0.274304  1.453484
 C  -4.507711 -0.319481  1.648927
 C  -5.551997  0.476990  0.914095
 C  -6.338383  0.090314 -0.104247
 C  -6.333965 -1.283931 -0.729959
 C  -7.333559  1.049544 -0.716032
 H  -1.561601 -3.788822  2.419666
 H   0.434720 -2.759603  1.729924
 H   0.268494 -1.775932  3.189613
 H  -1.372568 -1.591350  0.610049
 H  -1.238451  0.696091  3.888060
 H  -2.916974  0.208210  4.117067
 H  -1.615464 -0.981214  4.320254
 H  -3.104813  1.333620  1.747920
 H  -2.830525  0.238850  0.388414
 H  -4.499814 -1.367528  1.333318
 H  -4.743212 -0.319250  2.722900
 H  -5.658022  1.505894  1.264359
 H  -5.599992 -1.961236 -0.287269
 H  -7.323626 -1.752954 -0.638564
 H  -6.124037 -1.215599 -1.806449
 H  -8.358109  0.659285 -0.636539
 H  -7.305247  2.031972 -0.233926
 H  -7.138959  1.189179 -1.788883
 H   2.741353 -1.437963 -3.137305
 H   5.521756 -2.139501  1.577790
 H   4.137071 -4.074956  0.272761
 H   4.741221  2.961217  2.556048
 H   2.815054  4.761401  1.914144
 H  -1.549397  4.648481 -1.007206
 H  -2.846318  2.742741 -2.440446
 H  -2.139944 -2.386256 -3.309334
 H  -0.294229 -4.225510 -2.549993

TS C1-HAT formation
(d, Cpd I + Geraniol)
 C   3.974442 -0.205218  2.314093
 S   3.370507  1.080592  1.160308
 H   3.253328 -0.396207  3.113460
 H   4.897211  0.175836  2.765128
 N   1.913191 -1.581818  0.170341
 N   0.568649  0.094197  2.027847
 N   2.157481  0.590949 -1.654143
 C   3.074772 -1.655598 -1.997864
 C   0.991587 -2.300240  2.335570
 C  -0.225462  2.384660  2.428129
 C   1.581151  2.947844 -2.030900
 C   2.632966 -2.225366 -0.811380
 C   0.492760 -1.071355  2.748291
 C   0.151985  2.941225  1.213724
 C   2.148552  1.737098 -2.407935
 O  -0.246386  0.009189 -0.520083
 C   2.844103 -3.602993 -0.446866
 C  -0.168961 -0.834831  4.009306
 C  -0.095813  4.311357  0.833308
 C   2.845830  1.518097 -3.653333
 C   2.239664 -3.790781  0.760037
 C  -0.492926  0.487204  4.040023
 C   0.398583  4.466791 -0.425610
 C   3.281156  0.228296 -3.638418
 C   1.666928 -2.525059  1.142502
 C  -0.038009  1.058776  2.794821
 C   0.955296  3.193556 -0.815898
 C   2.843106 -0.343130 -2.387207
 N   0.795962  2.279780  0.196492
 Fe  1.277159  0.312724  0.146883
 H   3.631125 -2.290850 -2.679503
 H   0.867325 -3.144056  3.006709
 H  -0.727222  3.032793  3.139513
 H   1.647073  3.771388 -2.734744
 C  -1.068820 -2.242532 -1.643300
 O  -0.462188 -2.311950 -2.907358
 C  -2.546600 -2.155794 -1.646169
 C  -3.382408 -2.463809 -0.628962
 C  -2.913176 -2.987776  0.704930
 C  -4.875372 -2.284466 -0.788745
 C  -5.502840 -1.211725  0.147421
 C  -4.976598  0.176884 -0.098682
 C  -5.654422  1.270564 -0.485481
 C  -7.139461  1.323824 -0.755269
 C  -4.940604  2.588721 -0.681673
 H  -0.875242 -1.648194 -3.485180
 H  -0.661087 -1.255101 -1.089164
 H  -0.655133 -3.057520 -1.043097
 H  -2.981667 -1.741746 -2.557410
 H  -2.995984 -2.213719  1.479500
 H  -3.538631 -3.828376  1.031938
 H  -1.871329 -3.317710  0.692656
 H  -5.109851 -2.025021 -1.827508
 H  -5.370580 -3.243053 -0.573578
 H  -6.589243 -1.255459  0.022708
 H  -5.304164 -1.498657  1.190397
 H  -3.901953  0.288031  0.054305
 H  -7.332022  1.670682 -1.780122
 H  -7.641022  0.361692 -0.627419
 H  -7.627339  2.048604 -0.088401
 H  -5.359685  3.366022 -0.026764
 H  -3.868812  2.508909 -0.473028
 H  -5.062459  2.954380 -1.711234
 H   4.196675 -1.142045  1.796852
 H   2.979563  2.268745 -4.421755
 H   3.845101 -0.304741 -4.393111
 H   3.384451 -4.320505 -1.050847
 H   2.184431 -4.693537  1.354761
 H  -0.348214 -1.593350  4.760555
 H  -0.997994  1.041643  4.820595
 H   0.400041  5.353187 -1.046871
 H  -0.587764  5.042595  1.461769

TS C1-HAT formation
(q, Cpd I + Geraniol)
 C   4.539645  0.278563  1.264551
 S   2.883515  0.648567  1.935639
 H   4.598600 -0.769917  0.955312
 H   5.267320  0.444719  2.064881
 N   2.565245  0.491502 -1.211188
 N   1.682155 -1.651503  0.468557
 N   0.788293  2.277363  0.108792
 C   2.289997  2.872686 -1.739044
 C   3.293684 -1.855089 -1.365797
 C   0.265067 -2.204982  2.391919
 C  -0.827692  2.496720  1.945039
 C   2.857912  1.618619 -1.937056
 C   2.543522 -2.373414 -0.314485
 C  -0.268386 -0.942032  2.616110
 C  -0.101467  2.995028  0.873085
 O  -0.065923 -0.054330 -0.837188
 C   3.817250  1.311191 -2.969708
 C   2.574349 -3.751728  0.117899
 C  -1.230809 -0.639612  3.647347
 C  -0.144338  4.369238  0.428083
 C   4.086441 -0.020699 -2.871901
 C   1.723788 -3.848810  1.177361
 C  -1.542439  0.680985  3.522120
 C   0.738381  4.477704 -0.600992
 C   3.296691 -0.527580 -1.776202
 C   1.173516 -2.530149  1.391700
 C  -0.770178  1.187461  2.414128
 C   1.320944  3.169322 -0.791445
 N   0.007207  0.190718  1.883731
 Fe  1.205956  0.331972  0.261979
 H   2.620000  3.678486 -2.386985
 H   3.931876 -2.544385 -1.909829
 H  -0.054716 -3.003550  3.054041
 H  -1.488695  3.184002  2.463523
 C  -0.889299 -2.354615 -1.664732
 O  -0.141093 -2.537871 -2.836293
 C  -2.338246 -2.158166 -1.841436
 C  -3.323363 -2.432745 -0.955310
 C  -3.084697 -3.067367  0.392217
 C  -4.761698 -2.104950 -1.290596
 C  -5.430766 -1.066440 -0.345642
 C  -4.761339  0.281925 -0.363180
 C  -5.269515  1.461702 -0.756865
 C  -6.671281  1.674401 -1.277511
 C  -4.431946  2.719117 -0.699679
 H  -0.460289 -1.909664 -3.506095
 H  -0.466195 -1.289293 -1.136087
 H  -0.590116 -3.129414 -0.956420
 H  -2.620079 -1.670248 -2.776830
 H  -3.169467 -2.326108  1.198046
 H  -3.838023 -3.839982  0.593432
 H  -2.095026 -3.522711  0.478618
 H  -4.828479 -1.737421 -2.321116
 H  -5.355669 -3.030010 -1.237119
 H  -6.486083 -0.992170 -0.626376
 H  -5.416054 -1.465995  0.678923
 H  -3.727560  0.279679 -0.014423
 H  -6.644844  2.113082 -2.284800
 H  -7.261970  0.756666 -1.327648
 H  -7.214412  2.390199 -0.644444
 H  -4.909020  3.482244 -0.067926
 H  -3.429029  2.527365 -0.304356
 H  -4.325375  3.167941 -1.697576
 H   4.779222  0.915829  0.410358
 H  -0.769079  5.136407  0.867310
 H   0.990850  5.352540 -1.186478
 H   4.213975  2.031220 -3.673990
 H   4.752102 -0.621848 -3.478059
 H   3.176916 -4.526388 -0.338834
 H   1.482613 -4.720172  1.772787
 H  -2.231180  1.271952  4.112212
 H  -1.609744 -1.358052  4.363018

TS C4-HAT formation
(d, Cpd I + Geraniol)
 C   4.21842 -1.761495  0.052300
 S   3.193658 -0.757331  1.176804
 H   3.779620 -2.747878 -0.117350
 H   5.211516 -1.870154  0.499105
 N   1.185336 -1.978769 -1.045246
 N   0.064120 -1.602248  1.568062
 N   2.008307  0.745807 -1.082587
 C   2.365051 -0.870823 -2.888836
 C   0.065787 -3.702335  0.291772
 C  -0.206064 -0.013817  3.416407
 C   1.984562  2.846865  0.181657
 C   1.748987 -1.970365 -2.294019
 C  -0.211708 -2.939307  1.422871
 C   0.353193  1.097956  2.799779
 C   2.299723  2.081718 -0.932096
 O  -0.507354  0.052951 -0.660821
 C   1.630861 -3.271475 -2.909327
 C  -0.807389 -3.464140  2.628746
 C   0.478480  2.399160  3.410147
 C   2.990012  2.576405 -2.101659
 C   0.983552 -4.068782 -2.014781
 C  -0.879305 -2.428916  3.511836
 C   1.089630  3.201819  2.495532
 C   3.105628  1.529337 -2.962951
 C   0.707355 -3.251550 -0.856395
 C  -0.330414 -1.275845  2.840778
 C   1.340870  2.389453  1.328638
 C   2.482540  0.393160 -2.322632
 N   0.880090  1.111873  1.529058
 Fe  0.938662 -0.407742  0.206551
 H   2.777802 -1.010750 -3.883262
 H  -0.231765 -4.746364  0.315500
 H  -0.568197  0.108339  4.432623
 H   2.286160  3.889745  0.167977
 C  -5.177124 -1.751113 -1.699916
 O  -6.467099 -1.286777 -2.123728
 C  -4.285780 -0.628214 -1.249371
 C  -3.669055 -0.467223 -0.052753
 C  -3.768221 -1.463002  1.080894
 C  -2.803883  0.703906  0.212422
 C  -3.034472  1.987396 -0.586228
 C  -2.328451  3.192445 -0.022591
 C  -1.652221  4.145660 -0.684865
 C  -1.416211  4.164292 -2.176068
 C  -1.070576  5.330819  0.050569
 H  -6.327872 -0.696172 -2.881469
 H  -4.688790 -2.304589 -2.519538
 H  -5.376360 -2.469572 -0.902523
 H  -4.133414  0.135490 -2.011936
 H  -2.788980 -1.906986  1.289324
 H  -4.091739 -0.961876  2.002077
 H  -4.465417 -2.278948  0.879262
 H  -2.677385  0.892389  1.284619
 H  -1.586631  0.328418 -0.104941
 H  -2.764420  1.829548 -1.634952
 H  -4.120448  2.185943 -0.583167
 H  -2.430508  3.310666  1.058151
 H  -1.832873  3.297041 -2.693002
 H  -1.853883  5.066855 -2.625666
 H  -0.339666  4.198970 -2.392806
 H  -1.505389  6.272397 -0.314541
 H  -1.245575  5.269987  1.129360
 H   0.013088  5.405130 -0.115579
 H   4.330175 -1.255259 -0.913053
 H   0.711955 -5.112133 -2.111662
 H   2.002364 -3.523339 -3.894371
 H  -1.117183 -4.492280  2.765373
 H  -1.260825 -2.429339  4.524797
 H   0.141973  2.645901  4.408958
 H   1.359702  4.245908  2.586899
 H   3.334530  3.594904 -2.227120
 H   3.562610  1.508640 -3.944030

TS C4-HAT formation
(q, Cpd I + Geraniol)
 C   3.134204 -2.989605  1.040403
 S   2.980613 -1.176208  1.176332
 H   2.288905 -3.485203  1.527860
 H   4.052604 -3.282460  1.558671
 N   0.921189 -2.066038 -1.099405
 N   0.040288 -1.475331  1.556520
 N   2.033752  0.531614 -1.337148
 C   2.081469 -1.190180 -3.080316
 C  -0.335728 -3.587799  0.372584
 C   0.060331  0.232272  3.321671
 C   2.461299  2.632587 -0.136404
 C   1.398699 -2.179455 -2.384135
 C  -0.429823 -2.759541  1.488333
 C   0.753166  1.221030  2.632436
 C   2.536071  1.808036 -1.249926
 O  -0.490045  0.178182 -0.569609
 C   1.069962 -3.475582 -2.924533
 C  -1.031541 -3.138527  2.745999
 C   1.097493  2.508814  3.184051
 C   3.194355  2.167886 -2.485223
 C   0.379061 -4.142173 -1.958859
 C  -0.921542 -2.061527  3.573196
 C   1.783014  3.180462  2.216905
 C   3.091764  1.093744 -3.313772
 C   0.285999 -3.257610 -0.823312
 C  -0.250410 -1.026059  2.823550
 C   1.849489  2.305832  1.070955
 C   2.371522  0.074493 -2.586113
 N   1.222864  1.119534  1.344935
 Fe  0.994668 -0.448513  0.069094
 H   2.405749 -1.421890 -4.089830
 H  -0.773316 -4.578292  0.448375
 H  -0.241817  0.454456  4.340345
 H   2.922673  3.612297 -0.211160
 C  -5.304754 -1.292845 -1.563112
 O  -6.540426 -0.706315 -1.996925
 C  -4.311116 -0.257988 -1.116766
 C  -3.706738 -0.130534  0.093652
 C  -3.933348 -1.080421  1.248205
 C  -2.745033  0.957170  0.353844
 C  -2.868098  2.264416 -0.434131
 C  -2.068212  3.401713  0.144866
 C  -1.289384  4.286841 -0.498972
 C  -1.014492  4.281156 -1.983462
 C  -0.616403  5.411388  0.253660
 H  -6.341464 -0.141067 -2.760737
 H  -4.868527 -1.897607 -2.375646
 H  -5.577563 -1.980545 -0.760661
 H  -4.066326  0.470718 -1.889390
 H  -2.985564 -1.518486  1.578484
 H  -4.347138 -0.538043  2.108416
 H  -4.612675 -1.900949  1.008988
 H  -2.582514  1.123706  1.424969
 H  -1.564954  0.483336  0.012011
 H  -2.610059  2.097206 -1.484312
 H  -3.934015  2.550894 -0.429243
 H  -2.185676  3.530743  1.222902
 H  -1.510331  3.465501 -2.514247
 H  -1.338165  5.227119 -2.440066
 H   0.064559  4.196242 -2.171890
 H  -0.944311  6.389994 -0.125312
 H  -0.82945  5.371618  1.326535
 H   0.473799  5.376060  0.122054
 H   3.181813 -3.309982 -0.002977
 H  -0.036599 -5.141129 -1.993334
 H   1.338515 -3.811656 -3.917832
 H  -1.474698 -4.105015  2.949460
 H  -1.255004 -1.960184  4.598074
 H   0.843682  2.836309  4.184116
 H   2.207783  4.175177  2.256893
 H   3.672162  3.121563 -2.670035
 H   3.466216  0.978879 -4.322915

TS C5-HAT formation
(d, Cpd I + Geraniol)
 C   0.829618 -3.167299  2.745891
 S   1.650247 -2.766128  1.159542
 H   0.869386 -2.325420  3.441941
 H   1.376849 -4.009283  3.183397
 N  -1.216681 -1.590941  0.737112
 N   0.699258  0.143027  1.909115
 N   0.501365 -1.948343 -1.524078
 C  -1.691661 -3.008582 -1.216339
 C  -1.505974 -0.394210  2.858632
 C   2.961500  1.087616  1.676877
 C   2.658928 -1.320745 -2.512950
 C  -2.007207 -2.452914  0.016782
 C  -0.269541  0.238421  2.882695
 C   3.269723  0.553332  0.432792
 C   1.437360 -1.979514 -2.527546
 O  -0.151255  0.414482 -0.733711
 C  -3.251454 -2.679283  0.712473
 C   0.193015  1.079388  3.957476
 C   4.504488  0.791882 -0.275553
 C   0.982247 -2.832110 -3.600442
 C  -3.209890 -1.931572  1.850343
 C   1.453938  1.481339  3.631624
 C   4.412184  0.125375 -1.459518
 C  -0.234611 -3.316861 -3.230240
 C  -1.936880 -1.252961  1.856284
 C   1.764775  0.895011  2.353035
 C   3.123663 -0.525742 -1.473131
 C  -0.530002 -2.757849 -1.932232
 N   2.447347 -0.252915 -0.311045
 Fe  0.572326 -0.845749  0.177810
 H  -2.421702 -3.674612 -1.664825
 H  -2.173500 -0.222992  3.696972
 H   3.709665  1.712302  2.154079
 H   3.306305 -1.446773 -3.374802
 C  -4.944764  0.869330 -1.150892
 O  -5.791000  1.131816 -2.277190
 C  -3.715964  1.718349 -1.286661
 C  -3.105917  2.453573 -0.342429
 C  -3.597084  2.593859  1.080428
 C  -1.850764  3.231424 -0.686321
 C  -0.596487  2.837498  0.123063
 C   0.625685  3.654185 -0.037336
 C   1.316582  3.921095 -1.174968
 C   0.980481  3.346517 -2.527068
 C   2.523738  4.821887 -1.140992
 H  -6.499416  0.469234 -2.265696
 H  -4.651277 -0.193962 -1.137668
 H  -5.476010  1.065898 -0.209195
 H  -3.275499  1.685386 -2.283616
 H  -3.517346  3.636464  1.414875
 H  -4.637429  2.279888  1.199021
 H  -2.995884  1.990581  1.772984
 H  -1.649806  3.158756 -1.758581
 H  -2.034295  4.298126 -0.475228
 H  -0.839843  2.731825  1.183948
 H  -0.330738  1.693234 -0.214633
 H   1.038579  4.058508  0.887356
 H   0.498251  4.101523 -3.164552
 H   1.898856  3.042985 -3.045131
 H   0.326181  2.475260 -2.462297
 H   3.424946  4.273650 -1.449514
 H   2.409140  5.652126 -1.851858
 H   2.698824  5.242723 -0.146281
 H  -0.887869 -3.989909 -3.770601
 H   1.539753 -3.025047 -4.508033
 H  -0.211219 -3.466233  2.598694
 H   5.138107  0.064535 -2.260262
 H   5.321217  1.394729  0.100356
 H   2.126980  2.115947  4.193786
 H  -0.383116  1.313610  4.843472
 H  -3.959350 -1.837096  2.625734
 H  -4.042519 -3.327273  0.357249

TS C5-HAT formation
(q, Cpd I + Geraniol)
 C   1.261461 -2.922474  2.881248
 S   2.198391 -2.156586  1.514343
 H   0.845778 -2.151657  3.537751
 H   1.962762 -3.530003  3.461812
 N  -0.868523 -1.762625  0.842047
 N   0.717030  0.422754  1.789117
 N   0.857676 -2.056374 -1.404727
 C  -1.136490 -3.415281 -0.952986
 C  -1.331416 -0.410728  2.846150
 C   2.754770  1.718687  1.367787
 C   2.932461 -1.263267 -2.450127
 C  -1.525864 -2.800093  0.232908
 C  -0.236760  0.445215  2.772832
 C   3.170149  1.068127  0.213364
 C   1.812792 -2.080784 -2.392994
 O  -0.172908  0.309827 -0.763453
 C  -2.708081 -3.150134  0.981970
 C   0.064048  1.481129  3.734358
 C   4.353977  1.418371 -0.532953
 C   1.504930 -3.111304 -3.357819
 C  -2.766986 -2.296917  2.043425
 C   1.215435  2.078617  3.318342
 C   4.409085  0.579127 -1.604431
 C   0.363279 -3.717986 -2.935387
 C  -1.619622 -1.428488  1.944684
 C   1.616806  1.408152  2.103215
 C   3.256676 -0.284153 -1.517087
 C  -0.031795 -3.059865 -1.711496
 N   2.521913  0.019354 -0.399165
 Fe  0.769537 -0.811317  0.152841
 H  -1.752566 -4.230992 -1.317897
 H  -2.016593 -0.274470  3.677131
 H   3.375899  2.532092  1.729531
 H   3.607836 -1.397191 -3.289166
 C  -5.147983 -0.095705 -0.824604
 O  -5.839572 -0.430169 -2.036207
 C  -3.956278  0.740224 -1.185751
 C  -3.588337  1.931335 -0.685938
 C  -4.361575  2.711260  0.351931
 C  -2.317017  2.597473 -1.180957
 C  -1.179100  2.592147 -0.138305
 C  -0.155394  3.641151 -0.110236
 C   0.440273  4.329053 -1.123820
 C   0.189704  4.114902 -2.593898
 C   1.451074  5.403608 -0.810792
 H  -6.501958 -1.102590 -1.811732
 H  -4.803202 -1.014644 -0.322162
 H  -5.818039  0.423559 -0.126063
 H  -3.320581  0.283385 -1.944679
 H  -4.600166  3.713881 -0.028539
 H  -5.296903  2.228987  0.644350
 H  -3.768544  2.859529  1.263812
 H  -1.991166  2.101547 -2.100742
 H  -2.542468  3.640824 -1.447967
 H  -1.558474  2.411399  0.871453
 H  -0.580518  1.472924 -0.346495
 H   0.208108  3.872854  0.890895
 H  -0.373951  4.956446 -3.022725
 H   1.145079  4.078444 -3.133708
 H  -0.353646  3.195446 -2.810606
 H   2.427662  5.163375 -1.254640
 H   1.145070  6.365993 -1.245500
 H   1.587286  5.543490  0.265837
 H  -0.174976 -4.539389 -3.390795
 H   2.102914 -3.329450 -4.233375
 H   0.449605 -3.552840  2.511583
 H   5.149008  0.537638 -2.393520
 H   5.040372  2.209021 -0.258042
 H   1.759656  2.892534  3.780230
 H  -0.534696  1.702005  4.608773
 H  -3.510325 -2.245344  2.828793
 H  -3.392644 -3.944590  0.713612

TS C8-HAT formation
(d, Cpd I + Geraniol)
 C   5.000364 -0.483150  1.444956
 S   3.777903  0.874437  1.320029
 H   4.644857 -1.291348  2.088661
 H   5.908638 -0.055982  1.883155
 N   2.868798 -0.387560 -1.334294
 N   2.113888 -1.786251  1.033201
 N   1.578660  1.965951 -0.341337
 C   2.826834  1.798563 -2.453564
 C   3.319627 -2.756387 -0.879108
 C   0.975036 -1.587850  3.200579
 C   0.178969  2.866174  1.460721
 C   3.176294  0.457474 -2.372361
 C   2.736540 -2.823499  0.379687
 C   0.537360 -0.277953  3.065362
 C   0.854619  2.968805  0.251346
 O   0.397000 -0.442098 -0.601702
 C   3.910880 -0.258468 -3.387910
 C   2.719723 -4.005939  1.204906
 C  -0.229568  0.430029  4.060200
 C   0.906403  4.162029 -0.558587
 C   4.034679 -1.543528 -2.954915
 C   2.081486 -3.672593  2.361558
 C  -0.472553  1.673532  3.560905
 C   1.667900  3.865435 -1.647860
 C   3.383523 -1.615781 -1.668635
 C   1.696224 -2.288265  2.241480
 C   0.154187  1.734143  2.263397
 C   2.076459  2.488170 -1.510817
 N   0.752719  0.530138  1.973203
 Fe  1.766228  0.057524  0.304972
 H   3.144319  2.345402 -3.335597
 H   3.775986 -3.661144 -1.267694
 H   0.721733 -2.109502  4.117806
 H  -0.346166  3.745755  1.818609
 C  -7.160631  2.447679 -0.589513
 O  -8.527732  2.595036 -0.179129
 C  -6.791776  1.014222 -0.857641
 C  -6.360775  0.459332 -2.003269
 C  -6.154562  1.213216 -3.296785
 C  -6.053181 -1.025347 -2.060168
 C  -4.560993 -1.377918 -2.314849
 C  -3.649879 -0.920298 -1.216185
 C  -2.706866 -1.636328 -0.543829
 C  -2.362459 -3.078946 -0.825555
 C  -1.907961 -0.979866  0.500264
 H  -8.645041  2.069312  0.628512
 H  -6.489822  2.864684  0.179775
 H  -7.056017  3.073291 -1.478522
 H  -6.900304  0.360385  0.011087
 H  -5.090300  1.272585 -3.560119
 H  -6.651997  0.691395 -4.125008
 H  -6.539536  2.234772 -3.263359
 H  -6.374174 -1.507051 -1.129246
 H  -6.638030 -1.477957 -2.874010
 H  -4.483364 -2.457220 -2.475493
 H  -4.244401 -0.906135 -3.257981
 H  -3.777699  0.121555 -0.921190
 H  -2.483048 -3.687168  0.080161
 H  -2.967923 -3.523057 -1.618038
 H  -1.308291 -3.160092 -1.121163
 H  -1.630087 -1.626188  1.339369
 H  -0.816554 -0.712017  0.030277
 H  -2.299705 -0.018685  0.842388
 H   5.246655 -0.887196  0.459549
 H   0.418673  5.095394 -0.308087
 H   1.933046  4.503097 -2.481434
 H   4.273749  0.182567 -4.307442
 H   4.522791 -2.377316 -3.443108
 H   3.148727 -4.958213  0.920364
 H   1.874549 -4.294344  3.223192
 H  -1.012986  2.492093  4.018868
 H  -0.532511  0.012492  5.011855

TS C8-HAT formation
(q, Cpd I + Geraniol)
 C   4.148078 -0.868149  2.835890
 S   3.647489  0.294873  1.521036
 H   3.362138 -0.948872  3.593215
 H   5.047798 -0.465002  3.310644
 N   2.052964 -2.390937  0.973657
 N   0.791908 -0.161793  2.251527
 N   2.614078 -0.793975 -1.302685
 C   3.254283 -3.146154 -1.030241
 C   0.899680 -2.439483  3.148937
 C   0.342021  2.242314  2.052063
 C   2.419212  1.461582 -2.257644
 C   2.704010 -3.345262  0.230816
 C   0.532368 -1.097949  3.216283
 C   0.879900  2.439023  0.785803
 C   2.795013  0.127282 -2.307885
 O   0.156413 -0.905323 -0.271966
 C   2.694228 -4.608930  0.926089
 C  -0.150354 -0.486311  4.333385
 C   0.866724  3.699310  0.083784
 C   3.488609 -0.481270 -3.419734
 C   2.010128 -4.414198  2.087789
 C  -0.290061  0.834460  4.031175
 C   1.439536  3.479680 -1.132763
 C   3.740564 -1.771254 -3.069161
 C   1.609900 -3.028596  2.111458
 C   0.300279  1.029056  2.727631
 C   1.807531  2.085146 -1.173792
 C   3.194414 -1.958255 -1.744949
 N   1.467224  1.471922  0.003242
 Fe  1.684924 -0.496627  0.440739
 H   3.744292 -3.991978 -1.501891
 H   0.625174 -3.074124  3.985723
 H  -0.090926  3.105361  2.548076
 H   2.633888  2.073171 -3.128222
 C  -5.310431  4.715756 -1.900542
 O  -6.494754  5.506798 -1.724847
 C  -5.588125  3.238789 -1.836380
 C  -5.379565  2.304759 -2.779928
 C  -4.798684  2.582680 -4.147194
 C  -5.739534  0.855885 -2.509363
 C  -4.532946 -0.123512 -2.464018
 C  -3.575181  0.170226 -1.348249
 C  -3.100016 -0.674787 -0.389092
 C  -3.445244 -2.143593 -0.299081
 C  -2.149685 -0.179519  0.603876
 H  -6.867594  5.279411 -0.857890
 H  -4.562000  4.984922 -1.136815
 H  -4.902435  5.016572 -2.867920
 H  -6.013961  2.910083 -0.885215
 H  -3.801453  2.136366 -4.255976
 H  -5.427990  2.133234 -4.926751
 H  -4.704436  3.648946 -4.364390
 H  -6.288019  0.780675 -1.563143
 H  -6.416334  0.504735 -3.301983
 H  -4.921699 -1.144130 -2.399338
 H  -3.995838 -0.063435 -3.423222
 H  -3.230555  1.203806 -1.303053
 H  -3.863460 -2.380272  0.687686
 H  -4.159736 -2.468921 -1.057908
 H  -2.537321 -2.750394 -0.410507
 H  -2.189937 -0.668552  1.580852
 H  -0.958174 -0.517846  0.209011
 H  -2.071689  0.906875  0.682290
 H   4.357812 -1.861659  2.432997
 H   3.748589  0.033697 -4.335767
 H   4.248128 -2.540157 -3.637448
 H   3.147183 -5.518374  0.552651
 H   1.788376 -5.130080  2.868982
 H  -0.466046 -1.012930  5.225021
 H  -0.747247  1.617969  4.621973
 H   1.604296  4.182956 -1.939101
 H   0.461881  4.620282  0.483496
```

**Table S9** Cartesian coordinates (xyz, in Å) of all DFT optimized structures in **Fig. S7** and **Table S8**

```
Nerol (2)
 O  -3.721164 -1.593448 -0.854277
 H  -4.069693 -2.477592 -0.657598
 C  -3.178687 -1.073848  0.370253
 H  -2.300118 -1.654353  0.680203
 H  -3.927118 -1.160081  1.175631
 C  -2.844596  0.372495  0.153983
 H  -3.693824  0.972084 -0.176657
 C  -1.660286  0.984028  0.316637
 C  -1.526547  2.467594  0.058720
 H  -0.747551  2.676381 -0.686475
 H  -2.463301  2.905012 -0.300354
 H  -1.229590  2.998275  0.974034
 C  -0.383571  0.285902  0.744931
 H   0.158499  0.928066  1.452299
 H  -0.600041 -0.644006  1.280506
 C   0.560148 -0.036431 -0.445139
 H   0.006743 -0.673909 -1.149835
 H   0.798675  0.884977 -0.985493
 C   1.808381 -0.751348 -0.000843
 H   1.626147 -1.728624  0.451920
 C   3.084729 -0.339783 -0.073678
 C   3.538667  0.977223 -0.656035
 H   4.227525  0.811594 -1.496150
 H   2.716578  1.600612 -1.014986
 H   4.098716  1.555997  0.091597
 C   4.206722 -1.210902  0.442623
 H   3.836015 -2.155196  0.854105
 H   4.924828 -1.445149 -0.356013
 H   4.777064 -0.696079  1.228774

TS C2-O formation
(d, conformer 1, Cpd I + Nerol)
 C  -3.971650  2.283952  0.182077
 S  -2.999423  1.340278  1.411622
 H  -3.368922  3.067161 -0.285692
 H  -4.797376  2.760738  0.721831
 N  -0.509567  2.155884 -0.222988
 N  -0.186493  0.535374  2.112171
 N  -2.121959  0.121697 -1.398673
 C  -1.478536  2.237778 -2.474277
 C   0.740634  2.788918  1.793812
 C  -0.717446 -1.632184  3.129377
 C  -3.147318 -2.081395 -1.038377
 C  -0.728136  2.767119 -1.432435
 C   0.531964  1.629109  2.528612
 C  -1.501525 -2.138863  2.100381
 C  -2.907484 -0.936203 -1.788298
 O   0.378919 -0.273981 -0.374466
 C  -0.081412  4.058511 -1.459960
 C   1.067165  1.396407  3.849336
 C  -2.182801 -3.410348  2.150384
 C  -3.435701 -0.702112 -3.109772
 C   0.529502  4.217473 -0.254077
 C   0.669527  0.148639  4.221429
 C  -2.874397 -3.532975  0.983594
 C  -2.951330  0.503732 -3.519590
 C   0.263373  3.021472  0.510986
 C  -0.113456 -0.381064  3.130159
 C  -2.613137 -2.338593  0.216964
 C  -2.134153  1.012821 -2.445488
 N  -1.766964 -1.506135  0.910925
 Fe -1.058287  0.278365  0.302970
 H  -1.575957  2.834867 -3.375461
 H   1.337980  3.570084  2.253234
 H  -0.589672 -2.250821  4.011972
 H  -3.798155 -2.835361 -1.469699
 C   1.114759 -1.283829 -2.773790
 O   1.633413 -2.121987 -3.817503
 C   0.868520 -2.053254 -1.496962
 C   1.862727 -2.559827 -0.688461
 C   1.521828 -3.475892  0.449536
 C   3.311123 -2.185500 -0.852669
 C   3.771806 -1.088197  0.155722
 C   5.234705 -0.774709  0.001428
 C   5.806601  0.382013 -0.373501
 C   5.057940  1.648087 -0.714503
 C   7.309772  0.499215 -0.479453
 H   0.974513 -2.813622 -3.991377
 H   0.186507 -0.793845 -3.089787
 H   1.865347 -0.506018 -2.622955
 H  -0.134021 -2.448146 -1.360700
 H   0.463587 -3.748653  0.457152
 H   2.125397 -4.392814  0.402459
 H   1.751912 -3.003286  1.415659
 H   3.930534 -3.079267 -0.694413
 H   3.510110 -1.838006 -1.869540
 H   3.141949 -0.205451  0.021573
 H   3.583801 -1.454144  1.175000
 H   5.899905 -1.614970  0.212656
 H   3.973419  1.548543 -0.630437
 H   5.373075  2.470731 -0.057343
 H   5.289841  1.967303 -1.740196
 H   7.693030  1.283985  0.188127
 H   7.813024 -0.439345 -0.225712
 H   7.611929  0.785034 -1.497021
 H  -4.386301  1.635693 -0.593837
 H   1.112155  5.056885  0.103291
 H  -0.107923  4.740713 -2.300051
 H   1.668179  2.107375  4.401722
 H   0.872823 -0.379224  5.144433
 H  -2.134134 -4.100173  2.983237
 H  -3.511630 -4.344796  0.656883
 H  -4.088408 -1.383920 -3.639875
 H  -3.124840  1.019014 -4.455692

TS C2-O formation
(q, conformer 1, Cpd I + Nerol)
 C  -4.316532  1.625798  0.094928
 S  -3.123386  1.073396  1.363359
 H  -3.927746  2.463261 -0.489003
 H  -5.228667  1.936605  0.614684
 N  -1.439727  1.474541 -1.307303
 N  -0.033934  1.862275  1.136195
 N  -2.225760 -1.137477 -0.460902
 C  -2.735479 -0.138804 -2.640708
 C  -0.207319  3.504045 -0.679292
 C   0.515570  0.854814  3.308588
 C  -2.174464 -2.709406  1.424671
 C  -2.110603  1.086005 -2.445140
 C   0.237463  3.082057  0.565603
 C  -0.181103 -0.339783  3.152697
 C  -2.547917 -2.328916  0.140993
 O   0.362013 -0.307919 -0.341556
 C  -2.084101  2.148199 -3.421548
 C   1.052112  3.877554  1.455027
 C  -0.248900 -1.383953  4.147372
 C  -3.337097 -3.139443 -0.757322
 C  -1.387862  3.178338 -2.865722
 C   1.264578  3.127503  2.569938
 C  -1.013662 -2.380016  3.619666
 C  -3.489256 -2.421219 -1.904727
 C  -0.980879  2.745430 -1.550688
 C   0.576482  1.872949  2.368375
 C  -1.411308 -1.947359  2.301137
 C  -2.795244 -1.169456 -1.706520
 N  -0.898856 -0.698300  2.042163
 Fe -1.073562  0.358570  0.320581
 H  -3.230388 -0.298665 -3.593574
 H   0.082770  4.497990 -1.004904
 H   1.042524  1.007183  4.245208
 H  -2.505108 -3.683515  1.771578
 C   1.139632 -1.141927 -2.703980
 O   1.627325 -1.975239 -3.765361
 C   0.880315 -1.929248 -1.438534
 C   1.891526 -2.512292 -0.685830
 C   1.561807 -3.494341  0.399610
 C   3.343854 -2.157405 -0.860281
 C   3.827089 -1.060525  0.137174
 C   5.305617 -0.811018  0.015207
 C   5.936633  0.311959 -0.366891
 C   5.252417  1.601059 -0.753936
 C   7.445856  0.363018 -0.432533
 H   0.942782 -2.636964 -3.955601
 H   0.224197 -0.620463 -3.004230
 H   1.914097 -0.389991 -2.540964
 H  -0.111001 -2.363341 -1.347047
 H   0.484539 -3.639061  0.513994
 H   2.025413 -4.469379  0.188312
 H   1.967658 -3.166870  1.367014
 H   3.951526 -3.058661 -0.698983
 H   3.543324 -1.820587 -1.881134
 H   3.239884 -0.153571 -0.026758
 H   3.599348 -1.397955  1.158315
 H   5.928334 -1.673852  0.262175
 H   4.162582  1.546919 -0.703589
 H   5.579499  2.422401 -0.101002
 H   5.529814  1.891449 -1.776885
 H   7.843990  1.145486  0.229049
 H   7.900960 -0.590142 -0.144491
 H   7.788328  0.612763 -1.446827
 H  -4.568771  0.803337 -0.581303
 H  -1.155347  4.145441 -3.292960
 H  -2.544497  2.093983 -4.399900
 H   1.401674  4.878494  1.236296
 H   1.824170  3.383730  3.460542
 H   0.231837 -1.341960  5.116457
 H  -1.290217 -3.327466  4.064497
 H  -3.719107 -4.124547 -0.521345
 H  -4.023729 -2.692909 -2.806201

TS C2-O formation
(d, conformer 2, Cpd I + Nerol)
 C   3.664136  1.898083 -1.924978
 S   2.536853  0.504690 -2.289661
 H   3.109367  2.818348 -1.724568
 H   4.277459  2.051871 -2.819918
 N   0.377253  1.986556 -0.652384
 N  -0.222774 -0.561378 -1.816791
 N   2.469117  0.789681  0.865393
 C   1.842650  3.167061  0.924018
 C  -1.376685  1.553049 -2.314911
 C   0.398714 -2.933987 -1.871175
 C   3.641991 -1.314433  1.344265
 C   0.794471  3.111919  0.014090
 C  -1.161077  0.191244 -2.481541
 C   1.413625 -2.883777 -0.925218
 C   3.451857  0.053693  1.485764
 O  -0.107419 -0.057749  0.876578
 C  -0.003529  4.248071 -0.384839
 C  -1.908871 -0.643207 -3.390530
 C   2.219742 -4.015699 -0.536602
 C   4.250788  0.906763  2.329132
 C  -0.905308  3.794187 -1.297794
 C  -1.409780 -1.905067 -3.269764
 C   3.128297 -3.560121  0.369444
 C   3.737857  2.164297  2.219085
 C  -0.661774  2.379758 -1.459059
 C  -0.358252 -1.844216 -2.283799
 C   2.880873 -2.148365  0.536086
 C   2.626696  2.085458  1.304085
 N   1.820124 -1.758640 -0.247524
 Fe  1.036602  0.089554 -0.380392
 H   2.073857  4.131569  1.364591
 H  -2.166732  2.008009 -2.903864
 H   0.188120 -3.894447 -2.330463
 H   4.456266 -1.766216  1.901599
 C  -0.571757 -0.505346  3.446225
 O   0.728391  0.054422  3.471044
 C  -0.733107 -1.485459  2.312321
 C  -1.940991 -1.898749  1.778474
 C  -1.997927 -3.080667  0.858119
 C  -3.245006 -1.196989  2.050892
 C  -3.591123 -0.131889  0.963399
 C  -4.873522  0.587366  1.281307
 C  -6.043342  0.544659  0.621643
 C  -6.304905 -0.263066 -0.626990
 C  -7.230650  1.344584  1.106271
 H   0.876548  0.337956  2.546951
 H  -1.340757  0.277897  3.394523
 H  -0.706928 -1.033638  4.400728
 H   0.137711 -2.110212  2.130045
 H  -2.621287 -3.871875  1.299770
 H  -2.469614 -2.816337 -0.098332
 H  -1.008473 -3.492965  0.649534
 H  -4.054546 -1.937103  2.075419
 H  -3.235459 -0.704468  3.028690
 H  -2.759271  0.581716  0.921255
 H  -3.631040 -0.619267 -0.015339
 H  -4.828634  1.209474  2.177730
 H  -6.615180  0.392142 -1.452911
 H  -5.439113 -0.838617 -0.962573
 H  -7.133982 -0.965536 -0.463246
 H  -7.571383  2.051494  0.336545
 H  -8.085791  0.688875  1.323482
 H  -6.998778  1.913692  2.012225
 H   4.323241  1.680140 -1.081079
 H   0.128535  5.253708 -0.006528
 H  -1.670072  4.348886 -1.826287
 H  -2.705628 -0.289943 -4.032550
 H  -1.710734 -2.804278 -3.791930
 H   2.097488 -5.018261 -0.926242
 H   3.908997 -4.109694  0.879621
 H   5.088138  0.568649  2.925921
 H   4.068024  3.073120  2.705635

TS C2-O formation
(q, conformer 2, Cpd I + Nerol)
 C   4.119854  0.101975 -1.954462
 S   2.362938  0.364798 -2.383675
 H   4.483850  0.855732 -1.252486
 H   4.694624  0.166037 -2.884184
 N   0.452306  2.133097 -0.572793
 N  -0.418353 -0.295721 -1.772303
 N   2.548134  0.735224  0.750293
 C   2.203179  3.166769  0.805592
 C  -1.476617  1.896487 -2.075504
 C   0.023836 -2.710734 -1.939869
 C   3.479400 -1.493155  1.225815
 C   1.059625  3.214092  0.021432
 C  -1.367000  0.530660 -2.315631
 C   1.081963 -2.787286 -1.041859
 C   3.450010 -0.111768  1.358513
 O  -0.069504 -0.020283  0.915783
 C   0.338985  4.425235 -0.296357
 C  -2.241208 -0.221007 -3.183755
 C   1.771535 -4.007090 -0.689481
 C   4.384442  0.650085  2.148020
 C  -0.704835  4.065450 -1.091017
 C  -1.807214 -1.512503 -3.156979
 C   2.727662 -3.669478  0.220195
 C   4.042916  1.962343  2.011297
 C  -0.621507  2.634192 -1.270308
 C  -0.672012 -1.553894 -2.266704
 C   2.629468 -2.240914  0.415180
 C   2.890760  2.006871  1.145971
 N   1.617238 -1.729290 -0.352742
 Fe  0.998258  0.213339 -0.424297
 H   2.579188  4.104953  1.200842
 H  -2.279904  2.429662 -2.573813
 H  -0.295449 -3.633573 -2.413955
 H   4.246916 -2.029905  1.774402
 C  -0.462482 -0.258194  3.451793
 O   0.809306  0.362770  3.450326
 C  -0.577352 -1.303852  2.368273
 C  -1.786137 -1.853952  1.945508
 C  -1.805421 -3.122924  1.148334
 C  -3.116949 -1.194804  2.196816
 C  -3.498097 -0.169440  1.084241
 C  -4.837476  0.462835  1.346343
 C  -5.952389  0.405807  0.598504
 C  -6.079815 -0.328097 -0.715117
 C  -7.213420  1.112212  1.040789
 H   0.953261  0.595053  2.512117
 H  -1.267771  0.483244  3.355691
 H  -0.578080 -0.743948  4.430734
 H   0.314644 -1.911003  2.238632
 H  -2.433611 -3.875543  1.646227
 H  -2.249890 -2.959952  0.155572
 H  -0.806099 -3.540187  1.005427
 H  -3.898243 -1.963434  2.239304
 H  -3.129222 -0.678208  3.163364
 H  -2.716000  0.599719  1.057574
 H  -3.469611 -0.670731  0.112267
 H  -4.890918  1.030650  2.277913
 H  -6.385732  0.362740 -1.513170
 H  -5.156251 -0.817266 -1.033118
 H  -6.865427 -1.094239 -0.653820
 H  -7.533421  1.853607  0.294938
 H  -8.046210  0.402577  1.147490
 H  -7.080291  1.627541  1.997431
 H   4.269238 -0.892628 -1.524164
 H   0.615311  5.412201  0.052103
 H  -1.464659  4.695365 -1.535724
 H  -3.070818  0.201963 -3.735738
 H  -2.208588 -2.370802 -3.680681
 H   1.541278 -4.983467 -1.096531
 H   3.448329 -4.309456  0.713120
 H   5.192010  0.219818  2.726422
 H   4.509939  2.831326  2.456831

TS C1-HAT formation
(d, Cpd I + Nerol)
 C   4.63920 -1.006724  0.206391
 S   3.621168  0.222432  1.102265
 H   4.242221 -2.019080  0.317897
 H   5.642034 -0.974297  0.646257
 N   2.137301  0.227568 -1.653405
 N   1.376718 -1.816200  0.160825
 N   1.471475  2.206187  0.305854
 C   2.365933  2.658798 -1.935372
 C   2.147913 -2.175679 -2.145216
 C   0.588064 -2.277152  2.445718
 C   0.608976  2.560672  2.576202
 C   2.472516  1.346563 -2.377301
 C   1.711788 -2.621373 -0.903288
 C   0.452465 -0.964716  2.879443
 C   1.082902  3.008094  1.350205
 O  -0.244866  0.351269 -0.360738
 C   2.929101  0.967021 -3.692718
 C   1.558475 -4.008695 -0.542751
 C  -0.016399 -0.582054  4.190495
 C   1.262206  4.397774  1.001652
 C   2.852871 -0.390887 -3.760489
 C   1.134720 -4.035547  0.752468
 C  -0.017546  0.778680  4.223796
 C   1.765187  4.424749 -0.262739
 C   2.356829 -0.843962 -2.483186
 C   1.019596 -2.664887  1.183984
 C   0.453450  1.227358  2.934414
 C   1.891175  3.051591 -0.691100
 N   0.735231  0.150296  2.133564
 Fe  1.340025  0.188038  0.200047
 H   2.666284  3.441989 -2.624176
 H   2.369642 -2.926002 -2.897329
 H   0.331472 -3.064451  3.147376
 H   0.346639  3.311269  3.314892
 C  -1.685819 -1.454244 -1.636839
 O  -2.530084 -0.602552 -2.360174
 C  -2.350487 -2.443420 -0.777317
 C  -3.625151 -2.500961 -0.312152
 C  -4.030698 -3.621033  0.611614
 C  -4.705948 -1.487534 -0.615411
 C  -4.598766 -0.227880  0.292702
 C  -5.745868  0.726369  0.086263
 C  -5.742470  1.884645 -0.594466
 C  -4.538021  2.457985 -1.303687
 C  -6.992138  2.728546 -0.693129
 H  -1.969995  0.028779 -2.840299
 H  -0.995056 -0.773917 -0.927461
 H  -0.920783 -1.925259 -2.277400
 H  -1.659042 -3.212318 -0.433754
 H  -4.824363 -4.230577  0.157246
 H  -3.193884 -4.278070  0.866822
 H  -4.450400 -3.218363  1.544405
 H  -4.651436 -1.161465 -1.655116
 H  -5.688637 -1.951368 -0.460723
 H  -4.587959 -0.552624  1.342790
 H  -3.635236  0.253264  0.104226
 H  -6.689061  0.412664  0.538306
 H  -4.798983  2.728443 -2.336205
 H  -4.202532  3.385657 -0.818010
 H  -3.693001  1.767246 -1.344136
 H  -6.821200  3.735251 -0.285552
 H  -7.294300  2.867532 -1.740919
 H  -7.832533  2.281577 -0.152041
 H   2.032580  5.282508 -0.866431
 H   1.030705  5.228585  1.655746
 H   4.720090 -0.766053 -0.857016
 H  -0.303906  1.435635  5.035024
 H  -0.302337 -1.278700  4.968042
 H   0.916623 -4.894410  1.374167
 H   1.761157 -4.840596 -1.205009
 H   3.108299 -1.042496 -4.586277
 H   3.257753  1.664497 -4.452491

TS C1-HAT formation
(q, Cpd I + Nerol)
 C   4.710051 -0.401406  0.183978
 S   3.384139 -0.002890  1.373421
 H   4.513213 -1.365532 -0.295681
 H   5.646358 -0.479163  0.745296
 N   2.191759  0.486081 -1.518108
 N   1.436582 -1.813078  0.011077
 N   1.184650  2.193283  0.529165
 C   2.182079  2.939472 -1.587128
 C   2.445288 -1.859675 -2.222118
 C   0.493208 -2.541980  2.152744
 C   0.227757  2.256994  2.788365
 C   2.453422  1.685976 -2.127818
 C   1.929687 -2.473083 -1.083983
 C   0.280897 -1.289403  2.713532
 C   0.699251  2.857876  1.629755
 O  -0.284892  0.206599 -0.463894
 C   3.023862  1.467862 -3.434621
 C   1.838084 -3.901962 -0.887549
 C  -0.291989 -1.074141  4.019869
 C   0.779187  4.285980  1.425167
 C   3.086425  0.118637 -3.615333
 C   1.285975 -4.093047  0.343171
 C  -0.362492  0.273447  4.204271
 C   1.329168  4.477449  0.196070
 C   2.558288 -0.487638 -2.417755
 C   1.040946 -2.781464  0.897809
 C   0.163426  0.885507  3.008657
 C   1.587750  3.167193 -0.355135
 N   0.561812 -0.079563  2.118581
 Fe  1.278486  0.218666  0.261577
 H   2.449531  3.807454 -2.181533
 H   2.790833 -2.505182 -3.023674
 H   0.216620 -3.405494  2.749558
 H  -0.122217  2.907694  3.583415
 C  -1.537773 -1.627280 -1.750174
 O  -2.311151 -0.754542 -2.523367
 C  -2.269810 -2.586807 -0.922056
 C  -3.551911 -2.570841 -0.472860
 C  -4.040826 -3.674714  0.430798
 C  -4.559699 -1.482125 -0.771317
 C  -4.378692 -0.247414  0.158151
 C  -5.446872  0.793382 -0.054462
 C  -5.344053  1.955018 -0.721476
 C  -4.087053  2.442598 -1.403239
 C  -6.524492  2.892072 -0.832467
 H  -1.707206 -0.114457 -2.934061
 H  -0.862330 -0.897243 -0.980086
 H  -0.732152 -2.102552 -2.330398
 H  -1.632559 -3.401324 -0.579440
 H  -4.867493 -4.225964 -0.039289
 H  -3.251302 -4.389070  0.683709
 H  -4.442427 -3.260440  1.366617
 H  -4.471547 -1.144263 -1.804936
 H  -5.573720 -1.880630 -0.635841
 H  -4.413148 -0.587109  1.203135
 H  -3.378206  0.163459 -0.002948
 H  -6.419186  0.547475  0.377662
 H  -4.305276  2.730024 -2.441207
 H  -3.698893  3.345032 -0.909224
 H  -3.290886  1.695246 -1.424933
 H  -6.286419  3.878387 -0.408881
 H  -6.795374  3.065506 -1.883712
 H  -7.406437  2.503616 -0.312661
 H   1.553979  5.409315 -0.306833
 H   0.457095  5.027310  2.145201
 H   4.806429  0.367227 -0.586106
 H  -0.738961  0.820215  5.059256
 H  -0.597428 -1.864998  4.692941
 H   1.064806 -5.025350  0.847029
 H   2.162432 -4.644246 -1.605689
 H   3.450784 -0.432173 -4.473086
 H   3.324422  2.255748 -4.113316

TS C4-HAT formation
(d, Cpd I + Nerol)
 C   2.190659 -2.952305  2.228085
 S   1.157421 -3.096145  0.723869
 H   3.102278 -2.379929  2.040536
 H   2.468735 -3.971046  2.518749
 N   2.113001 -0.115031  0.724500
 N   1.189302 -1.283822 -1.718723
 N  -0.394166 -0.655841  1.929924
 C   1.468077  0.546246  2.997557
 C   3.430687 -0.359050 -1.340314
 C  -0.725356 -2.357155 -2.817886
 C  -2.563749 -1.782284  1.626891
 C   2.360543  0.481627  1.936992
 C   2.459525 -0.961202 -2.126935
 C  -1.583950 -2.380618 -1.726480
 C  -1.622211 -1.092699  2.377600
 O  -0.165636  0.577131 -0.583361
 C   3.699776  1.015456  1.961532
 C   2.643264 -1.335009 -3.509361
 C  -2.910725 -2.948911 -1.747749
 C  -1.808266 -0.706644  3.753019
 C   4.256807  0.737393  0.749923
 C   1.469298 -1.881252 -3.929168
 C  -3.422466 -2.804403 -0.493599
 C  -0.687169 -0.029265  4.131717
 C   3.255425  0.042745 -0.022397
 C   0.566427 -1.850488 -2.802565
 C  -2.412831 -2.139309  0.293567
 C   0.195804 -0.010588  2.993979
 N  -1.302720 -1.894458 -0.474040
 Fe  0.376013 -0.921518  0.107076
 H   1.799336  1.037703  3.906488
 H   4.395631 -0.164479 -1.797138
 H  -1.086473 -2.783264 -3.748253
 H  -3.494606 -2.049835  2.116217
 C  -3.193624  1.423645 -1.921666
 O  -3.820485  2.277038 -2.891853
 C  -3.716716  1.685215 -0.537840
 C  -3.017971  2.076436  0.550927
 C  -3.712001  2.231653  1.887676
 C  -1.558432  2.393093  0.556745
 C  -1.106513  3.628788 -0.242976
 C   0.310108  4.043862  0.047135
 C   1.342552  4.156074 -0.804840
 C   1.281112  3.868073 -2.285150
 C   2.699768  4.601310 -0.311191
 H  -3.649767  3.192330 -2.616587
 H  -2.101625  1.497080 -1.966215
 H  -3.459462  0.405468 -2.229411
 H  -4.790683  1.531546 -0.428648
 H  -3.261370  1.569205  2.637557
 H  -3.604210  3.256936  2.266654
 H  -4.779513  1.999869  1.820196
 H  -1.165935  2.433046  1.579042
 H  -0.925333  1.434283  0.074846
 H  -1.277883  3.473276 -1.311745
 H  -1.779889  4.451711  0.052657
 H   0.497616  4.282039  1.096051
 H   0.293887  3.549286 -2.626304
 H   1.569741  4.756795 -2.863631
 H   1.996926  3.077780 -2.549134
 H   3.037963  5.501850 -0.843065
 H   2.693520  4.820061  0.761428
 H   3.456532  3.826482 -0.495576
 H   1.636037 -2.493407  3.050510
 H   5.249037  0.978005  0.390209
 H   4.139236  1.530175  2.806348
 H   3.559136 -1.188132 -4.067346
 H   1.219171 -2.280348 -4.903804
 H  -3.370356 -3.399395 -2.618179
 H  -4.391634 -3.107744 -0.118847
 H  -2.689840 -0.934245  4.338553
 H  -0.457328  0.412791  5.092693

TS C4-HAT formation
(q, Cpd I + Nerol)
 C   2.752034 -2.426204  2.257646
 S   1.392710 -2.845238  1.115136
 H   3.565714 -1.910558  1.742962
 H   3.120828 -3.362648  2.687457
 N   2.191226  0.152227  0.504046
 N   1.218295 -1.332671 -1.715317
 N  -0.177796 -0.474449  1.982423
 C   1.642395  0.970693  2.760291
 C   3.361341 -0.139775 -1.634251
 C  -0.600072 -2.778446 -2.510590
 C  -2.244695 -1.794613  1.942700
 C   2.455665  0.887127  1.637891
 C   2.409032 -0.923101 -2.268249
 C  -1.379352 -2.763268 -1.358404
 C  -1.335211 -0.941034  2.554792
 O  -0.288534  0.522115 -0.548708
 C   3.723269  1.561828  1.505376
 C   2.540323 -1.441764 -3.610233
 C  -2.635632 -3.456764 -1.212675
 C  -1.473094 -0.414936  3.893149
 C   4.216680  1.237399  0.277246
 C   1.423025 -2.176201 -3.858882
 C  -3.093461 -3.182134  0.040827
 C  -0.384786  0.372599  4.120036
 C   3.247344  0.368674 -0.344209
 C   0.606471 -2.112112 -2.668356
 C  -2.119358 -2.319408  0.662604
 C   0.420644  0.322874  2.921785
 N  -1.077071 -2.084958 -0.205236
 Fe  0.516829 -0.882134  0.099199
 H   1.994817  1.582983  3.584525
 H   4.258490  0.108713 -2.192291
 H  -0.961988 -3.363354 -3.350226
 H  -3.125819 -2.075755  2.510870
 C  -3.302113  1.034893 -2.113516
 O  -3.860768  1.966258 -3.057933
 C  -3.931938  1.168091 -0.756104
 C  -3.346866  1.627823  0.375645
 C  -4.124260  1.640159  1.676032
 C  -1.960218  2.154833  0.461270
 C  -1.605790  3.388107 -0.383500
 C  -0.268567  3.988275 -0.041094
 C   0.770724  4.239583 -0.853800
 C   0.808746  3.937010 -2.332209
 C   2.030622  4.877905 -0.315535
 H  -3.768046  2.850385 -2.667591
 H  -2.211984  1.125339 -2.060803
 H  -3.525469  0.046440 -2.530556
 H  -4.976926  0.862137 -0.703392
 H  -3.595301  1.068128  2.448036
 H  -4.227786  2.664638  2.058460
 H  -5.125645  1.214484  1.557021
 H  -1.639153  2.285809  1.499996
 H  -1.113759  1.214322  0.040698
 H  -1.683393  3.160913 -1.450363
 H  -2.386339  4.143531 -0.181899
 H  -0.159259  4.260106  1.010863
 H  -0.094110  3.443496 -2.698490
 H   0.951389  4.859411 -2.912703
 H   1.662071  3.285961 -2.565713
 H   2.262493  5.808906 -0.852062
 H   1.947203  5.110851  0.751020
 H   2.894958  4.213320 -0.451492
 H   2.383870 -1.793024  3.071052
 H   5.143969  1.552596 -0.183756
 H   4.161773  2.198498  2.263113
 H   3.388132 -1.258825 -4.257994
 H   1.159625 -2.725154 -4.753912
 H  -3.092923 -4.067152 -1.980839
 H  -4.005941 -3.519557  0.515470
 H  -2.297910 -0.635908  4.558630
 H  -0.130068  0.933566  5.010157

TS C5-HAT formation
(d, Cpd I + Nerol)
 C   1.033249  3.183216  2.703866
 S   0.335356  3.228254  1.012243
 H   0.407517  2.595163  3.380199
 H   1.062515  4.217238  3.063983
 N   1.941719  0.551609  0.981138
 N  -0.743546  0.399594  1.882685
 N   1.040717  1.671775 -1.493426
 C   3.382600  1.259296 -0.882727
 C   1.221592 -0.420886  3.114425
 C  -3.075290  0.986436  1.353194
 C  -0.940315  2.355537 -2.772007
 C   3.164851  0.723279  0.380052
 C  -0.139453 -0.182588  2.974370
 C  -2.865553  1.484660  0.073998
 C   0.425852  2.163867 -2.617776
 O   0.055524 -0.547465 -0.612608
 C   4.211234  0.221478  1.238151
 C  -1.126073 -0.486023  3.979697
 C  -3.910855  1.966313 -0.797006
 C   1.414987  2.493886 -3.616653
 C   3.605392 -0.271842  2.353364
 C  -2.329058 -0.069325  3.492946
 C  -3.310980  2.333591 -1.963381
 C   2.631161  2.201429 -3.079412
 C   2.187741 -0.063690  2.183850
 C  -2.085172  0.479735  2.183524
 C  -1.897970  2.084644 -1.802964
 C   2.388872  1.685179 -1.752975
 N  -1.650693  1.568709 -0.555640
 Fe  0.137855  0.970197  0.183869
 H   4.410449  1.330472 -1.223729
 H   1.555876 -0.896190  4.030921
 H  -4.093438  0.979637  1.728699
 H  -1.285151  2.764782 -3.716137
 C   2.141391 -2.829558 -2.072964
 O   3.047299 -3.508440 -2.954845
 C   2.107323 -3.585075 -0.778688
 C   1.037443 -3.987947 -0.074045
 C   1.219937 -4.726160  1.233669
 C  -0.400946 -3.775458 -0.501723
 C  -1.138648 -2.674383  0.294432
 C  -2.605421 -2.564884  0.152478
 C  -3.331387 -2.377504 -0.979394
 C  -2.732633 -2.161760 -2.345121
 C  -4.836512 -2.341455 -0.924245
 H   3.183654 -2.931424 -3.722675
 H   1.144470 -2.744291 -2.524804
 H   2.494166 -1.803074 -1.884335
 H   3.098846 -3.806832 -0.381414
 H   2.278425 -4.860858  1.476790
 H   0.749814 -4.193279  2.070874
 H   0.748158 -5.717818  1.191457
 H  -0.467921 -3.572254 -1.572704
 H  -0.951896 -4.713670 -0.332501
 H  -0.864478 -2.715723  1.352618
 H  -0.637143 -1.620623 -0.072407
 H  -3.165413 -2.602588  1.087284
 H  -1.675662 -1.892889 -2.302280
 H  -2.841396 -3.061245 -2.968002
 H  -3.265883 -1.356083 -2.865349
 H  -5.212912 -1.362957 -1.253947
 H  -5.268356 -3.084889 -1.608759
 H  -5.218899 -2.538080  0.081880
 H   3.611993  2.316434 -3.522756
 H   1.187223  2.902363 -4.592926
 H   2.049898  2.782237  2.711525
 H  -3.763883  2.742603 -2.857512
 H  -4.959702  2.008436 -0.532542
 H  -3.302680 -0.121752  3.963128
 H  -0.906493 -0.948555  4.933443
 H   4.058314 -0.735076  3.220563
 H   5.265942  0.244983  0.995642

TS C5-HAT formation
(q, Cpd I + Nerol)
 C  -1.003024 -3.078092  2.801350
 S   0.384629 -2.918053  1.626165
 H  -1.045862 -2.206314  3.461681
 H  -0.816524 -3.966607  3.412580
 N  -1.893530 -0.979292  0.594641
 N   0.415202  0.043750  1.941278
 N  -0.236479 -2.108654 -1.420575
 C  -2.686116 -2.248423 -1.351931
 C  -1.908951  0.415324  2.623447
 C   2.867785  0.066577  1.926870
 C   2.087805 -2.491037 -2.116056
 C  -2.885435 -1.524840 -0.180247
 C  -0.533940  0.568021  2.777558
 C   3.066888 -0.689050  0.778384
 C   0.715112 -2.609136 -2.277866
 O   0.023263  0.447206 -0.702645
 C  -4.180241 -1.208798  0.372170
 C   0.098179  1.290756  3.857565
 C   4.363191 -1.001971  0.228200
 C   0.081356 -3.321605 -3.363423
 C  -3.959514 -0.448186  1.480709
 C   1.442676  1.187773  3.663033
 C   4.146841 -1.718389 -0.910172
 C  -1.259845 -3.260938 -3.144997
 C  -2.530417 -0.303139  1.610550
 C   1.631585  0.403754  2.464663
 C   2.717192 -1.844266 -1.057336
 C  -1.451211 -2.508170 -1.927009
 N   2.078265 -1.223376 -0.015357
 Fe  0.081350 -1.018244  0.222663
 H  -3.566371 -2.626490 -1.862173
 H  -2.549178  0.890668  3.360019
 H   3.751771  0.423245  2.446250
 H   2.721300 -2.943117 -2.872614
 C  -2.744841  2.083916 -2.169729
 O  -3.743968  2.574131 -3.077378
 C  -2.809187  2.911421 -0.921298
 C  -1.817582  3.578798 -0.308692
 C  -2.093208  4.354883  0.960037
 C  -0.390468  3.649550 -0.816548
 C   0.618043  2.843258  0.024716
 C   2.060350  3.049243 -0.166156
 C   2.780088  3.093679 -1.319295
 C   2.205845  2.858761 -2.694530
 C   4.261008  3.370259 -1.280416
 H  -3.802461  1.937287 -3.807067
 H  -1.748924  2.110906 -2.629518
 H  -2.951053  1.032032 -1.916492
 H  -3.805218  2.950466 -0.477893
 H  -3.141983  4.274685  1.262535
 H  -1.475546  4.001670  1.796161
 H  -1.855388  5.419336  0.824840
 H  -0.339097  3.349378 -1.864476
 H  -0.070750  4.703794 -0.786638
 H   0.365193  2.878702  1.087141
 H   0.368816  1.603121 -0.256538
 H   2.633459  3.137736  0.757219
 H   1.303651  2.243426 -2.667873
 H   1.959856  3.807636 -3.192556
 H   2.941568  2.352734 -3.331150
 H   4.827302  2.520469 -1.687399
 H   4.512727  4.236643 -1.908599
 H   4.618846  3.568650 -0.265492
 H  -2.061594 -3.684565 -3.736274
 H   0.613806 -3.806455 -4.171693
 H  -1.959391 -3.182125  2.284041
 H   4.873815 -2.127225 -1.600256
 H   5.305331 -0.699846  0.667432
 H   2.245518  1.592353  4.266134
 H  -0.433047  1.796257  4.653957
 H  -4.684434 -0.014537  2.157704
 H  -5.123654 -1.528024 -0.051902

TS C8-HAT formation
(d, Cpd I + Nerol)
 C   4.900079 -1.146388  0.958429
 S   3.762499  0.164819  1.537871
 H   4.476611 -2.143838  1.100989
 H   5.823409 -1.062787  1.541216
 N   2.869843  0.659389 -1.333799
 N   2.007173 -1.839461 -0.232759
 N   1.564241  2.026997  0.798602
 C   2.862385  3.088104 -0.997891
 C   3.303422 -1.547987 -2.303874
 C   0.785283 -2.909731  1.606804
 C   0.117432  1.744283  2.758413
 C   3.213153  1.939452 -1.694387
 C   2.663571 -2.320976 -1.340338
 C   0.357139 -1.752449  2.245302
 C   0.835827  2.516880  1.852141
 O   0.351915  0.265753 -0.828875
 C   3.988129  1.923015 -2.914205
 C   2.619664 -3.762388 -1.357519
 C  -0.448427 -1.730579  3.442554
 C   0.917911  3.957351  1.889936
 C   4.103325  0.619956 -3.288638
 C   1.930914 -4.149466 -0.247302
 C  -0.652807 -0.420821  3.752821
 C   1.699458  4.332766  0.839574
 C   3.402036 -0.161514 -2.294788
 C   1.545212 -2.943655  0.442825
 C   0.035646  0.357103  2.750014
 C   2.091706  3.122428  0.158488
 N   0.631017 -0.471794  1.829106
 Fe  1.703493  0.090952  0.227847
 H   3.204275  4.038144 -1.396612
 H   3.777970 -2.075321 -3.125714
 H   0.494436 -3.860204  2.042940
 H  -0.408169  2.268000  3.550887
 C  -6.464476 -1.054345  0.823012
 O  -7.574226 -1.680474  1.481195
 C  -6.772203  0.366773  0.443067
 C  -6.653878  0.953211 -0.761286
 C  -7.030697  2.406098 -0.944117
 C  -6.151046  0.256696 -2.010329
 C  -4.683314  0.608324 -2.399481
 C  -3.676650  0.157269 -1.387361
 C  -2.658304 -0.735579 -1.542031
 C  -2.317761 -1.434903 -2.837743
 C  -1.789458 -1.027045 -0.409507
 H  -8.330361 -1.618077  0.875356
 H  -6.134165 -1.649493 -0.038837
 H  -5.652467 -1.081811  1.560071
 H  -7.151679  0.965500  1.272570
 H  -6.184521  2.999309 -1.316110
 H  -7.832436  2.509481 -1.688408
 H  -7.371340  2.858444 -0.007723
 H  -6.230503 -0.830735 -1.917814
 H  -6.789690  0.550161 -2.854665
 H  -4.480311  0.181815 -3.385855
 H  -4.605173  1.700300 -2.510888
 H  -3.798209  0.594272 -0.396206
 H  -2.339736 -2.523744 -2.703038
 H  -2.993087 -1.181220 -3.657084
 H  -1.297670 -1.174108 -3.146842
 H  -1.359160 -2.030765 -0.371741
 H  -0.668240 -0.348512 -0.584529
 H  -2.130907 -0.670637  0.563186
 H   5.148470 -1.012179 -0.098755
 H   0.433610  4.581711  2.629746
 H   1.989474  5.330413  0.535685
 H   4.382014  2.804375 -3.403953
 H   4.613394  0.205844 -4.149000
 H   3.066585 -4.381086 -2.125137
 H   1.691725 -5.151627  0.084676
 H  -1.203138 -0.001043  4.585125
 H  -0.798884 -2.611876  3.964396

TS C8-HAT formation
(q, Cpd I + Nerol)
 C   4.715341 -1.320242  1.186898
 S   3.531887 -0.050071  1.750379
 H   4.224042 -2.296049  1.120974
 H   5.514315 -1.383332  1.931894
 N   2.847066 -0.609949 -1.292353
 N   1.269916 -1.749041  0.807566
 N   2.254028  1.999593 -0.346844
 C   3.680688  1.480509 -2.273929
 C   2.364557 -3.005525 -0.989227
 C  -0.008826 -1.223449  2.834881
 C   1.099627  3.274547  1.407546
 C   3.566071  0.095677 -2.225369
 C   1.593911 -2.915156  0.166982
 C   0.107521  0.160409  2.787973
 C   1.868587  3.173415  0.257442
 O   0.315573  0.223164 -0.785131
 C   4.154216 -0.805367 -3.185851
 C   1.042194 -4.045033  0.879062
 C  -0.497004  1.059522  3.740378
 C   2.424347  4.302931 -0.451899
 C   3.764467 -2.062922 -2.836141
 C   0.386892 -3.542826  1.962411
 C  -0.193147  2.324840  3.336850
 C   3.162565  3.800033 -1.477772
 C   2.945676 -1.934762 -1.655604
 C   0.533394 -2.107130  1.910058
 C   0.599696  2.200289  2.137901
 C   3.055033  2.361375 -1.403515
 N   0.779840  0.878321  1.825549
 Fe  1.743313  0.147497  0.198867
 H   4.287059  1.904813 -3.067816
 H   2.538999 -3.996888 -1.395651
 H  -0.575887 -1.647336  3.657709
 H   0.879530  4.272050  1.774471
 C  -6.521187 -1.127523  0.745540
 O  -7.680067 -1.727568  1.341455
 C  -6.730413  0.335265  0.472422
 C  -6.569055  0.999692 -0.685595
 C  -6.848359  2.483990 -0.758787
 C  -6.109639  0.367095 -1.984428
 C  -4.625950  0.660643 -2.355244
 C  -3.641288  0.072196 -1.389750
 C  -2.644951 -0.826612 -1.633240
 C  -2.320024 -1.392755 -2.996647
 C  -1.772741 -1.254326 -0.541885
 H  -8.420944 -1.566596  0.734885
 H  -6.216250 -1.675952 -0.155631
 H  -5.724801 -1.264262  1.487392
 H  -7.072778  0.895385  1.344031
 H  -5.962465  3.045964 -1.083603
 H  -7.636550  2.695876 -1.494585
 H  -7.164650  2.886266  0.208522
 H  -6.252695 -0.717783 -1.969401
 H  -6.733945  0.757050 -2.800257
 H  -4.452308  0.300852 -3.373566
 H  -4.483757  1.751544 -2.386150
 H  -3.755646  0.408806 -0.359044
 H  -2.358867 -2.489415 -2.975381
 H  -2.992474 -1.045097 -3.783443
 H  -1.296477 -1.118682 -3.283101
 H  -1.368251 -2.265990 -0.629190
 H  -0.683660 -0.544062 -0.618020
 H  -2.130086 -1.030725  0.465606
 H   5.139183 -1.068299  0.212075
 H   2.265576  5.337858 -0.176864
 H   3.736024  4.334481 -2.224303
 H   4.772899 -0.496442 -4.018631
 H   4.000122 -3.002063 -3.320259
 H   1.157675 -5.078489  0.578015
 H  -0.150549 -4.077900  2.734906
 H  -0.472696  3.265463  3.793906
 H  -1.079212  0.745130  4.597126
```

**Table S11** The number of models generated using Chai-1 for each design/substrate pair and the ligand SMILES notation used for modeling.

| Model | Number of models generated | Substrate SMILES notation |
| --- | --- | --- |
| Design 18/α-damascone (5) | 50 | C/C=C/C(=O)C1C(=CCCC1(C)C)C |
| Design 18/β-damascone (6) | 10 | O=C(/C1=C(/CCCC1(C)C)C)/C=C/C |
| Design 18/α-ionone (8) | 10 | CC1=CCCC(C1/C=C/C(=O)C)(C)C |
| Design 18/δ-damascone (7) | 10 | C/C=C/C(=O)C1C(C=CCC1(C)C)C |
| Design 11/δ-damascone (7) | 10 | C/C=C/C(=O)C1C(C=CCC1(C)C)C |

**II Supplementary Figs**

**Fig. S1** Calibration curves for determining TON using GC-MS for **A)** citral A, **B)** citral B, **C)** 2,3-epoxy nerol), **D)** carveol.

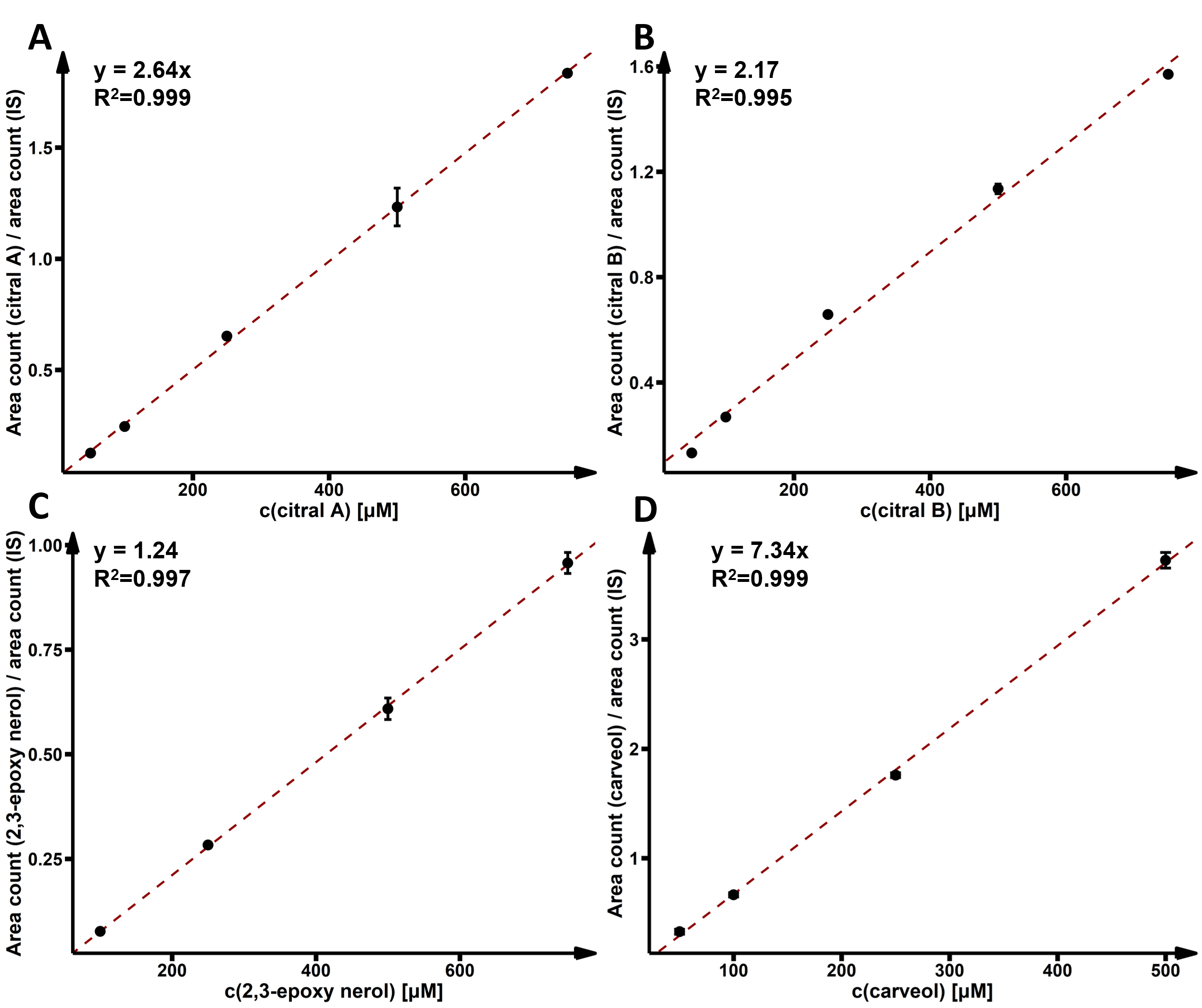

**Fig. S2** Fragmentation of 2-hydroxy-δ-damascone.

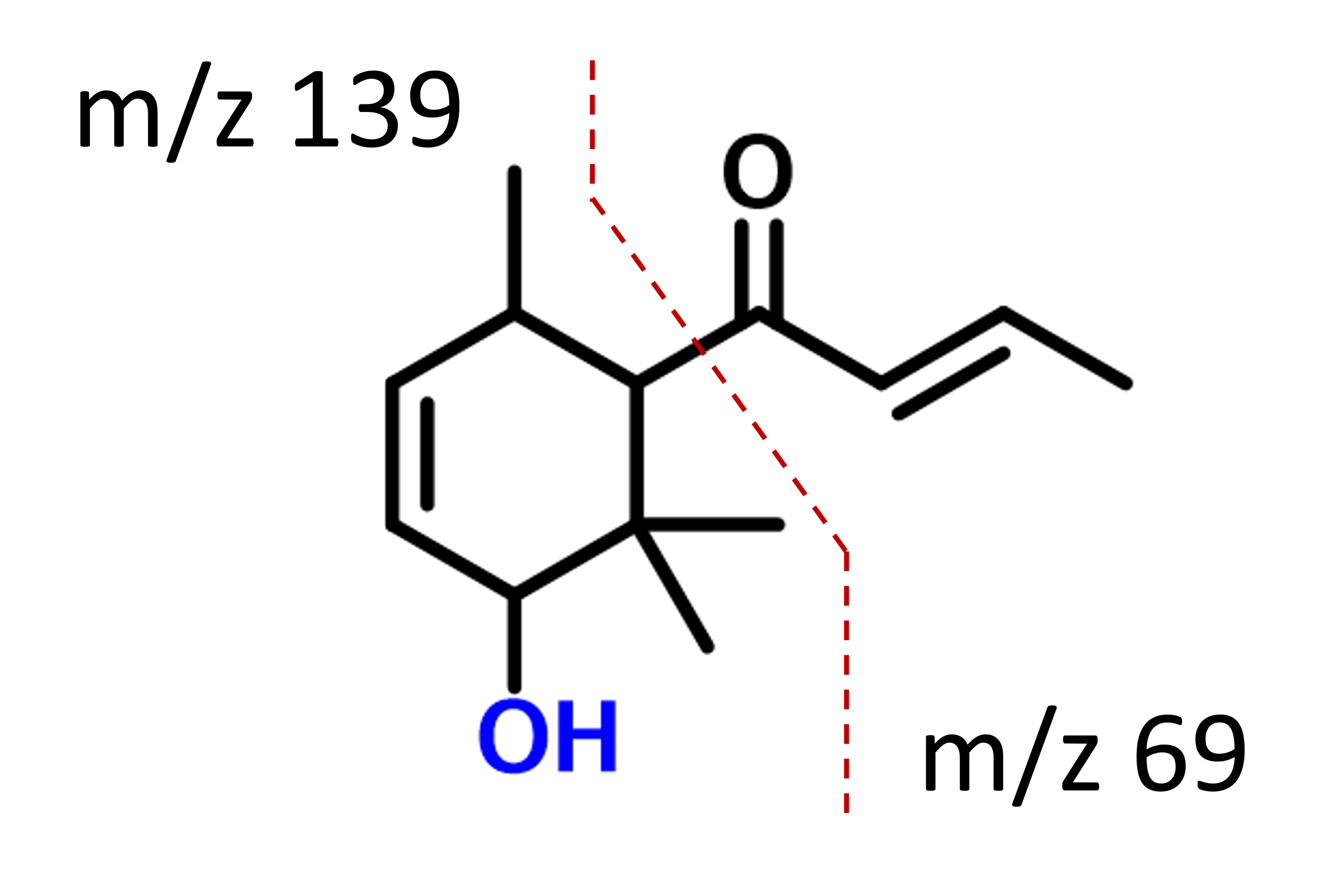

**Fig. S3** GC-MS chromatogram after the reaction of (*S*)-(-)-limonene with wt *Mth*UPO and extraction with EtOAc (IS: 4-octanone).

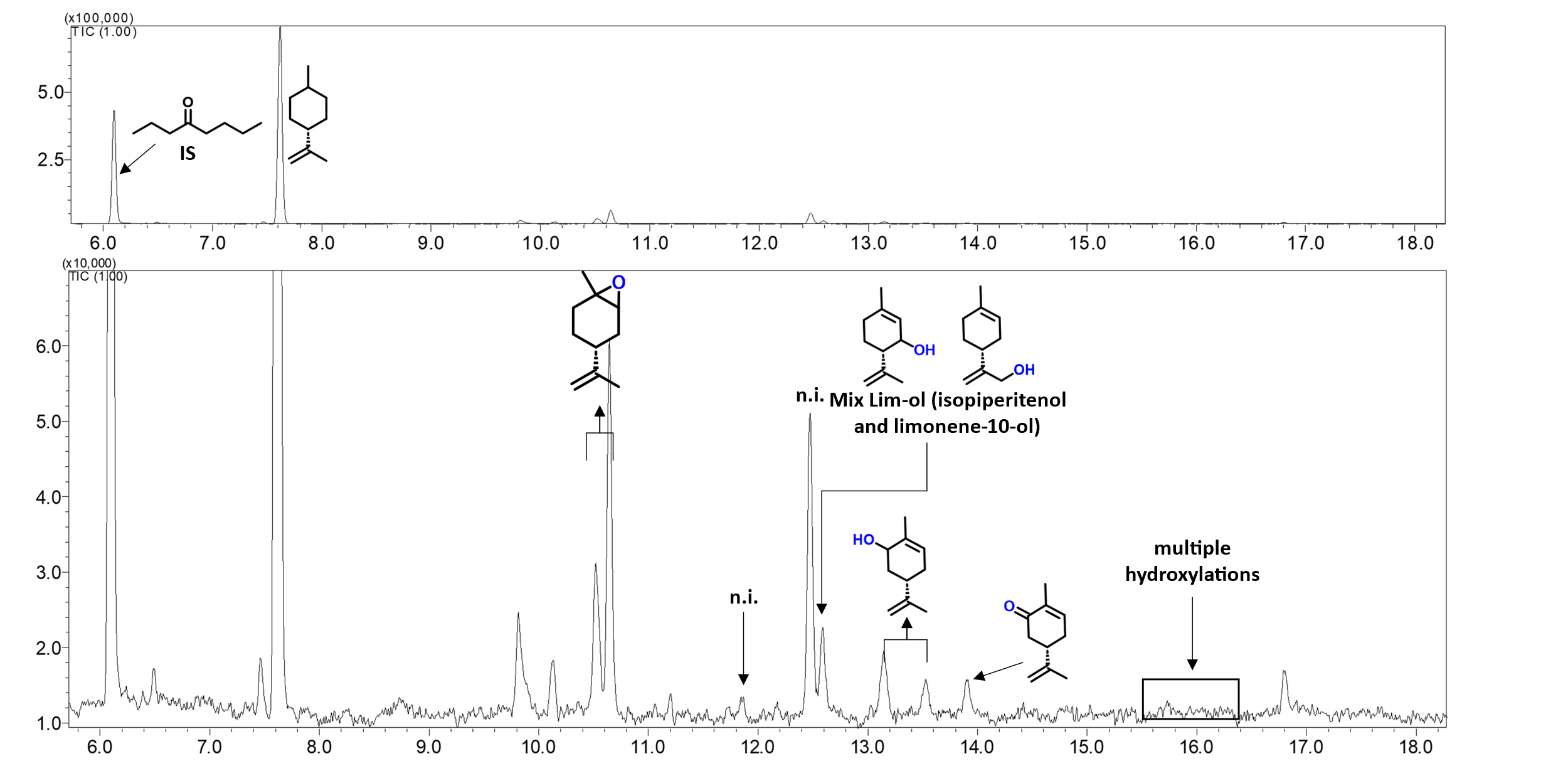

**Fig. S4** Chiral GC-MS chromatograms after enzymatic conversion of β-ionone catalyzed by **A)** wt *Mth*UPO, **B)** FuncLib Var 11, **C)** FuncLib Var 34.

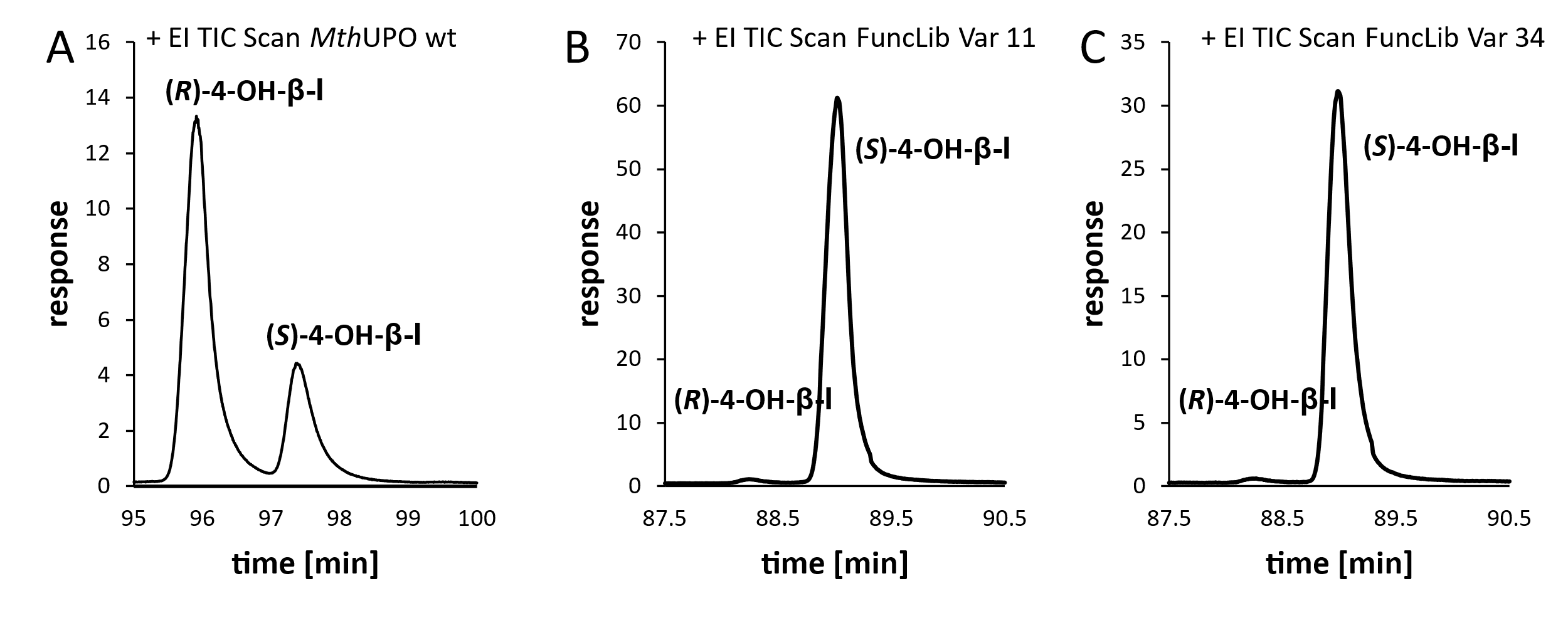

**Fig. S5** GC-MS chromatogram after the reaction of **A** δ-damascone and **B** α-damascone with wt *Mth*UPO (grey) and different FuncLib variants after extraction with EtOAc (β-ionone, elution time 12.25 min).

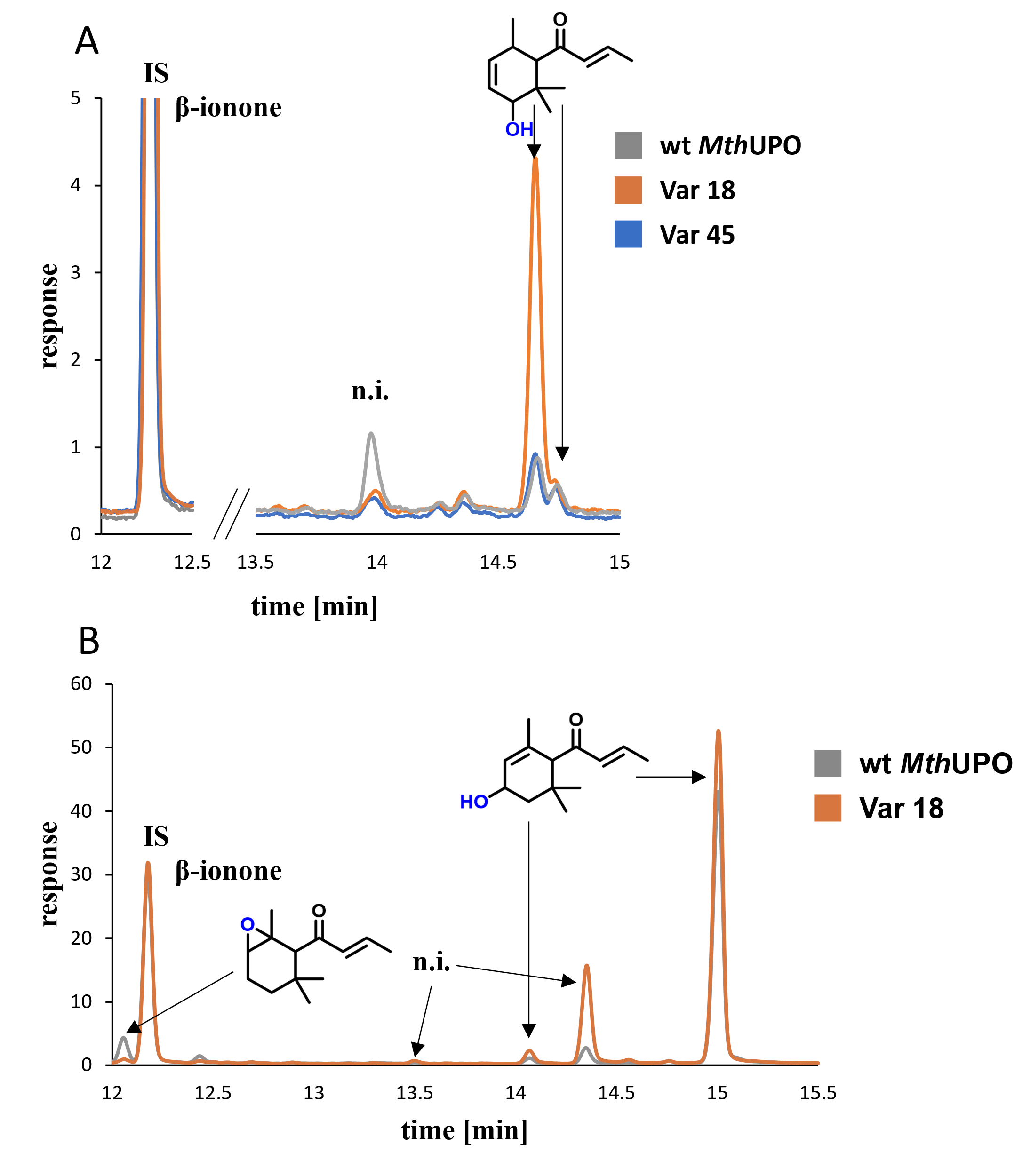

**Fig. S6** DFT model calculations exploring the intrinsic oxidation pattern of geraniol.

**A)** Schematic structure and atom numbering of geraniol (**1**).

**B)** DFT computed relative energies using a computational truncated model based on *Mth*UPO active site (iron-oxo species coordinated to the porphyrin pyrrole core and methane thiolate as the axial ligand) for all rate-limiting transition states (TS) leading to different oxidation products of **1**, including TS C2-O formation involved in geraniol epoxidation and Hydrogen Atom Transfers (TS HAT) involved in geraniol hydroxylation. Energy values were obtained at the (U)B3LYP/Def2TZVP/PCM(dichloromethane)// (U)B3LYP/6-31G(d)+SDD(Fe)/PCM(dichloromethane) level (see details on the Computational Methods section). The reported relative energies were computed in doublet (d) and quartet (q) electronic states in terms of electronic energy (ΔE), enthalpy (ΔH), and quasi-harmonic corrected Gibbs energy (ΔG). All energies are referred considering the quartet separated reactants **[Cpd I + geraniol (q)]** as zero.

**C)** Optimized geometries for the stationary points reported in B). Energies are referred considering as zero the lowest in energy TS **[TS C1-HAT (q)]**.

All energies are given in kcal·mol^-1^. Distances, and angles are given in Angstrom (Å) or degrees (º), respectively.

**A)**

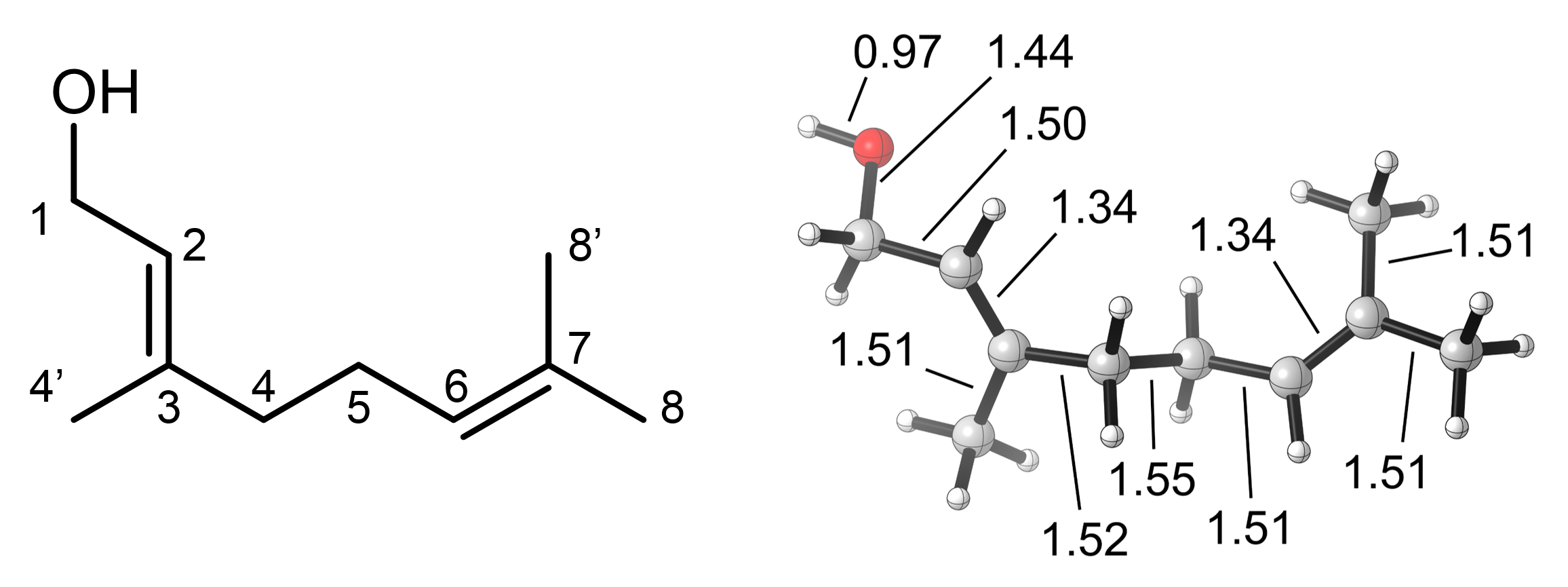

**B)**

| **Cpd I + Geraniol** | **Electronic state** | **ΔE** | **ΔH** | **ΔG** |
| --- | --- | --- | --- | --- |
| **TS C2-O formation** | doublet (d) | 3.1 | 3.3 | 20.5 |
|  | quartet (q) | 3.2 | 3.5 | 21.2 |
| **TS C1-HAT** | doublet (d) | 4.8 | 2.1 | 19.6 |
|  | quartet (q) | 2.9 | -0.1 | 18.3 |
| **TS C4-HAT** | doublet (d) | 7.1 | 3.5 | 21.1 |
|  | quartet (q) | 6.9 | 3.4 | 21.1 |
| **TS C5-HAT** | doublet (d) | 5.8 | 2.9 | 20.9 |
|  | quartet (q) | 7.5 | 4.4 | 21.9 |
| **TS C8-HAT** | doublet (d) | 9.3 | 6.6 | 23.5 |
|  | quartet (q) | 9.2 | 6.0 | 23.2 |

**C)**

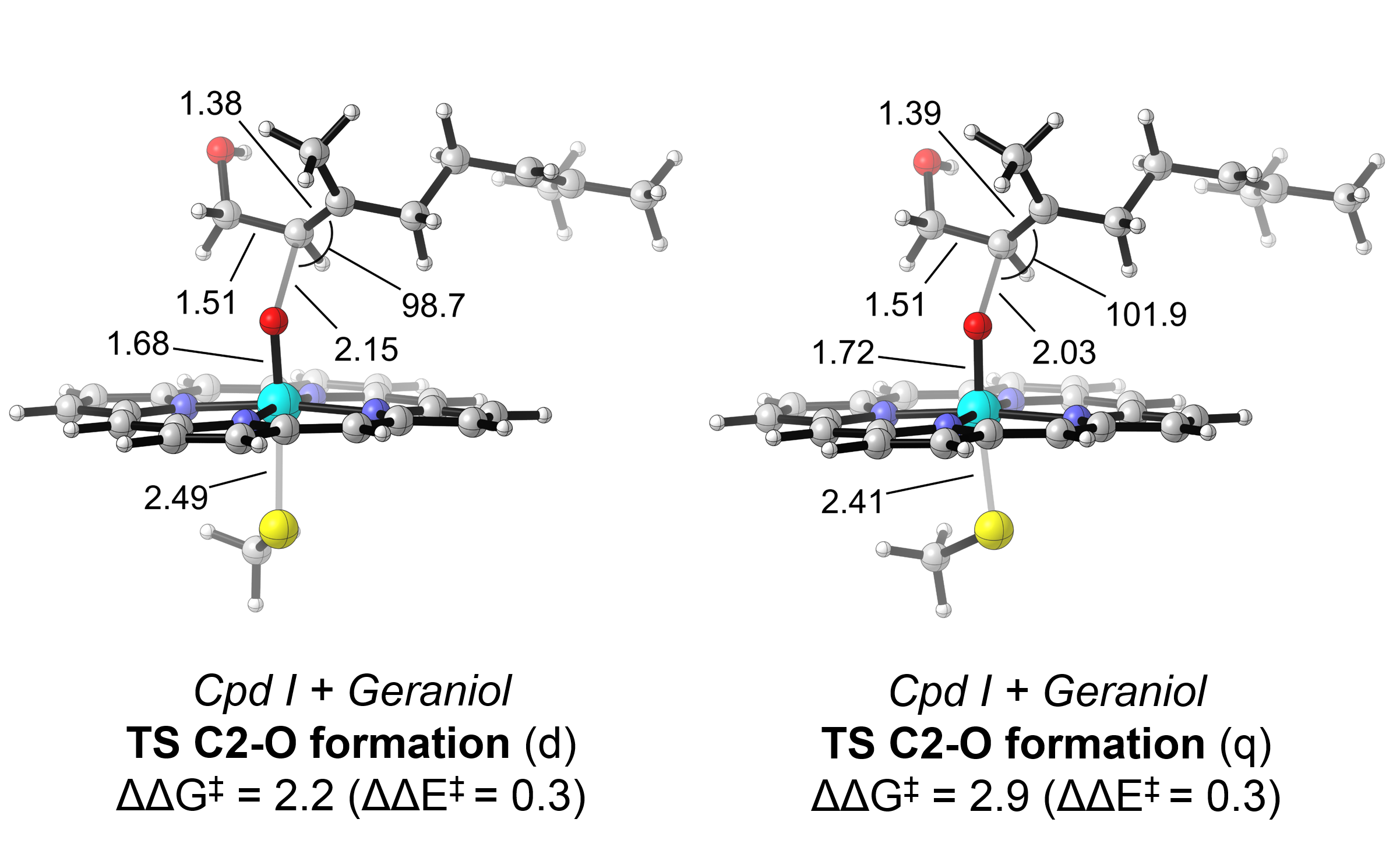

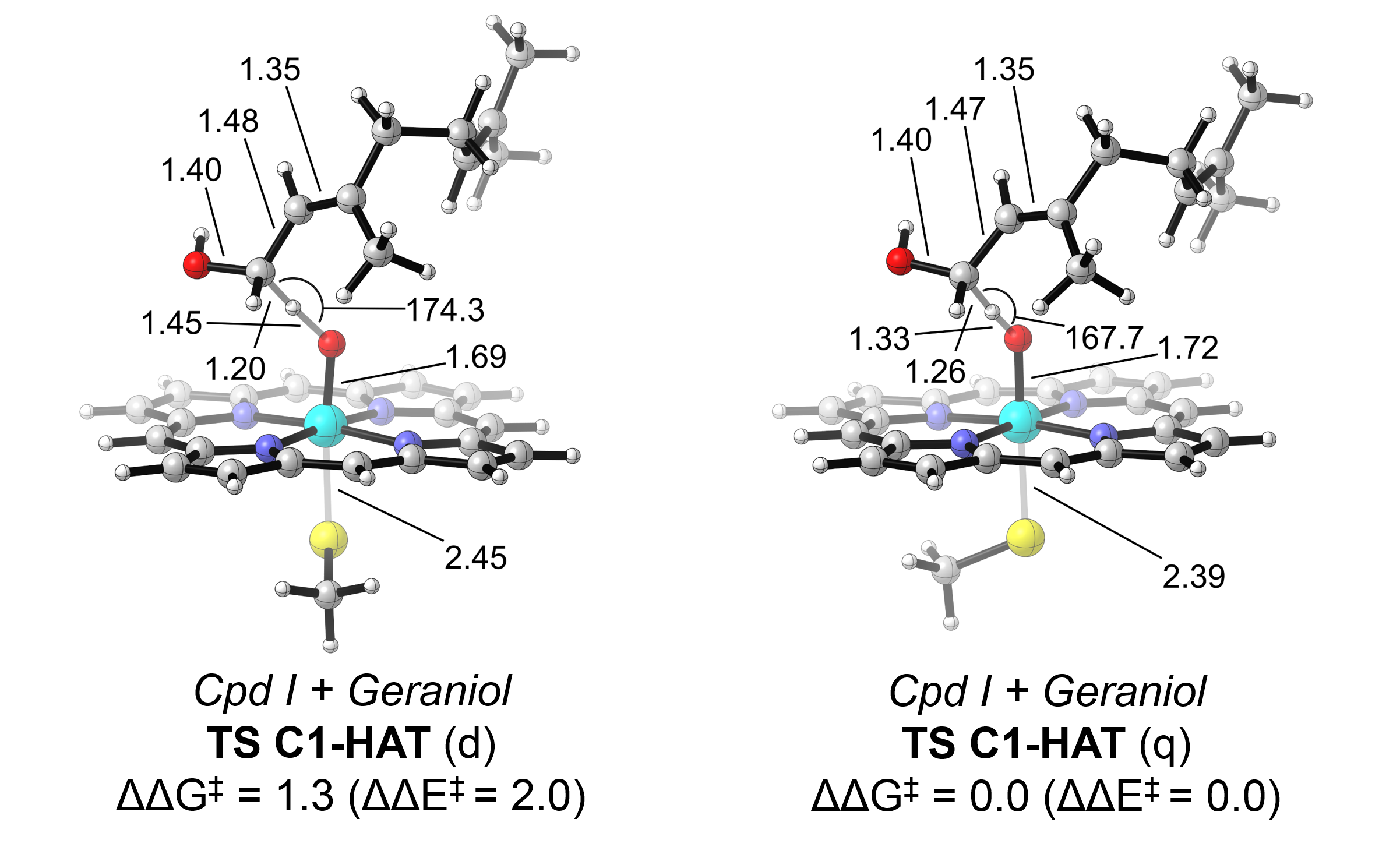

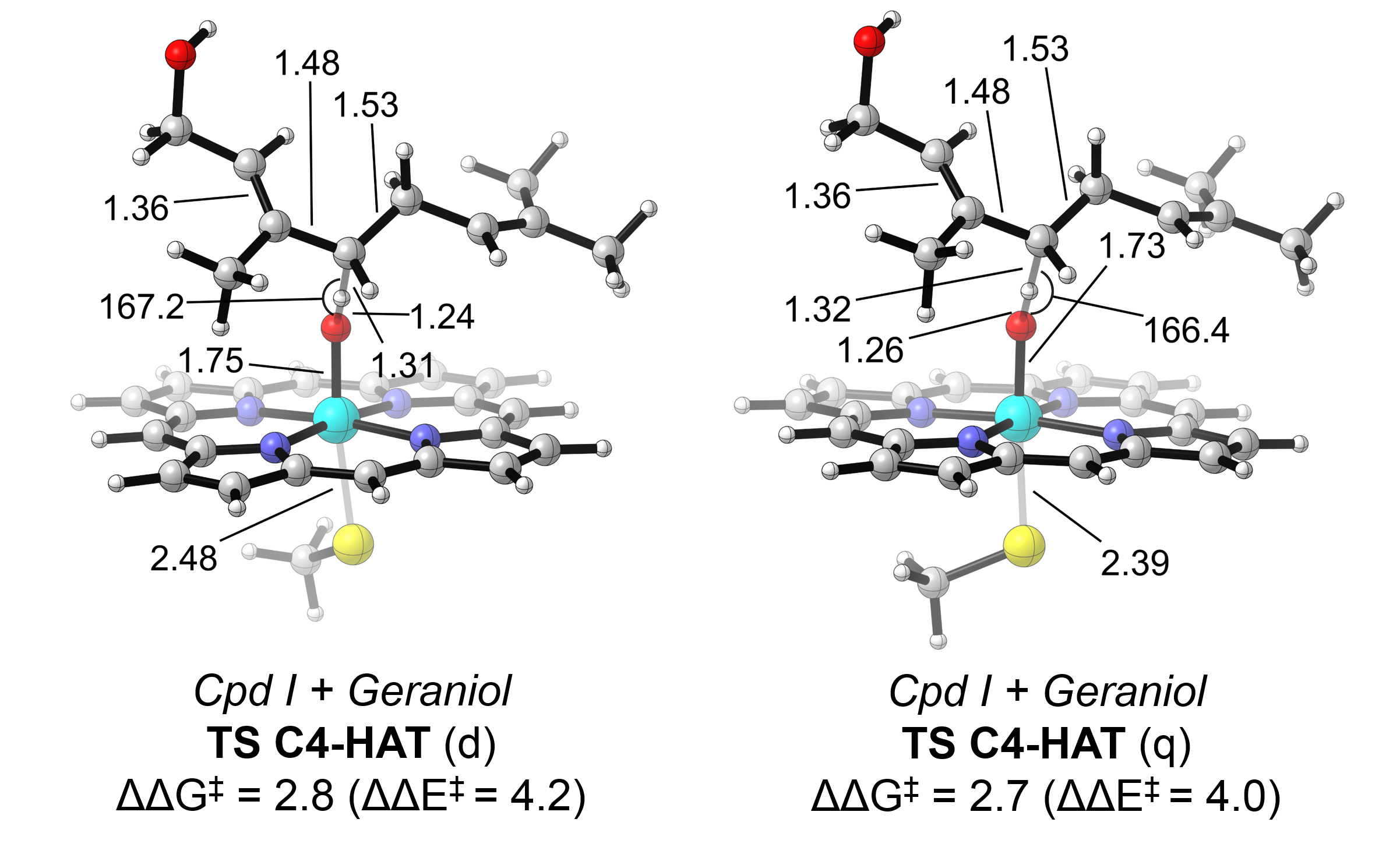

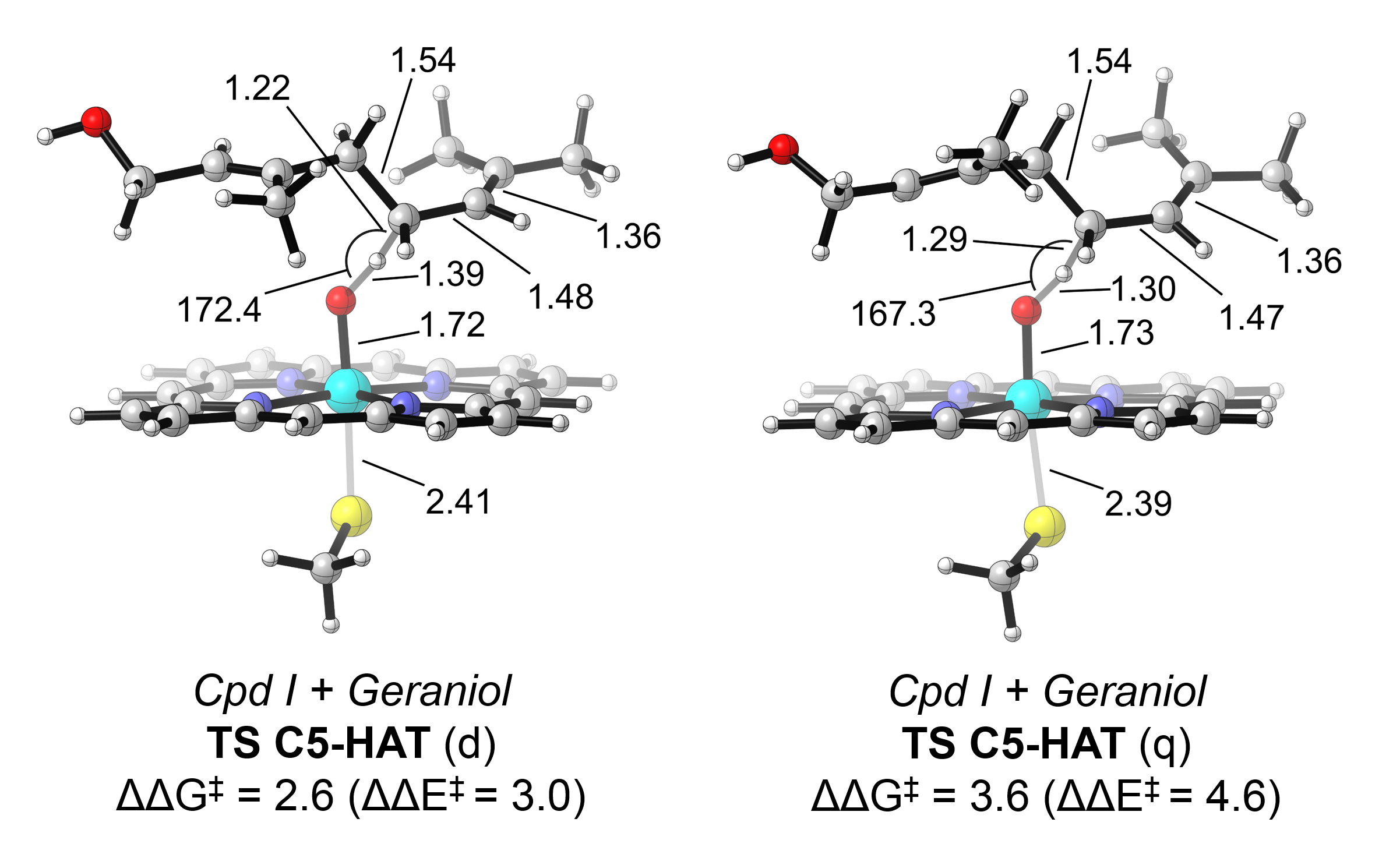

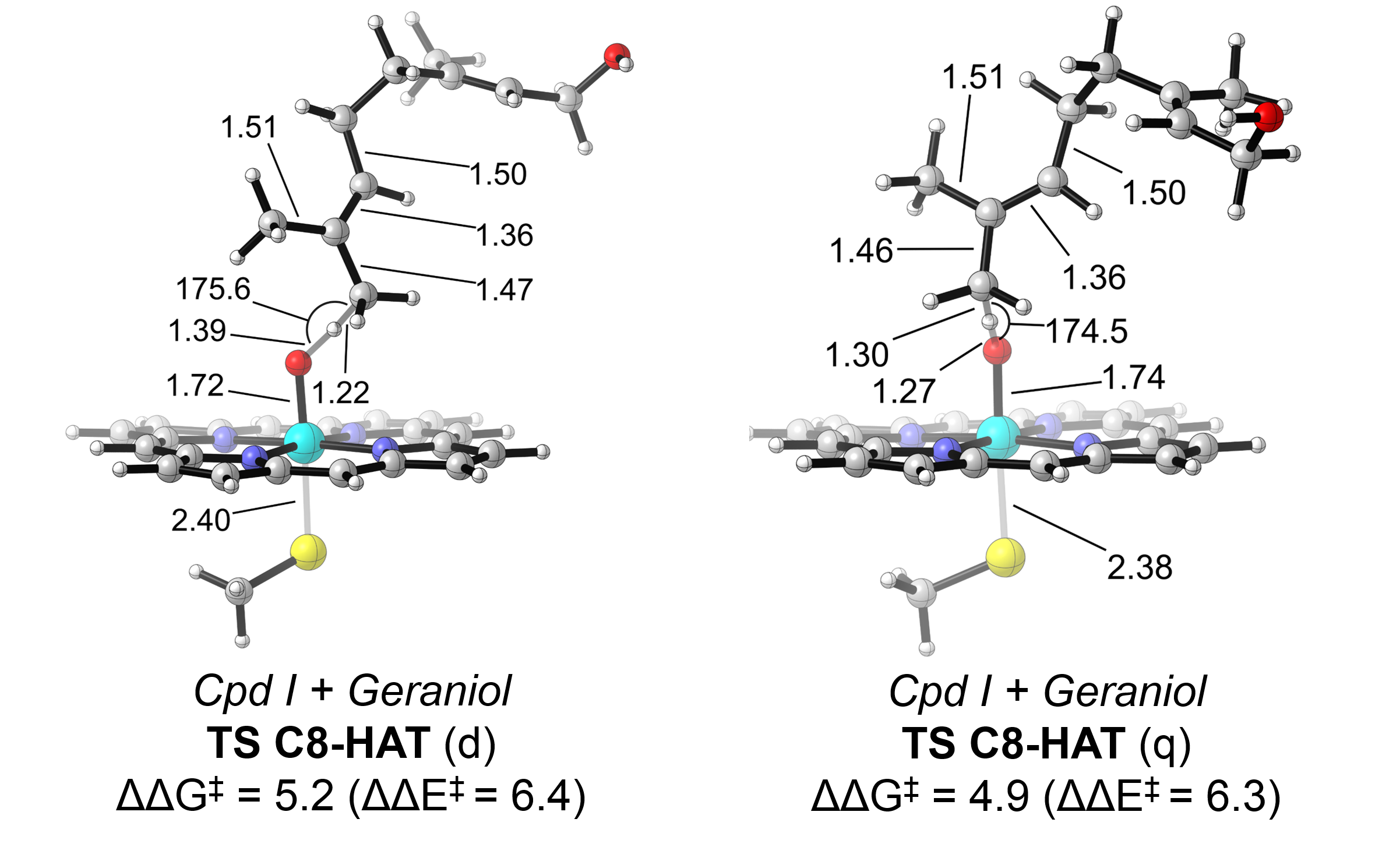

Truncated DFT models for geraniol oxidation by Cpd I indicates that allylic hydroxylation at C1 is the most energy accessible oxidation (**TS C1-HAT**), followed by epoxidation at the double C2-C3 bond (**TS C2-O formation**). The hydroxylations at positions 4, 5, and 8 are more energetically demanding and are not expected to occur in *Mth*UPO active site.

**Fig. S7** DFT model calculations exploring the intrinsic oxidation pattern of nerol.

**A)** Schematic structure and atom numbering of nerol (**2**).

**B)** DFT computed relative energies using a computational truncated model based on *Mth*UPO active site (iron-oxo species coordinated to the porphyrin pyrrole core and methane thiolate as the axial ligand) for all rate-limiting transition states (TS) leading to different oxidation products of **2**, including TS C2-O formation involved in geraniol epoxidation and Hydrogen Atom Transfers (TS HAT) involved in geraniol hydroxylation. Energy values were obtained at the (U)B3LYP/Def2TZVP/PCM(dichloromethane)// (U)B3LYP/6-31G(d)+SDD(Fe)/PCM(dichloromethane) level (see details on the Computational Methods section). The reported relative energies were computed in doublet (d) and quartet (q) electronic states in terms of electronic energy (ΔE), enthalpy (ΔH), and quasi-harmonic corrected Gibbs energy (ΔG). All energies are referred considering the quartet separated reactants **[Cpd I + nerol (q)]** as zero.

**C)** Optimized geometries for the stationary points reported in B). Energies are referred considering as zero the lowest in energy TS **[TS C1-HAT (q)]**.

All energy is given in kcal·mol^-1^. Distances, and angles are given in Angstrom (Å) or degrees (º), respectively.

**A)**

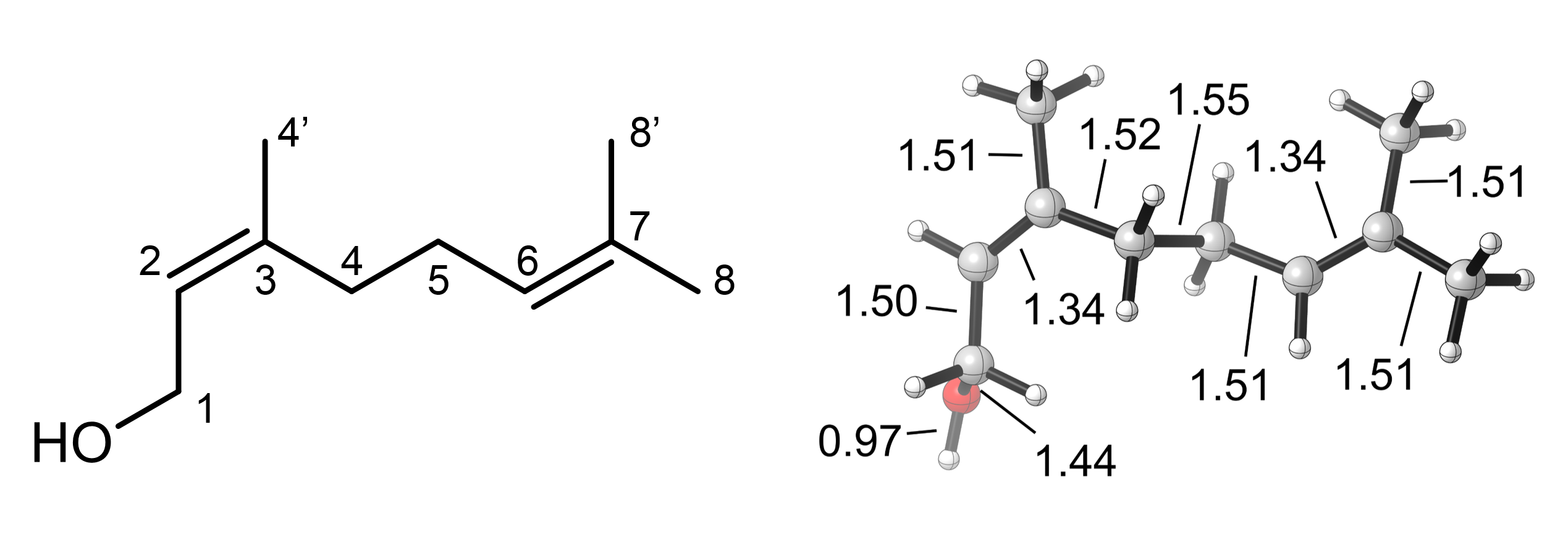

**B)**

| **Cpd I + Nerol** | **Electronic state** | **ΔE** | **ΔH** | **ΔG** |
| --- | --- | --- | --- | --- |
| **TS C2-O formation** | doublet (d) | 1.5 | 1.8 | 19.5 |
| (conformer 1) | quartet (q) | 2.6 | 2.9 | 21.3 |
| **TS C2-O formation** | doublet (d) | -2.6 | -2.2 | 17.0 |
| (conformer 2) | quartet (q) | -2.3 | -2.0 | 17.4 |
| **TS C1-HAT** | doublet (d) | 0.3 | -2.5 | 16.4 |
|  | quartet (q) | -0.1 | -3.1 | 16.0 |
| **TS C4-HAT** | doublet (d) | 4.8 | 1.6 | 20.1 |
|  | quartet (q) | 5.6 | 2.3 | 21.1 |
| **TS C5-HAT** | doublet (d) | 4.6 | 1.7 | 19.9 |
|  | quartet (q) | 4.7 | 1.5 | 19.8 |
| **TS C8-HAT** | doublet (d) | 11.5 | 8.6 | 26.2 |
|  | quartet (q) | 8.0 | 4.8 | 22.4 |

**C)**

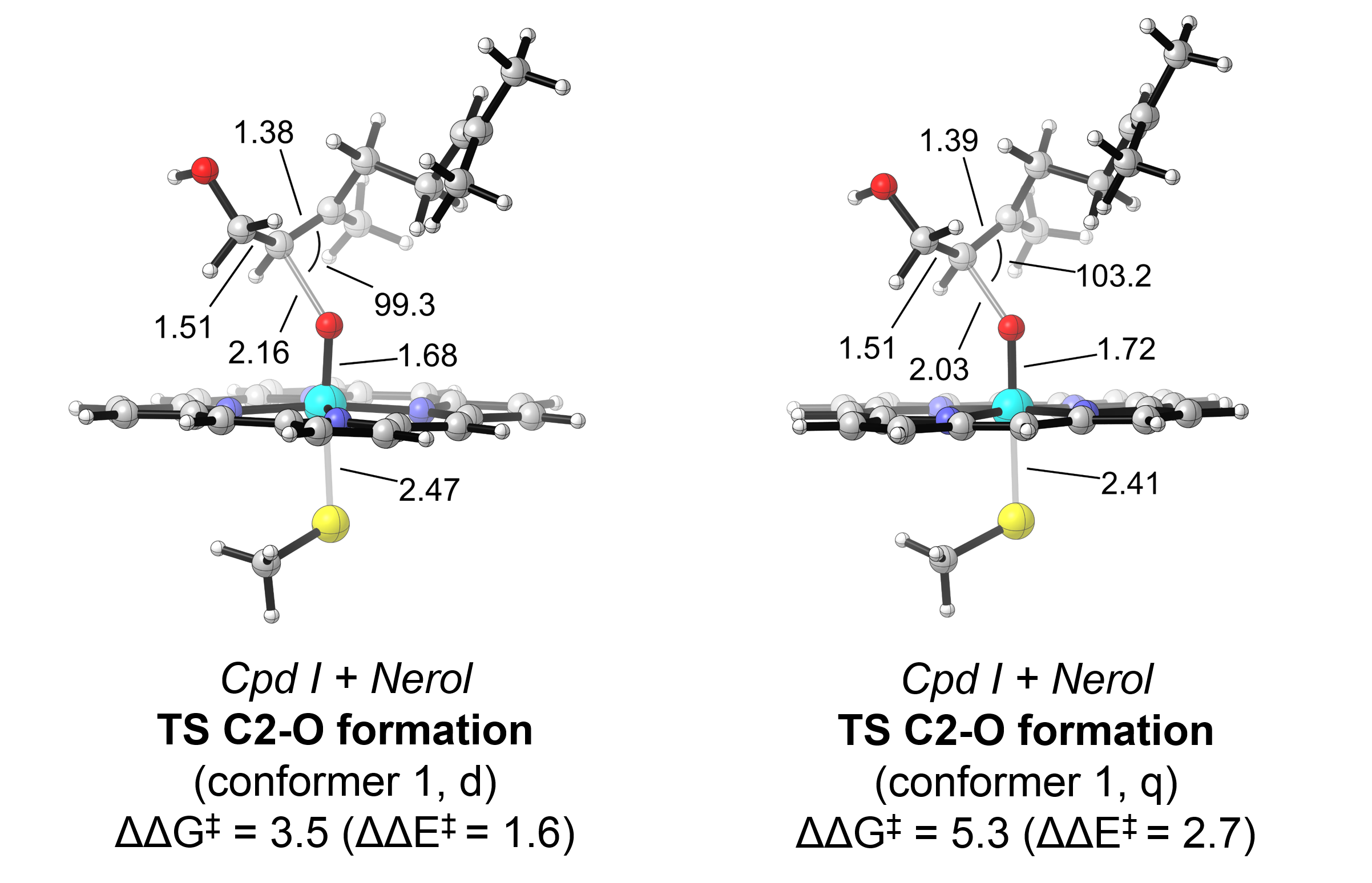

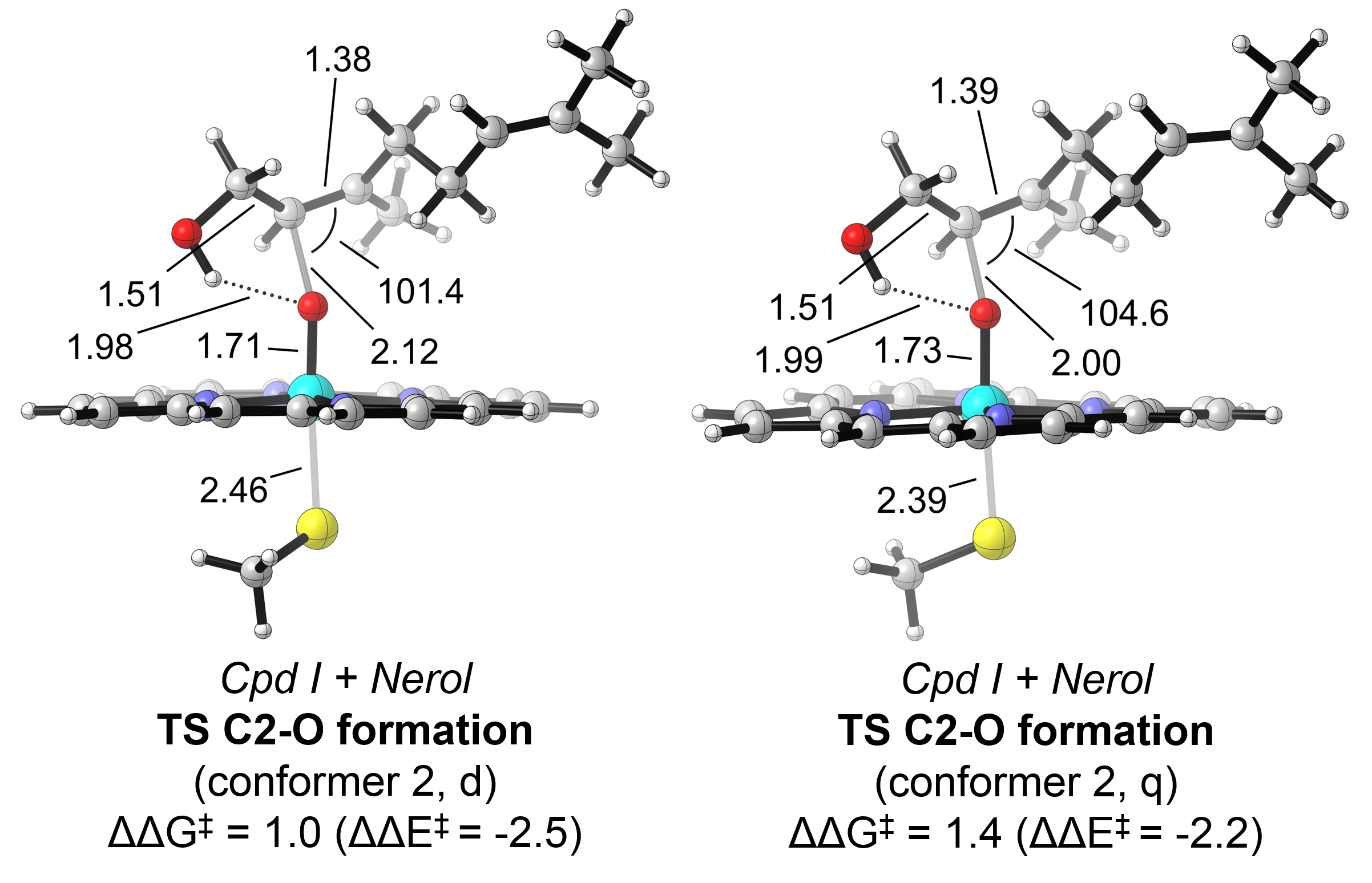

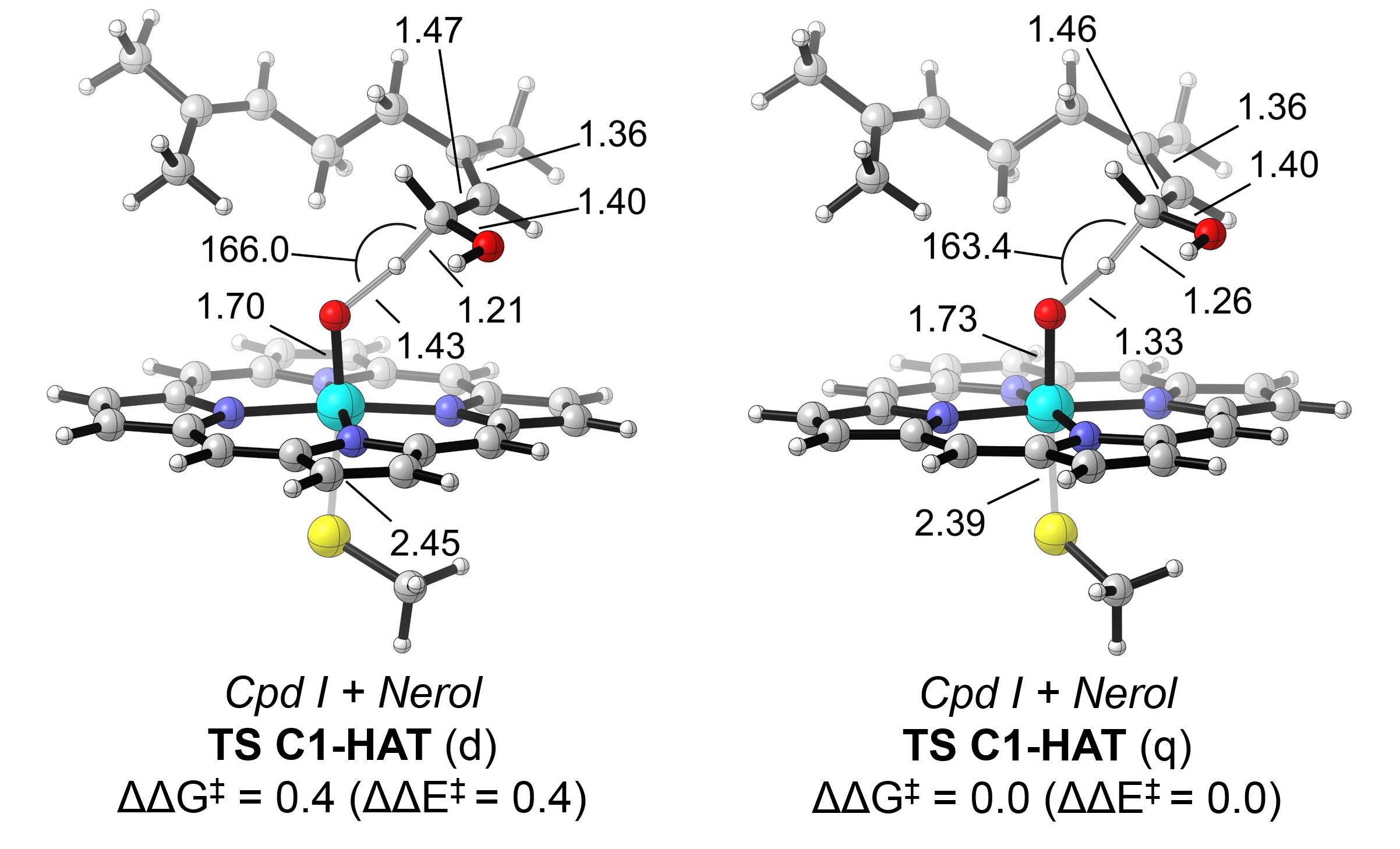

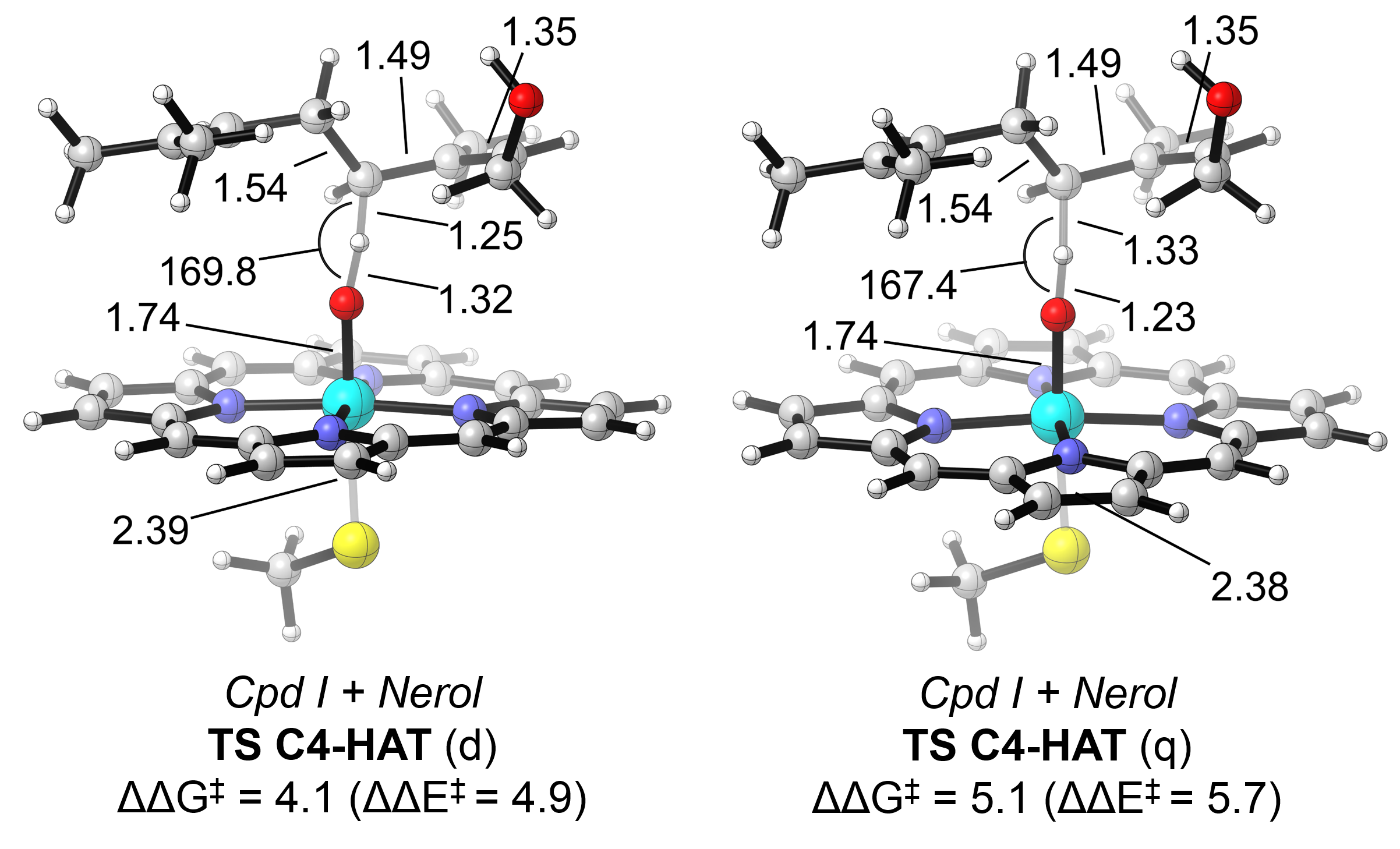

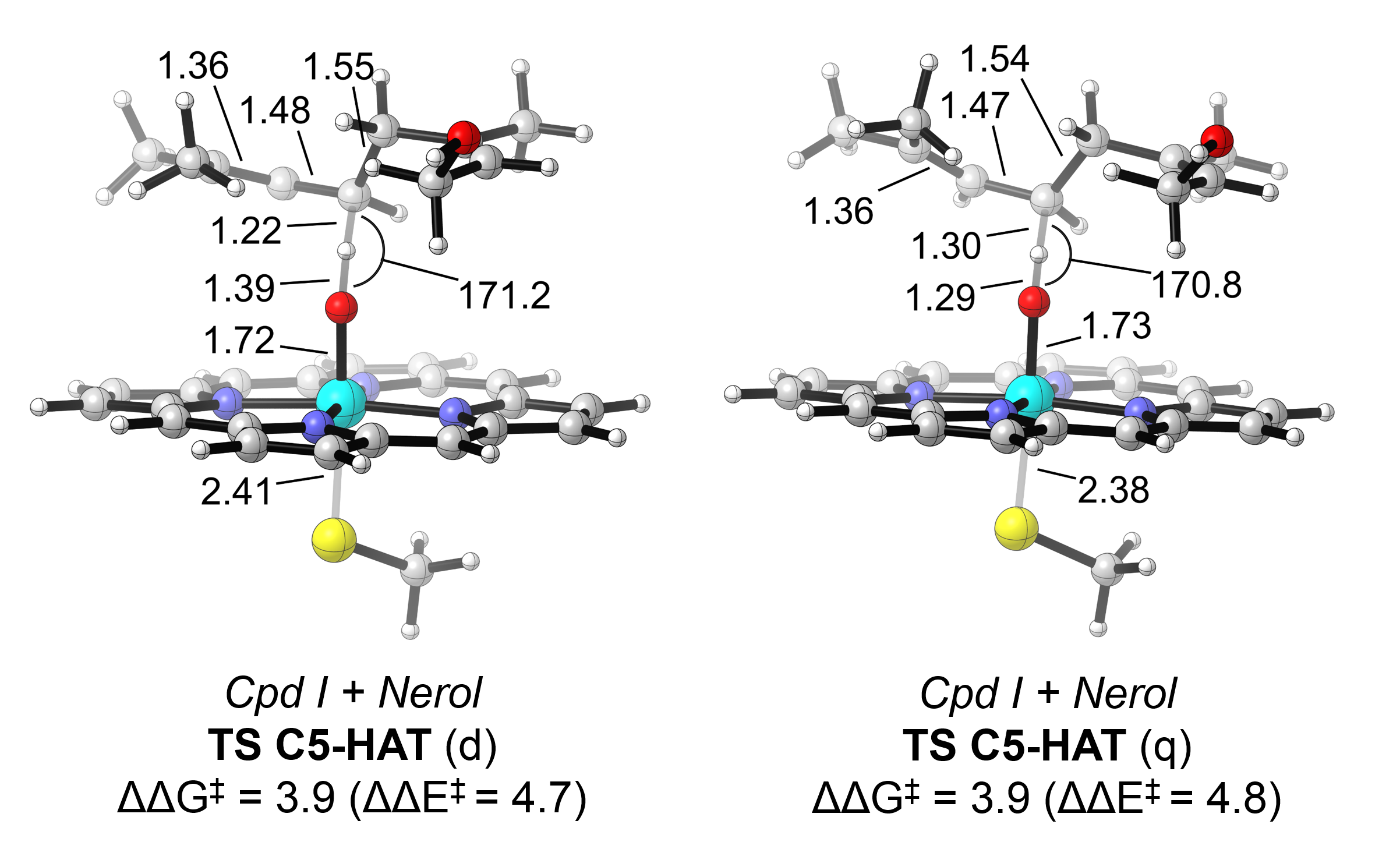

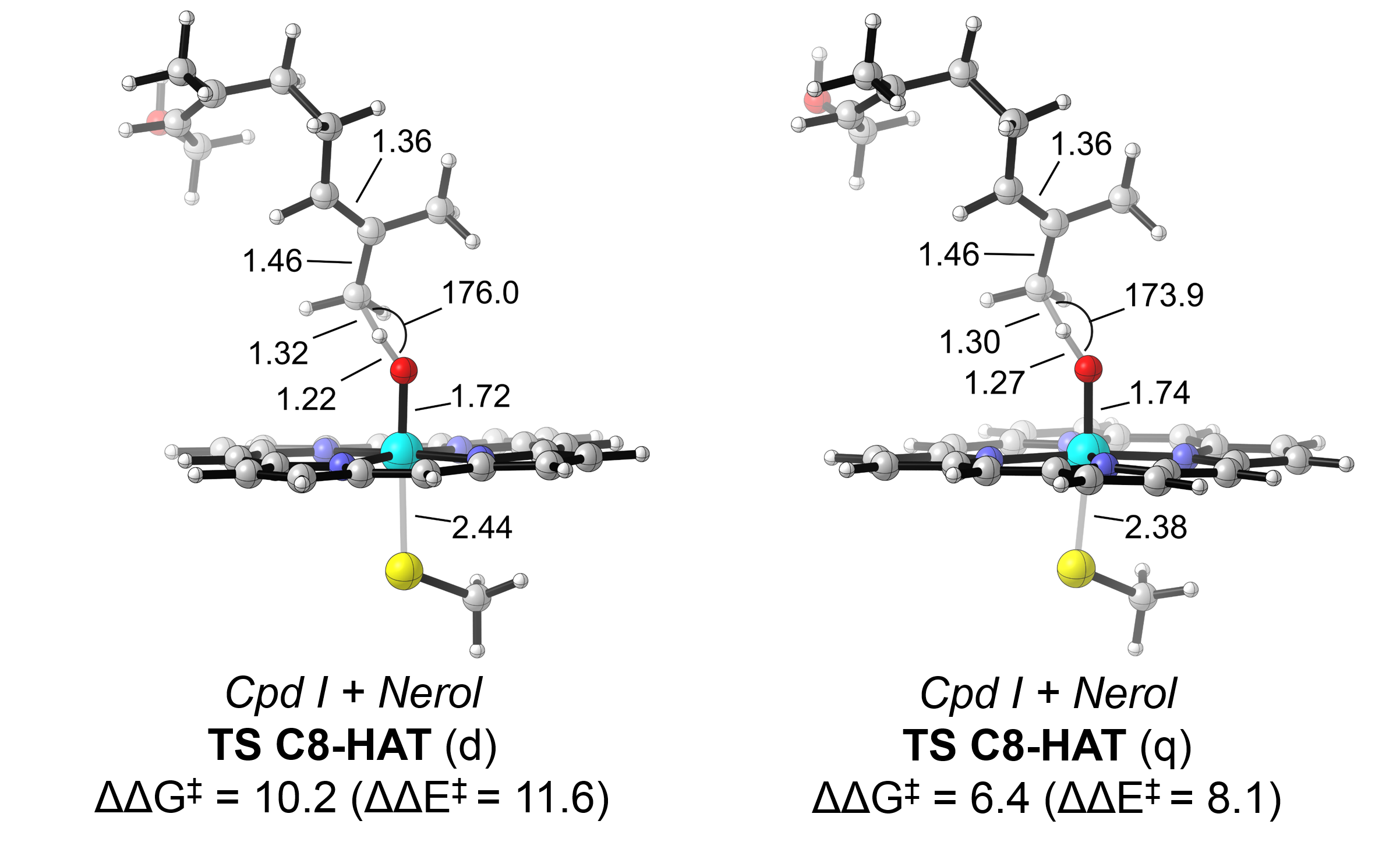

Truncated DFT models for nerol oxidation by Cpd I indicates that allylic hydroxylation at C1 is the most energy accessible oxidation (**TS C1-HAT**), followed by epoxidation at the double C2-C3 bond (**TS C2-O formation**).

MD simulations on Design 28 with nerol bound in the active site (**Fig. S11**) indicate that the substrate can perform a hydrogen bond with the iron-oxo group of the Cpd I species. This alternative conformation is also tested using truncated DFT models (**TS C2-O formation**, *conformer 2*), but even if lower in energy than *conformer 1*, it is found to be slightly higher in energy than HAT at C1.

The hydroxylations at positions 4, 5, and 8 are more energetically demanding and are not predicted to occur in *Mth*UPO active site.

**Fig. S8** Analysis of geraniol (**1**) binding mode in Design 4 (F63I/F154L/G157A/ A161F) through substrate-bound MD simulations.

The following analysis allows to characterize which is the preferred binding pose of the substrate in the active site and to elucidate which position of the substrate is more accessible for effective near-attack conformations (NACs) that resemble the geometric requirements of the DFT optimized TS during the MD simulations. Only substrate positions that exhibit a low energetic cost for oxidation as determined by truncated DFT calculations in **Fig. S6** are analyzed in the MD simulations (C2-C3 epoxidation or C1 hydroxylation).

The following geometric parameters are used to characterize the near-attack conformations in the heat map plots: the O–C2 distance and O–C2=C3 angle are used to characterize NACs for effective C2-C3 epoxidation while O–H1 distance and O–H1–C1 angle are used to characterize NACs for effective C1 hydroxylation.

The resulting heat maps were obtained from 3 independent MD simulations of 500 ns each (a total of 1500 ns) without any restraint on the substrate. Ideal geometric parameters obtained from DFT Transition State (TS) optimizations (**Fig. S6**) are shown as black dots. A schematic representation of the different geometric parameters used for the heat maps are also provided in each subfigure.

**A)** NACs for C1-HAT leading to geraniol hydroxylation.

**B)** NACs for C2-O formation leading to geraniol epoxidation.

**C)** NACs for pro-*R* and pro-*S* C1-HAT leading to enantioselective geraniol hydroxylation.

**D)** Polar interaction between the terminal alcohol group of geraniol and the sidechain of E158.

**E)** Most representative structure of the binding mode of geraniol in Design 4 active site as determined by clustering analysis.

Distances and angles are given in angstroms (Å) and degrees (º), respectively.

**A)**

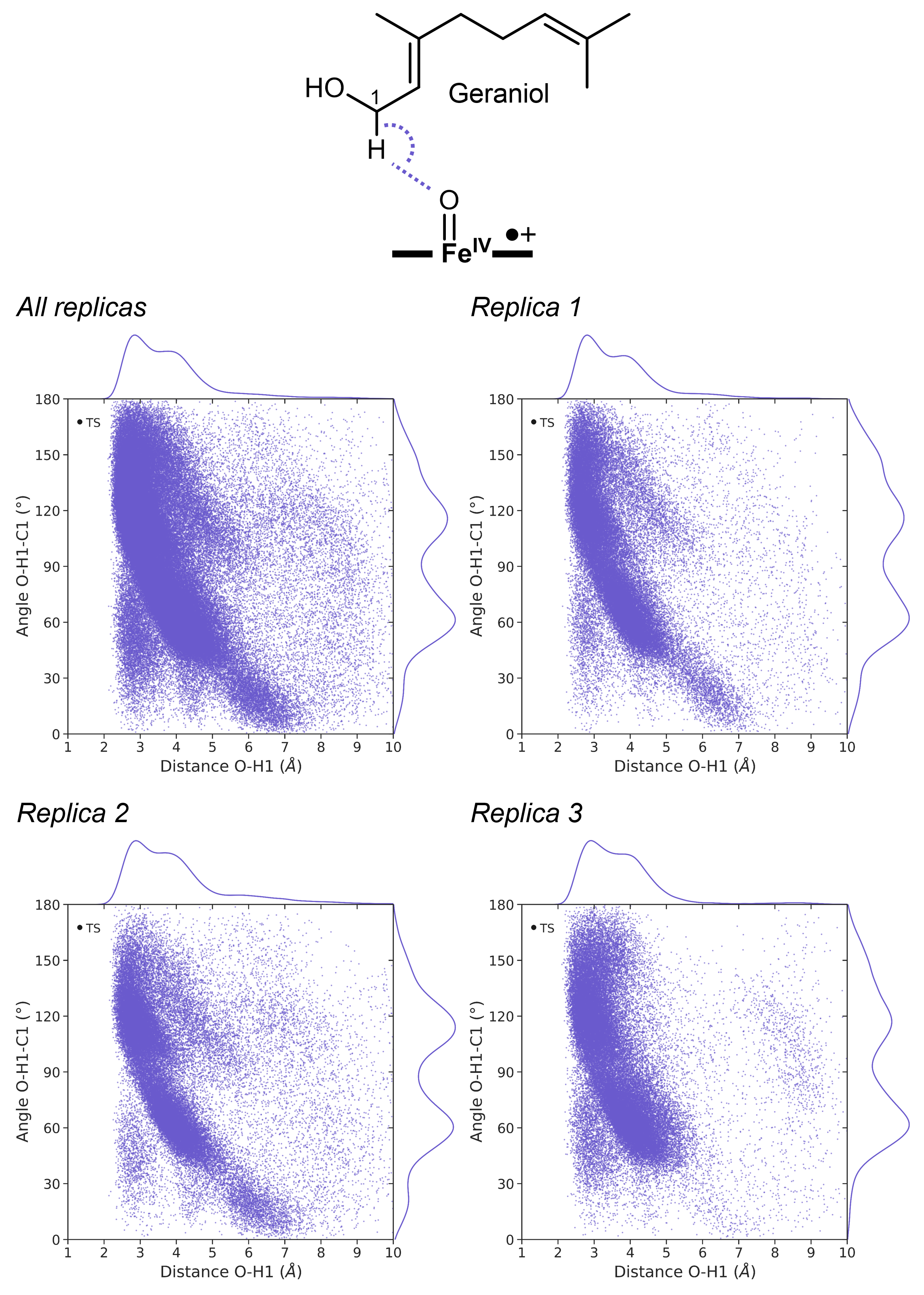

**B)**

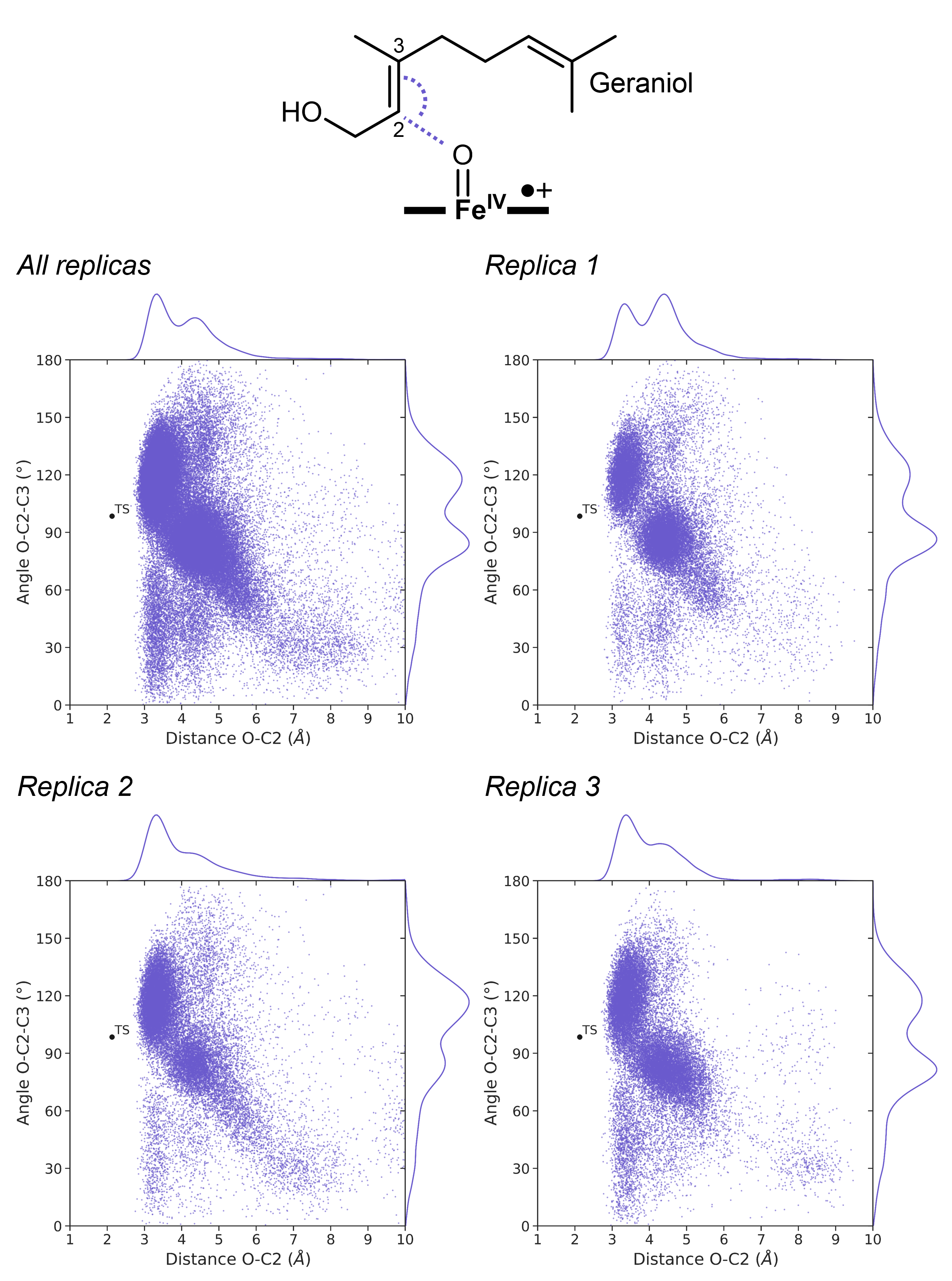

**C)**

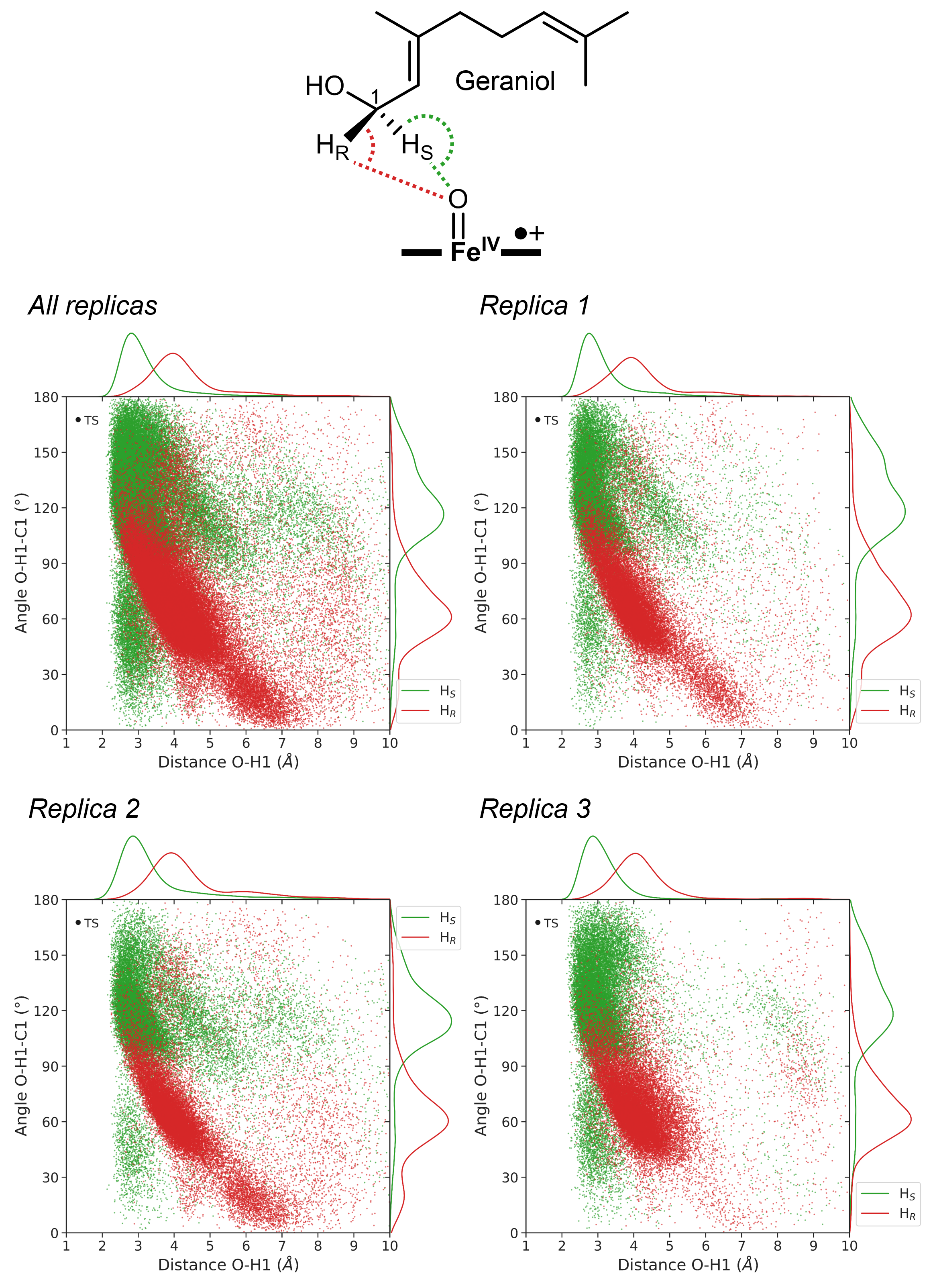

**D)**

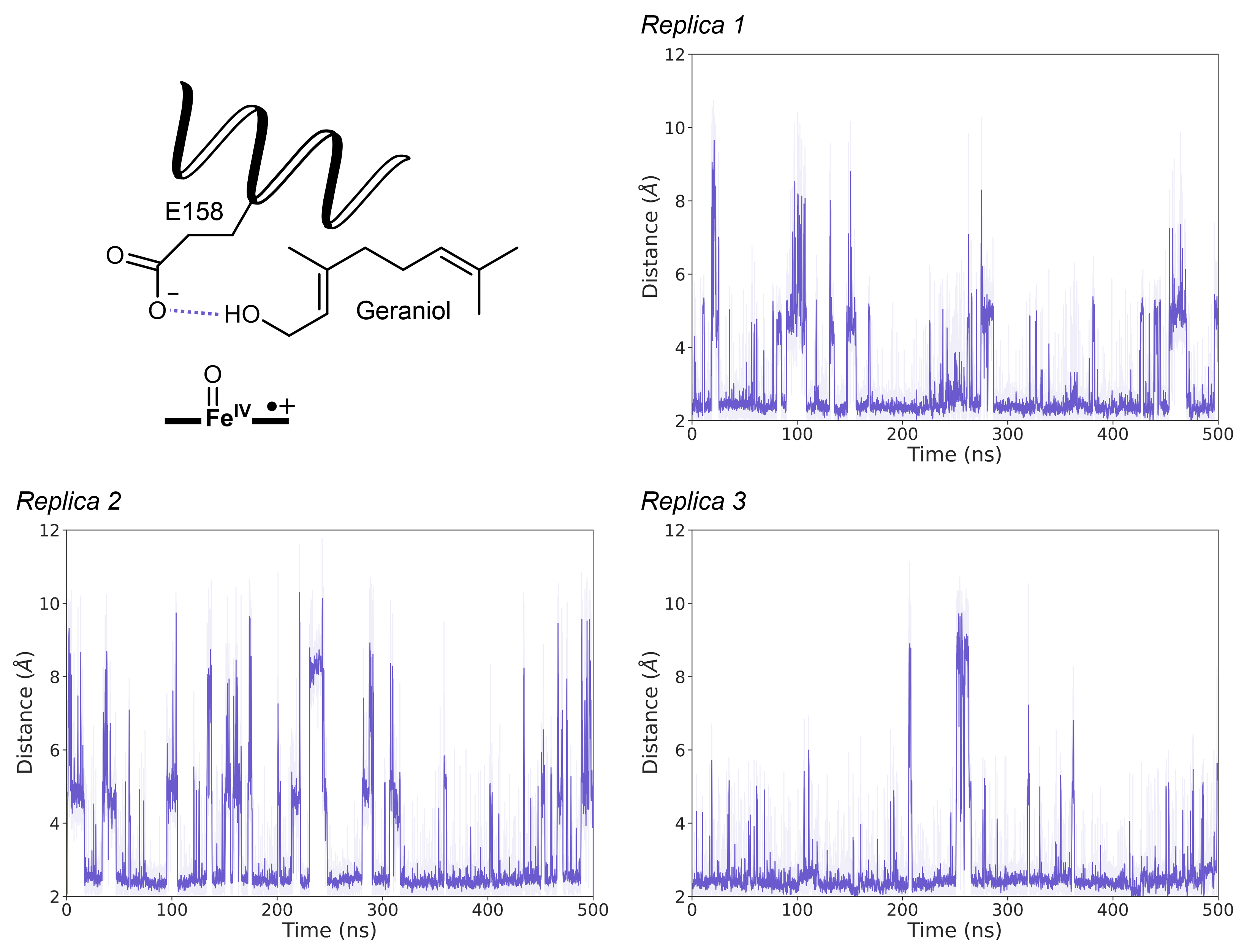

**E)**

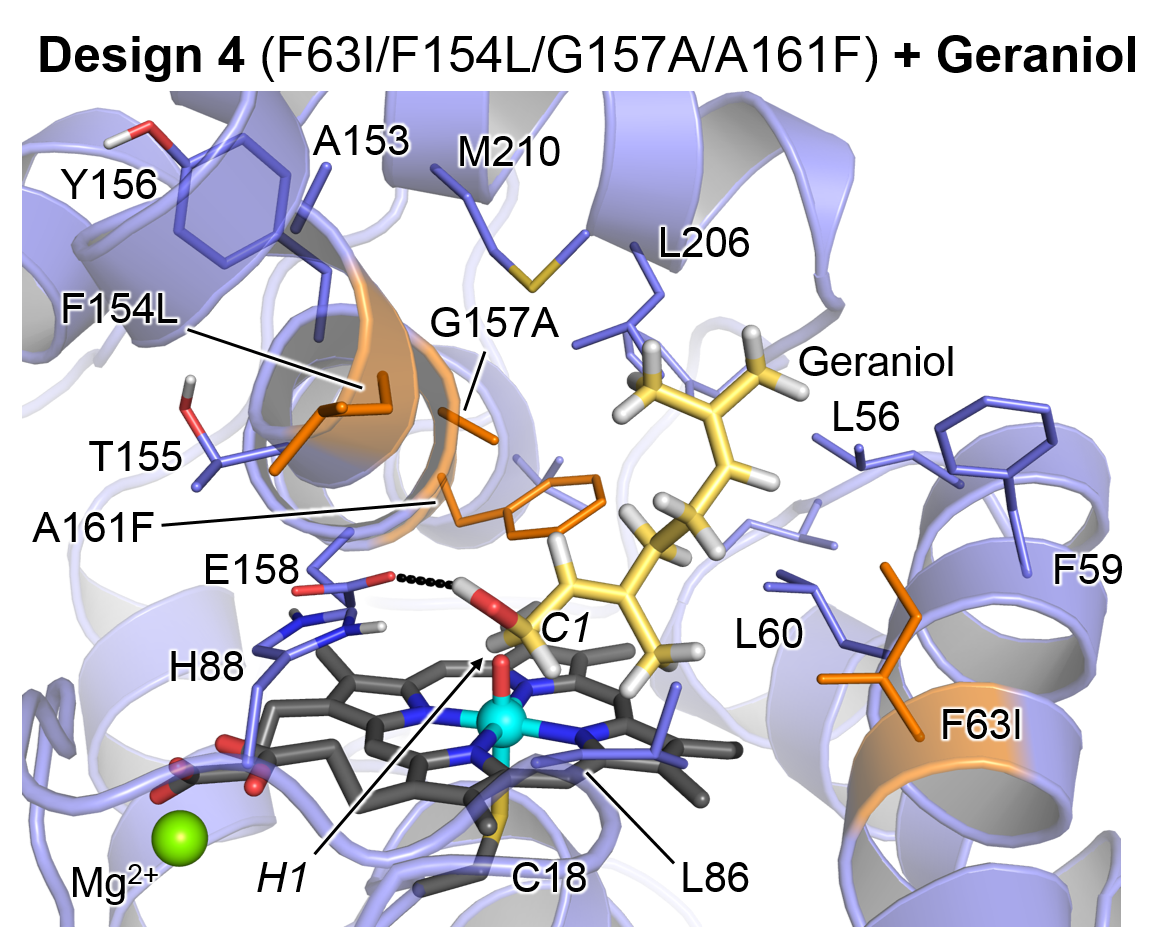

MD simulations with geraniol bound on Design 4 active site indicate that due to active site reshaping induced by the sidechain substitutions (specially by A161A, see **Fig. S8 E**), the C2-C3 double bond of geraniol is located slightly far from the catalytic iron-oxo species compared to C1, where HAT is more geometrically accessible (**Fig. S8-A** and **B**). This is further supported by truncated DFT models with geraniol (**Fig. S6**) where allylic hydroxylation at C1 is intrinsically less energetically demanding than C2-C3 epoxidation.

These simulations also suggest a preference for enantioselective pro-*S* HAT at C1 (**Fig. S8-C**).

Geraniol binds to the active site of Design 4 by anchoring its terminal alcohol group with the catalytic E158 residue (**Figs S8-D** and **E**).

**Fig. S9** Analysis of geraniol (**1**) binding mode in Design 2 (F63L/A153I/F154I/G157A) through substrate-bound MD simulations.

The following analysis allows to characterize which is the preferred binding pose of the substrate in the active site and to elucidate which position of the substrate is more accessible for effective near-attack conformations (NACs) that resemble the geometric requirements of the DFT optimized TS during the MD simulations. Only substrate positions that exhibit a low energetic cost for oxidation as determined by truncated DFT calculations in **Fig. S6** are analyzed in the MD simulations (C2-C3 epoxidation or C1 hydroxylation).

The following geometric parameters are used to characterize the near-attack conformations in the heat map plots: the O–C2 distance and O–C2=C3 angle are used to characterize NACs for effective C2-C3 epoxidation while O–H1 distance and O–H1–C1 angle are used to characterize NACs for effective C1 hydroxylation. The O-C2=C3-C4 dihedral angle is used to determine which enantioface of the C2-C3 bond of the substrate is exposed to the catalytic iron-oxo moiety of Cpd I. Positive dihedral values indicate that the pro-*S* face is exposed while negative values indicate that the pro-*R* face is exposed instead.

The resulting heat maps were obtained from 3 independent MD simulations of 500 ns each (a total of 1500 ns) without any restrain on the substrate. Ideal geometric parameters obtained from DFT Transition State (TS) optimizations (**Fig. S6**) are shown as black dots. A schematic representation of the different geometric parameters used for the heat maps are also provided in each subfigure.

**A)** NACs for C1-HAT leading to geraniol hydroxylation.

**B)** NACs for C2-O formation leading to geraniol epoxidation.

**C)** NACs for pro-*R* and pro-*S* C2-O formation leading to enantioselective geraniol epoxidation.

**D)** Polar interaction between the terminal alcohol group of geraniol and the sidechain of E158.

**E)** Most representative structure of the binding mode of geraniol in Design 2 active site as determined by clustering analysis.

Distances and angles are given in angstroms (Å) and degrees (º), respectively.

**A)**

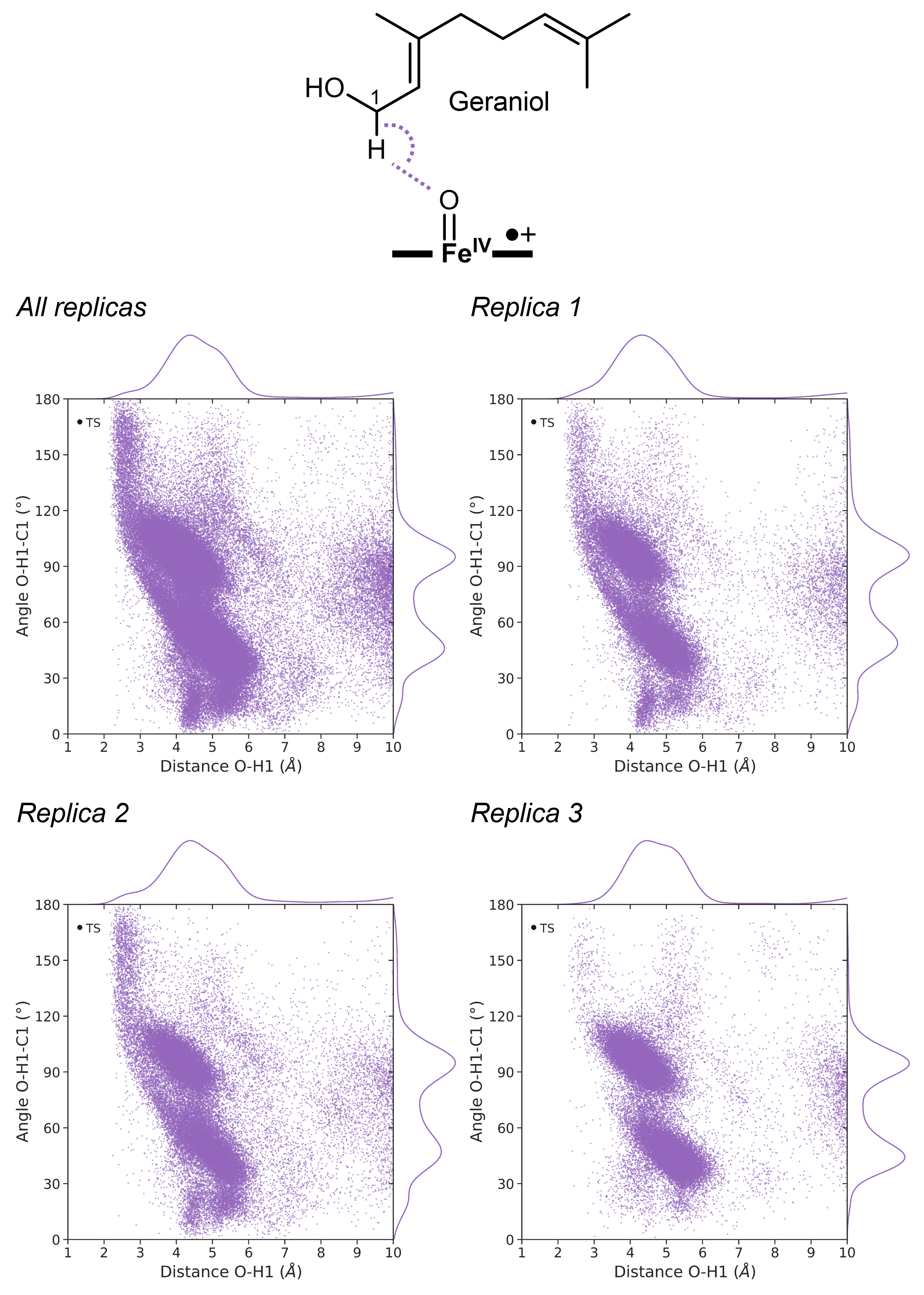

**B)**

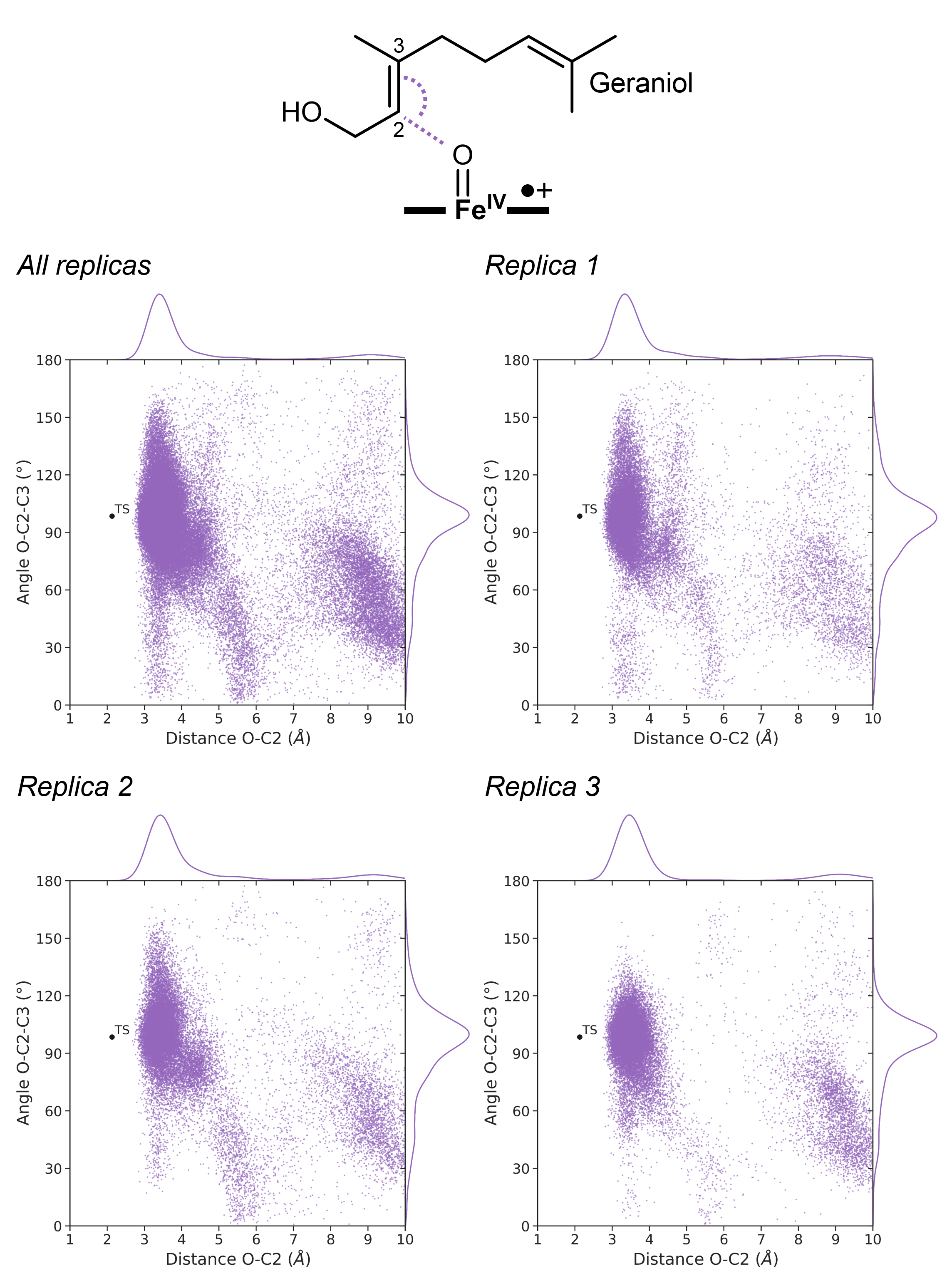

**C)**

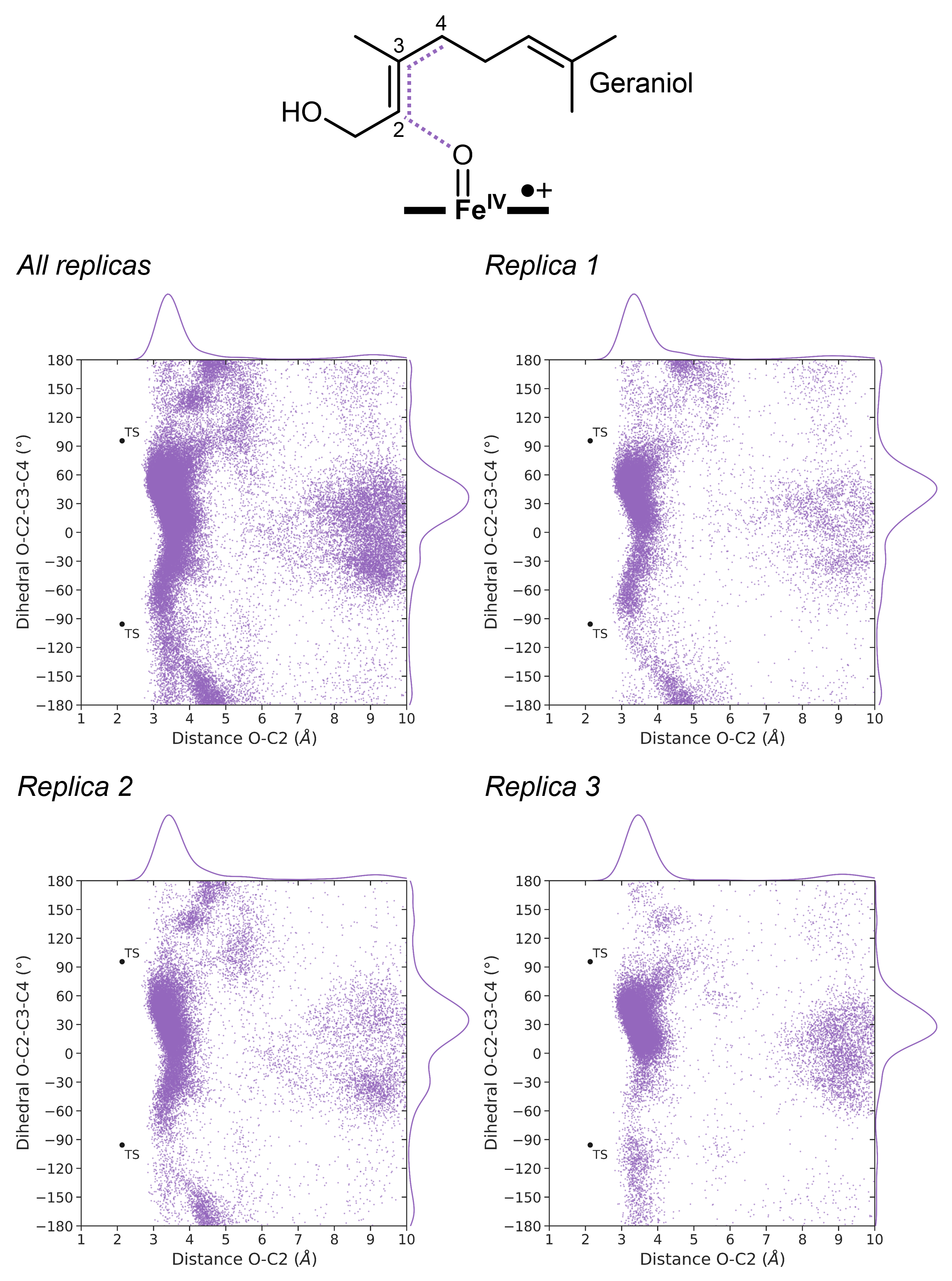

**D)**

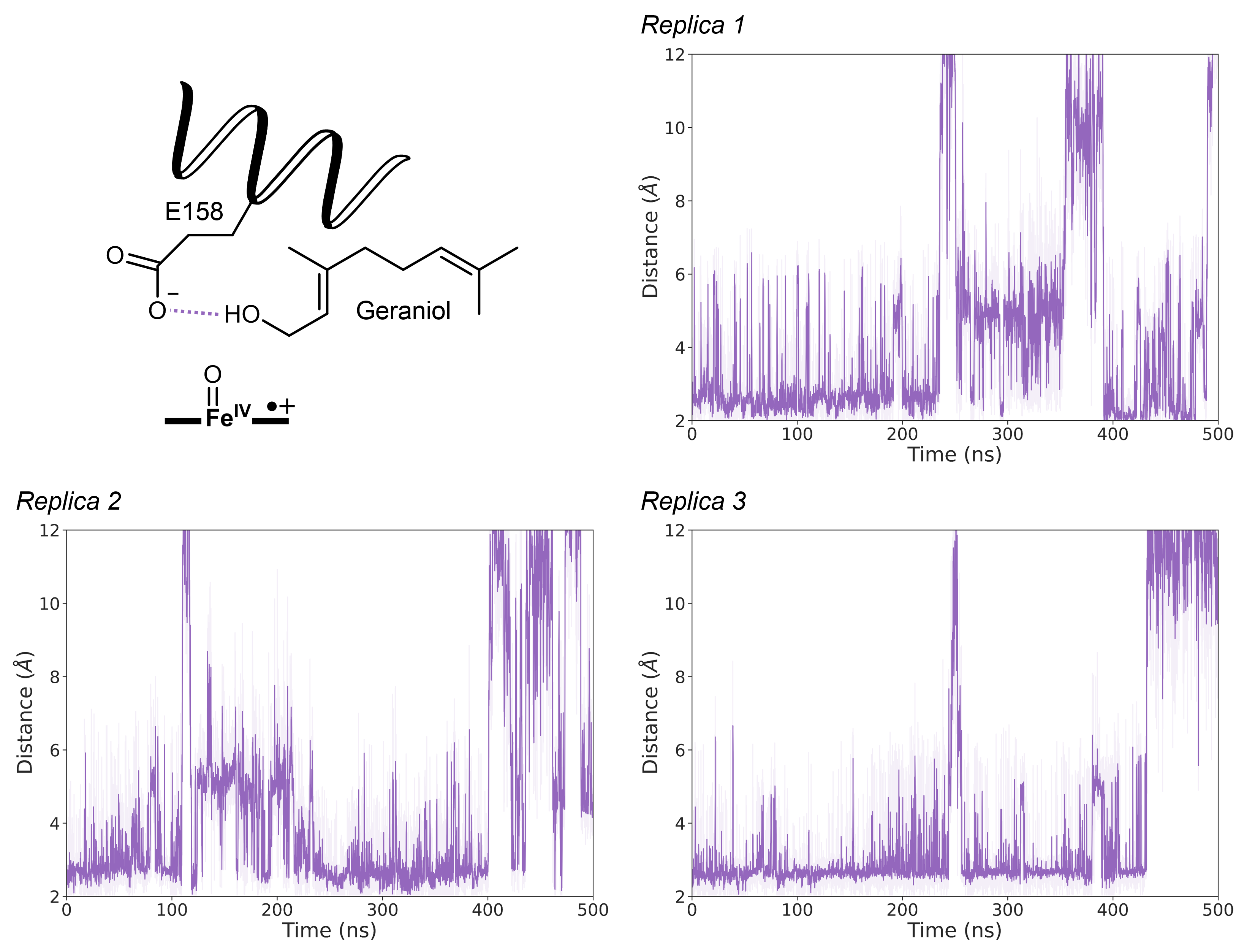

**E)**

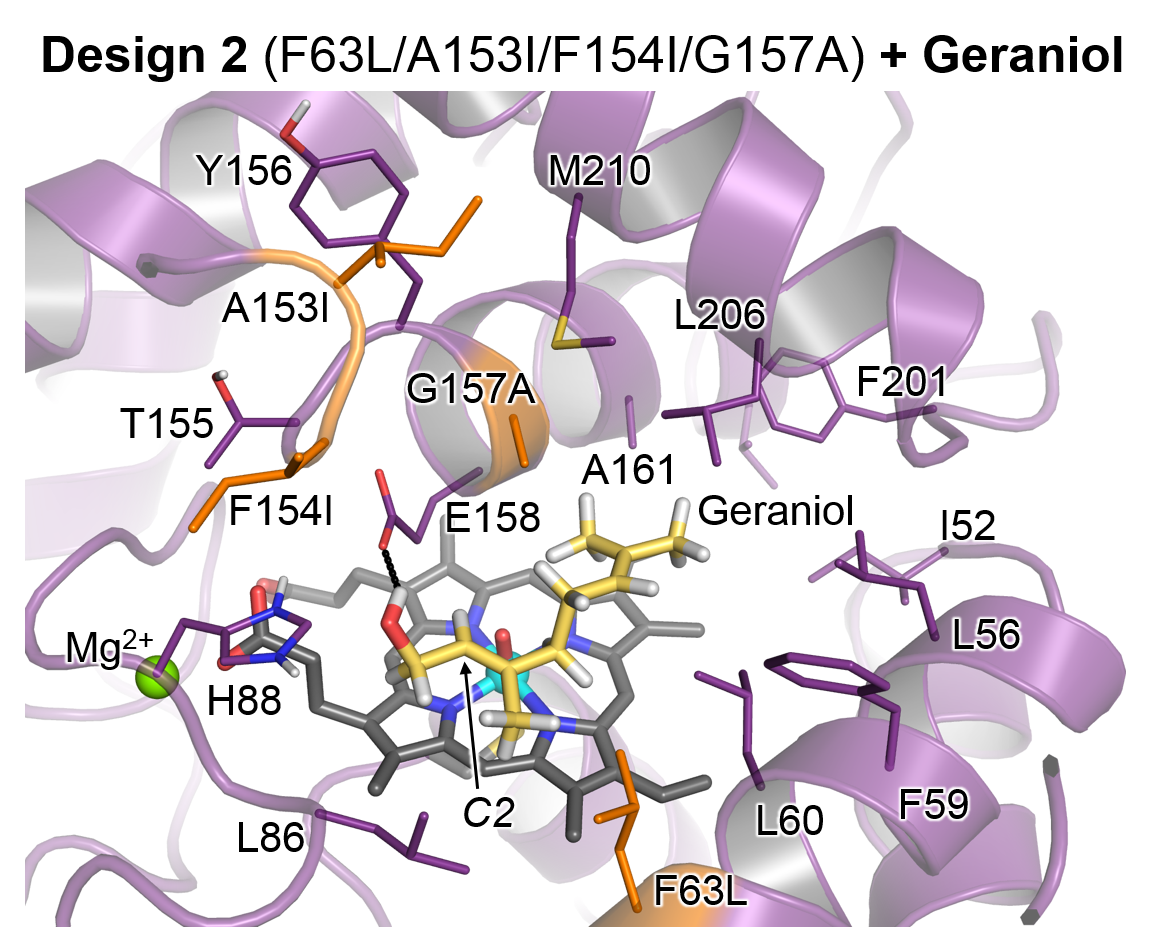

As in Design 4 (see **Fig. S8**), geraniol in Design 2 anchors its hydroxyl group on the catalytic E158 residue (**Figs S9 D** and **E**). However, Design 2 has a less sterically congested active site than Design 4 and geraniol can lay closer to the active Cpd I species, positioning the double C2-C3 bond next to the catalytic iron-oxo group (**Fig. S9 B** and **E**). As a consequence, the C-H bonds at C1 are not geometrically preorganized for effective HAT since they are not pointing to the catalyst (**Fig. S9 A**).

Therefore, even if the truncated DFT models on geraniol (**Fig. S6**) suggest that C1 hydroxylation is energetically preferred over C2-C3 epoxidation, MD simulations indicate that the required near-attack conformations for geraniol hydroxylation at C1 are not significantly sampled on Design 2 active site, thus hindering this intrinsically preferred reactivity.

Analysis of the substrate orientation along the MD simulations suggest that the substrate exposes the C2-C3 double bond to the Cpd I species leading to pro-*S* epoxidation (**Fig. S9-C**).

**Fig. S10** Analysis of nerol (**2**) binding mode in Design 26 (F63I/F154I/G157A/A161L) through substrate-bound MD simulations.

The following analysis allows to characterize which is the preferred binding pose of the substrate in the active site and to elucidate which position of the substrate is more accessible for effective near-attack conformations (NACs) that resemble the geometric requirements of the DFT optimized TS during the MD simulations. Only substrate positions that exhibit a low energetic cost for oxidation as determined by truncated DFT calculations in **Fig. S7** are analyzed in the MD simulations (C2-C3 epoxidation or C1 hydroxylation).

The following geometric parameters are used to characterize the near-attack conformations in the heat map plots: the O–C2 distance and O–C2=C3 angle are used to characterize NACs for effective C2-C3 epoxidation while O–H1 distance and O–H1–C1 angle are used to characterize NACs for effective C1 hydroxylation.

The resulting heat maps were obtained from 3 independent MD simulations of 500 ns each (a total of 1500 ns) without any restraint on the substrate. Ideal geometric parameters obtained from DFT Transition State (TS) optimizations (**Fig. S7**) are shown as black dots. A schematic representation of the different geometric parameters used for the heat maps are also provided in each subfigure.

**A)** NACs for C1-HAT leading to nerol hydroxylation.

**B)** NACs for C2-O formation leading to nerol epoxidation.

**C)** Polar interaction between the terminal alcohol group of nerol and the sidechain of E158.

**D)** Most representative structure of the binding mode of nerol in Design 26 active site as determined by clustering analysis.

Distances and angles are given in angstroms (Å) and degrees (º), respectively.

**A)**

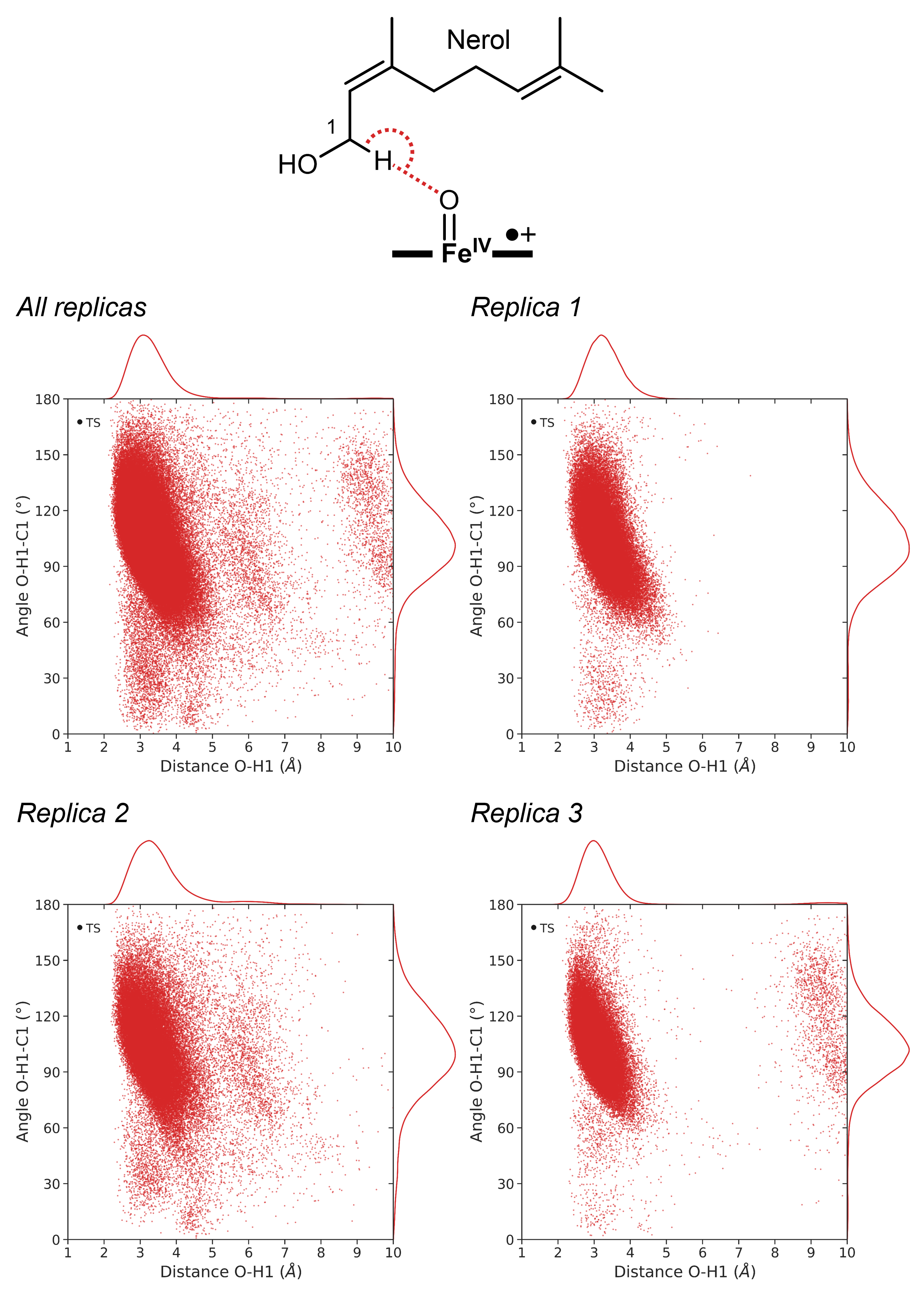

**B)**

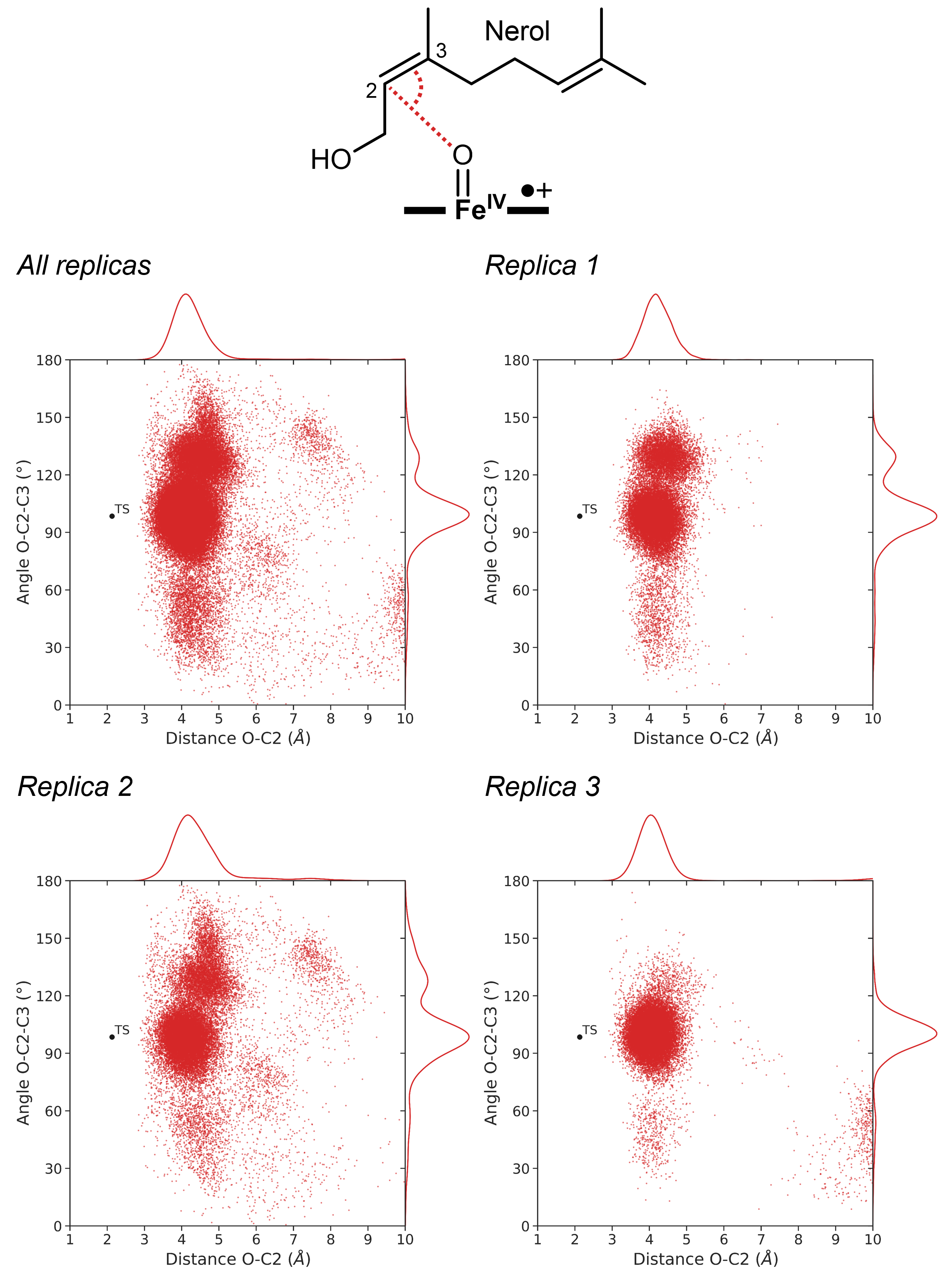

**C)**

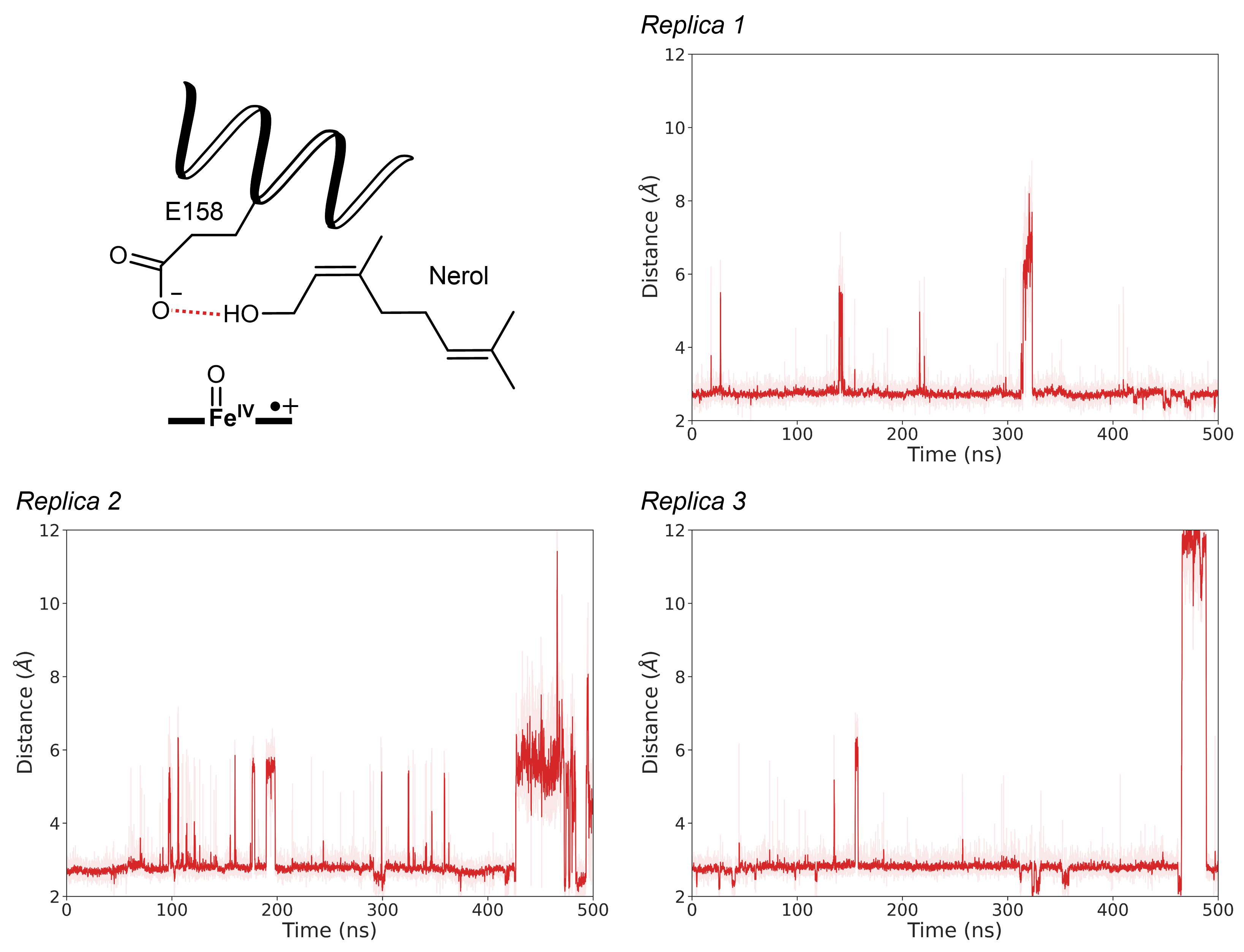

**D)**

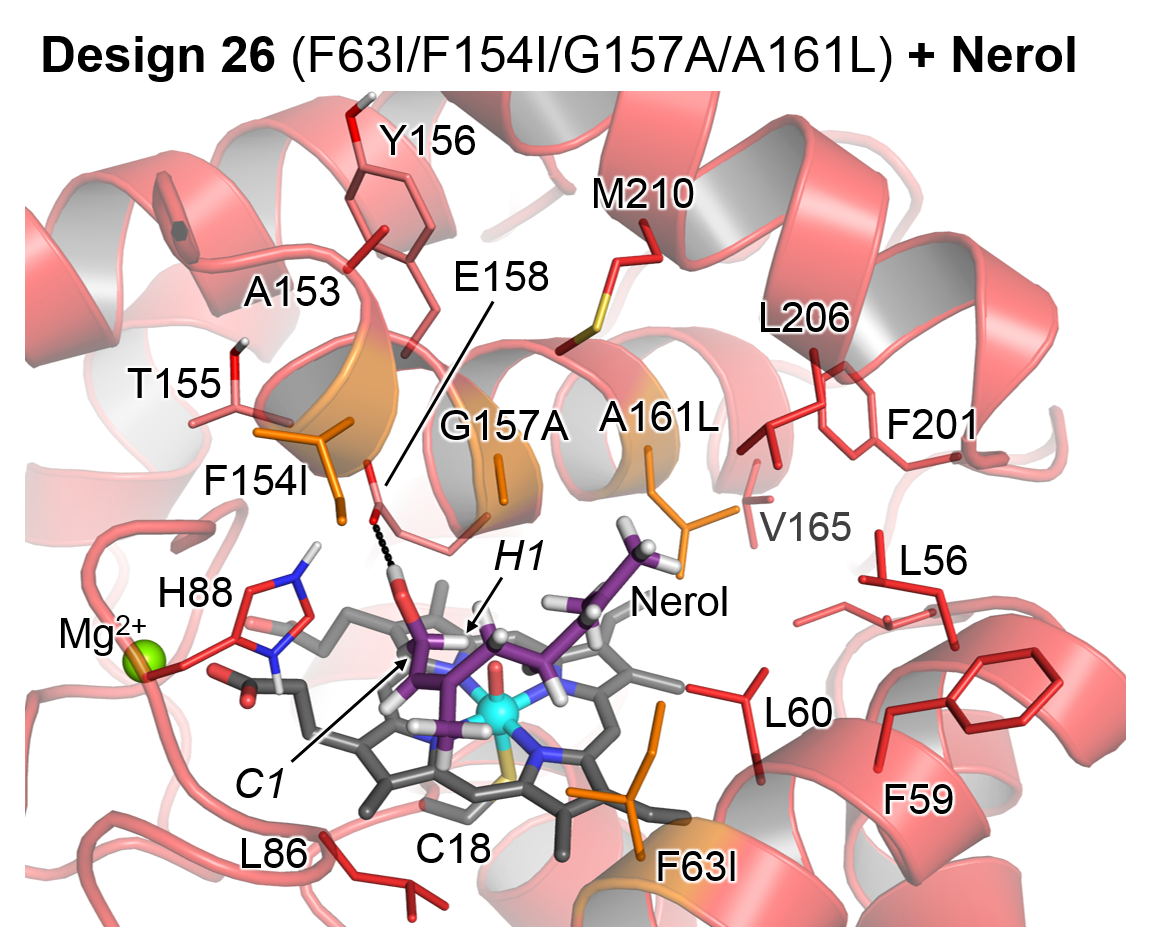

The binding mode of nerol in Design 26 is similar to the binding mode of geraniol in Design 4 E158 (**Fig. S10 D** and **Fig. S8 D**). In both cases, the terminal alcohol group of the substrate interact with E158 (**Fig. S10 C** and **Fig. S8 C**).

In addition, position A161 is also mutated by a residue with a bulky sidechain (leucine in Design 26, phenyalanine in Design 4) and the substrate exposes both the C1-H bonds and the C2-C3 double bond (**Figs S10 A** and **B**). As in the truncated DFT model calculation with geraniol, allylic C1 hydroxylation on nerol is energetically preferred, rationalizing the formation of neral as the major experimental product.

No enantioselective preference for pro-*R* or pro-*S* C1 HAT is observed on the MD simulations with nerol bound in the active site of Design 26.

**Fig. S11** Analysis of nerol (**2**) binding mode in Design 28 (F63L/F154L/Y156L/A161F) through substrate-bound MD simulations.

The following analysis allows to characterize which is the preferred binding pose of the substrate in the active site and to elucidate which position of the substrate is more accessible for effective near-attack conformations (NACs) that resemble the geometric requirements of the DFT optimized TS during the MD simulations. Only substrate positions that exhibit a low energetic cost for oxidation as determined by truncated DFT calculations in **Fig. S7** are analyzed in the MD simulations (C2-C3 epoxidation or C1 hydroxylation).

The following geometric parameters are used to characterize the near-attack conformations in the heat map plots: the O–C2 distance and O–C2=C3 angle are used to characterize NACs for effective C2-C3 epoxidation while O–H1 distance and O–H1–C1 angle are used to characterize NACs for effective C1 hydroxylation. The O-C2=C3-C4 dihedral angle is used to determine which enantioface of the C2-C3 bond of the substrate is exposed to the catalytic iron-oxo moiety of Cpd I. Positive dihedral values indicate that the pro-*S* face is exposed while negative values indicate that the pro-*R* face is exposed instead.

The resulting heat maps were obtained from 3 independent MD simulations of 500 ns each (a total of 1500 ns) without any restraint on the substrate. Ideal geometric parameters obtained from DFT Transition State (TS) optimizations (**Fig. S7**) are shown as black dots. A schematic representation of the different geometric parameters used for the heat maps are also provided in each subfigure.

**A)** NACs for C1-HAT leading to nerol hydroxylation.

**B)** NACs for C2-O formation leading to nerol epoxidation.

**C)** NACs for pro-*R* and pro-*S* C2-O formation leading to enantioselective nerol epoxidation.

**D)** Polar interaction between the terminal alcohol group of nerol and iron-oxo group of Cpd I

**E)** Most representative structure of the binding mode of nerol in Design 28 active site as determined by clustering analysis.

**F)** Probability to find specific interactions between the backbone atoms of E158 and Y162, the hydroxyl group of the substrate, and solvent molecules in the nerol-bound MD simulations with Design 28. The average value between all the replicas is also provided for each interaction. Additionally, the same analysis is performed for the designs in **Figs. S8** to **S10** to facilitate comparison.

Distances and angles are given in angstroms (Å) and degrees (º), respectively.

**A)**

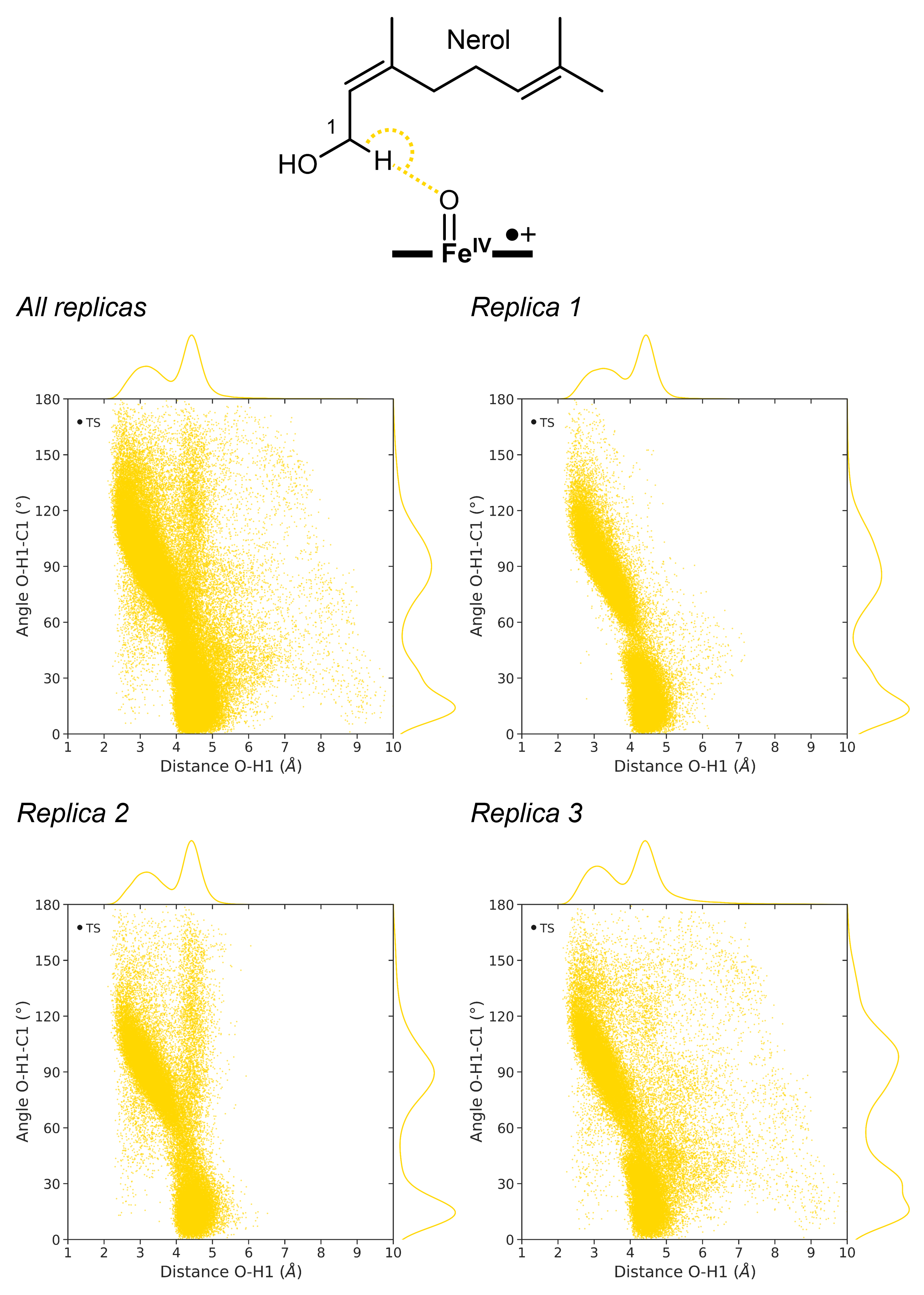

**B)**

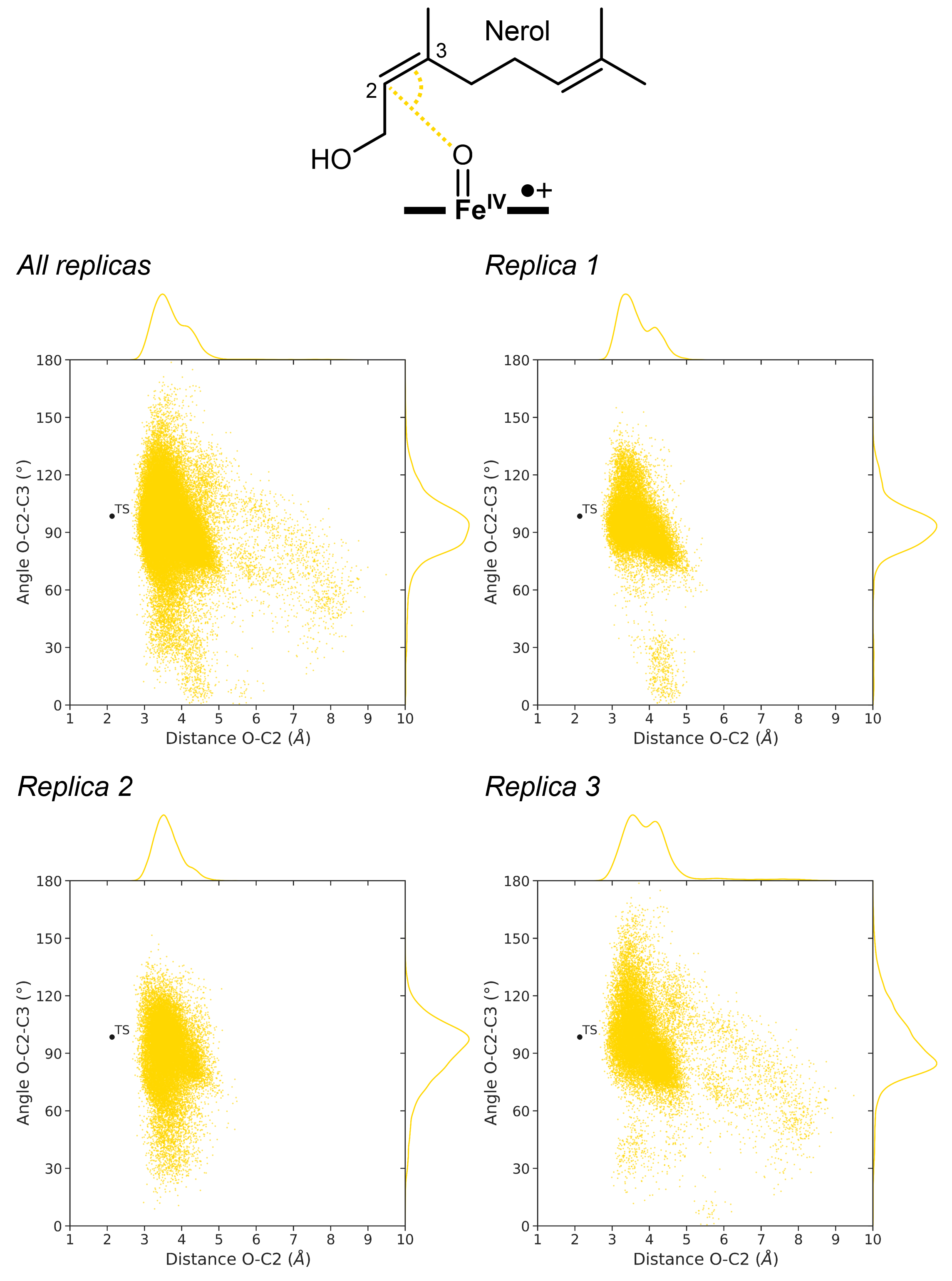

**C)**

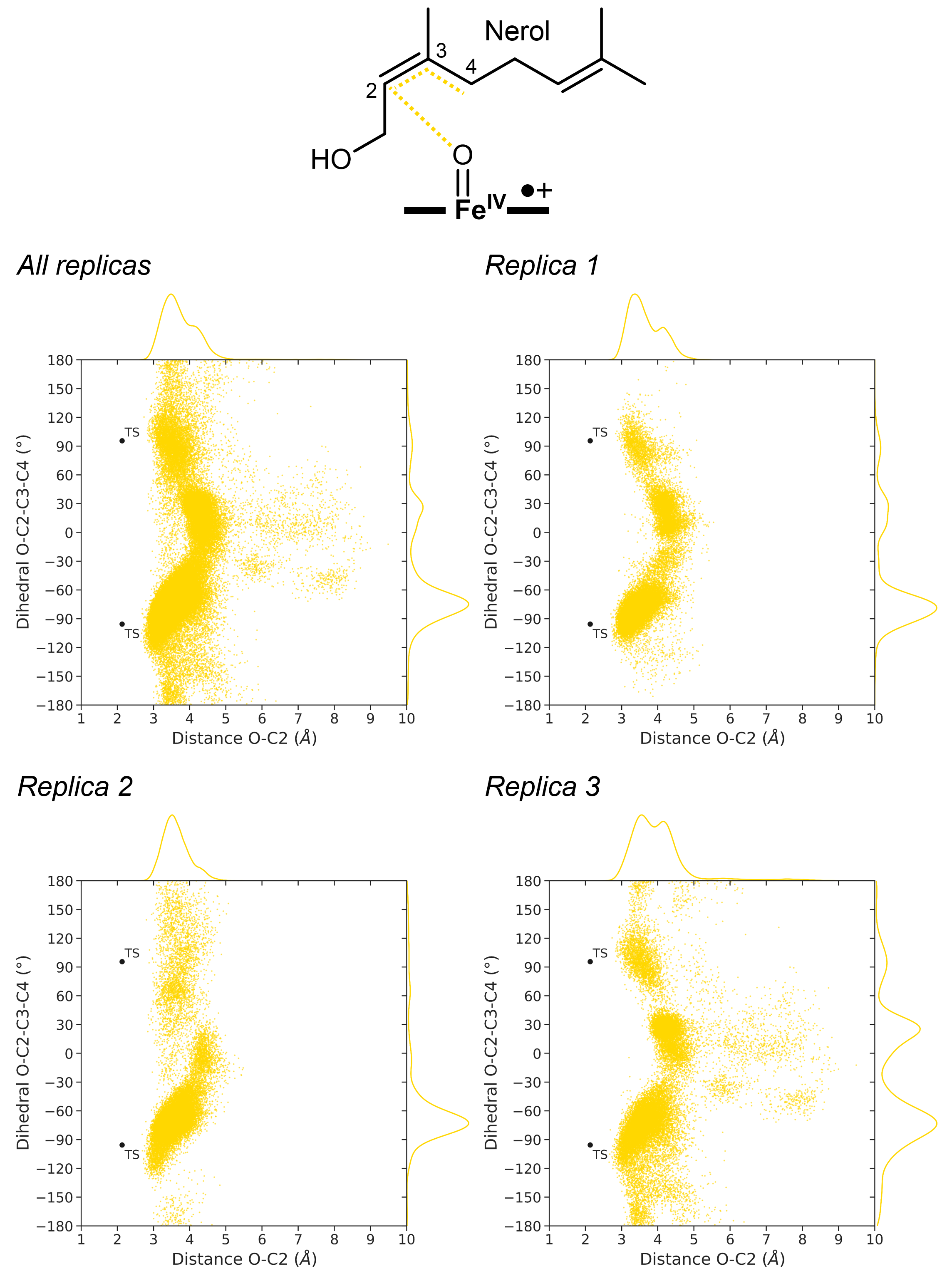

**D)**

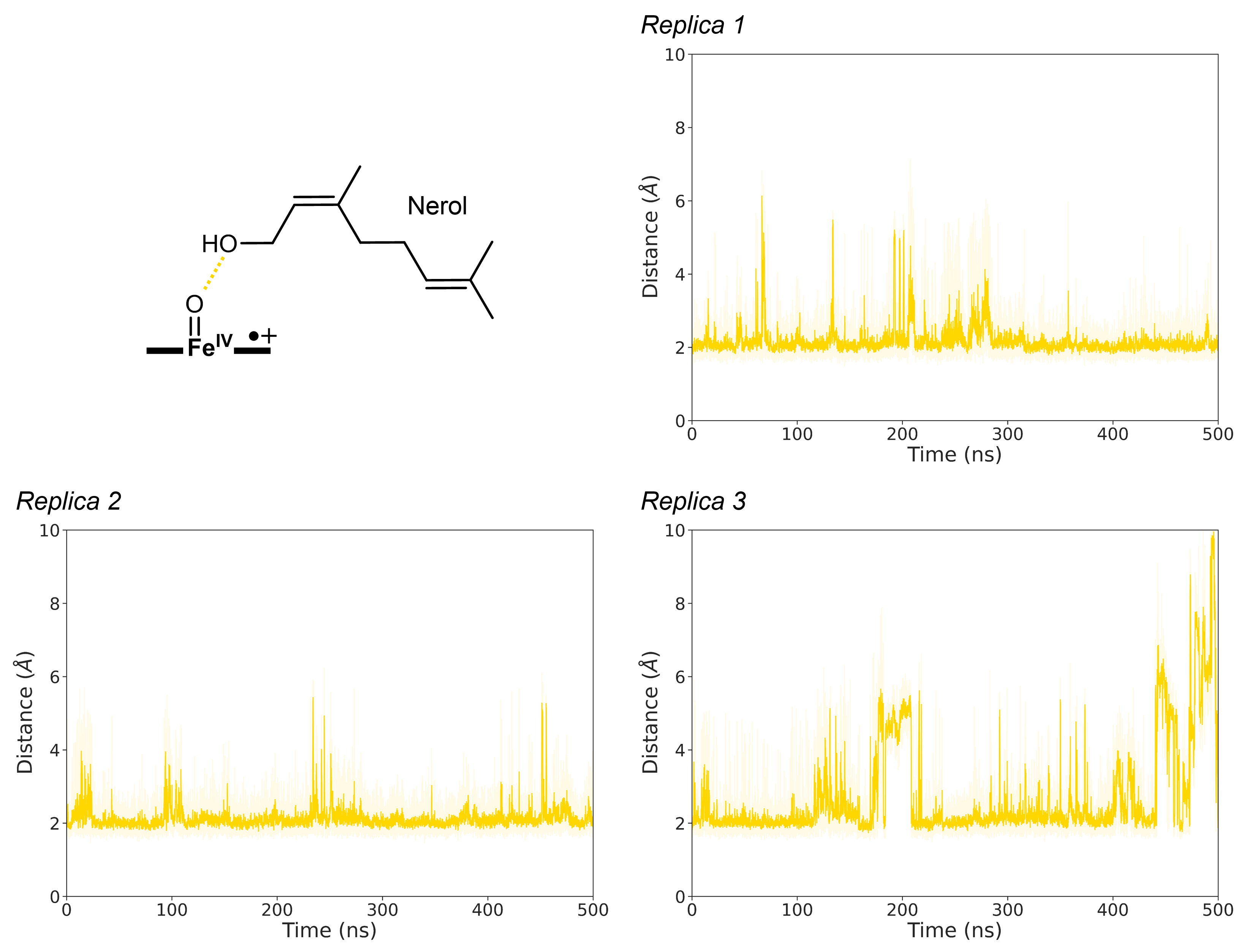

**E)**

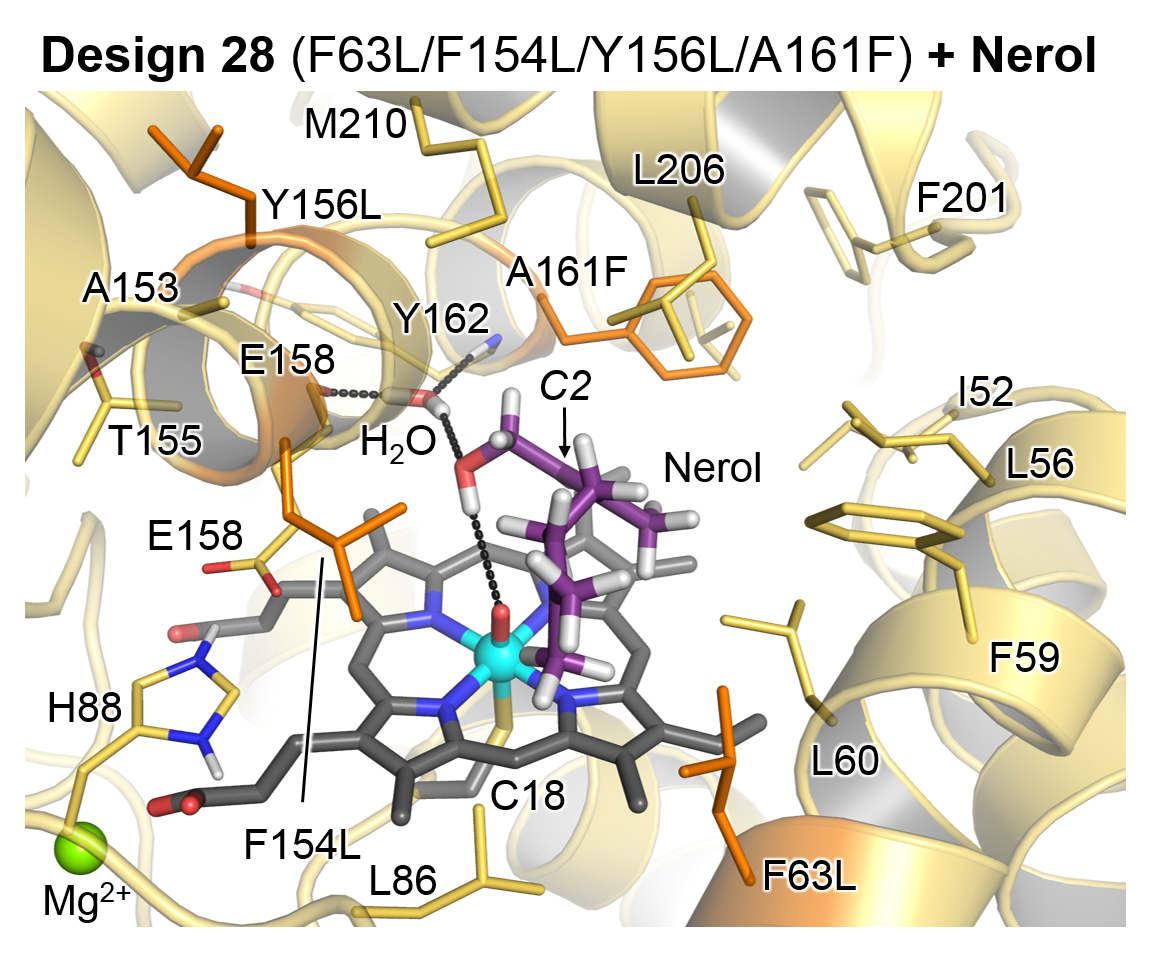

**F)**

| **Design 28 + Nerol** | **Replica 1** | **Replica 2** | **Replica 3** | **Average** |
| --- | --- | --- | --- | --- |
| Direct interaction between E158 - Y162 | 0 | 0 | 0 | 0.00 |
| Bridging water between E158 - Y162 | 0.25 | 0.19 | 0.20 | 0.21 |
| Bridging water between E158 - Nerol | 0.38 | 0.30 | 0.18 | 0.28 |
| Direct interaction between E158 - H_2_O | 0.66 | 0.56 | 0.61 | 0.61 |
| Direct interaction between Nerol - H_2_O | 0.63 | 0.54 | 0.61 | 0.59 |

*Other designs:*

| **Design 26 + Nerol** | **Replica 1** | **Replica 2** | **Replica 3** | **Average** |
| --- | --- | --- | --- | --- |
| Direct interaction between E158 - Y162 | 0.24 | 0.42 | 0.61 | 0.42 |
| Bridging water between E158 - Y162 | 0 | 0 | 0 | 0.00 |
| Bridging water between E158 - Nerol | 0 | 0 | 0 | 0.00 |
| Direct interaction between E158 - H_2_O | 0 | 0 | 0 | 0.00 |
| Direct interaction between Nerol - H_2_O | 0.32 | 0.37 | 0.46 | 0.39 |

| **Design 4 + Geraniol** | **Replica 1** | **Replica 2** | **Replica 3** | **Average** |
| --- | --- | --- | --- | --- |
| Direct interaction between E158 - Y162 | 0.01 | 0.01 | 0.01 | 0.01 |
| Bridging water between E158 - Y162 | 0 | 0 | 0.04 | 0.01 |
| Bridging water between E158 - Geraniol | 0 | 0 | 0 | 0.00 |
| Direct interaction between E158 - H_2_O | 0 | 0 | 0.09 | 0.03 |
| Direct interaction between Geraniol-H_2_O | 0.58 | 0.69 | 0.56 | 0.61 |

| **Design 2 + Geraniol** | **Replica 1** | **Replica 2** | **Replica 3** | **Average** |
| --- | --- | --- | --- | --- |
| Direct interaction between E158 - Y162 | 0.15 | 0.18 | 0.27 | 0.20 |
| Bridging water between E158 - Y162 | 0 | 0 | 0 | 0.00 |
| Bridging water between E158 - Geraniol | 0 | 0 | 0 | 0.00 |
| Direct interaction between E158 - H_2_O | 0 | 0 | 0 | 0.00 |
| Direct interaction between Geraniol-H_2_O | 0.46 | 0.68 | 0.65 | 0.60 |

Nerol binds to Design 28 active site in a novel binding mode not previously detected in any of the previous designs explored by MD simulations in **Figs. S8** to **S10**. This new binding mode is stabilized by an unprecedented direct interaction between the iron-oxo group of Cpd I and the terminal hydroxyl group of the substrate (**Fig. S11 D**) and by the presence of a persistent water molecule bridging the backbone atoms of E158 and Y162 residues, which interacts simultaneously with the alcohol group of nerol (**Figs. S11 E** and **F**). Additional analysis on the previous designs indicates that the presence of this water molecule in Design 28 is due to the mutations introduced in this specific design (mainly by Y156L) rather than an intrinsic feature of all *Mth*UPO designs (**Fig. S11 F**).

As a consequence of this binding pose, the C-H bonds in C1 are not properly oriented to the catalytic moiety (**Fig. S11 A**), positioning C2 in an effective near-attack conformation for C2-C3 epoxidation (**Fig. S11 B**). Analysis of the substrate orientation along the MD simulations suggest that the substrate exposes the C2-C3 double bond to the Cpd I species leading to pro-*R* epoxidation (**Fig. S11 C**).

**Fig. S12** First round of FuncLib evolution toward enantiodivergent *S*- and *R*-4-hydroxy-β-ionone formation starting from *Mth*UPO F63I parent enzyme. See Material and Methods section in the SI for further information about the parameters used for FuncLib. See results discussion in the main text.

**A)** FuncLib evolution round using the F63I *Mth*UPO variant as the parent enzyme. The colour gradient var indicates the Rosetta score difference between the 509 new FuncLib designs and the parent enzyme highlighted in grey (green (red) colour indicates FuncLib designs that have better (worse) Rosetta scores than the parent enzyme). The hits found in our previous study^35^ (Münch et al. 2024, Table 1) are also highlighted with black lines.

**B)** Summary table of the hits detected in our previous study^35^ (Münch et al. 2024, Table 1), the mutations that were introduced, the ordering in the FuncLib run, the FuncLib Rosetta score and the enantiomeric ratio (e.r.) are provided. Entries in bold exhibit the highest enantiomeric ratio improvements.

**A)**

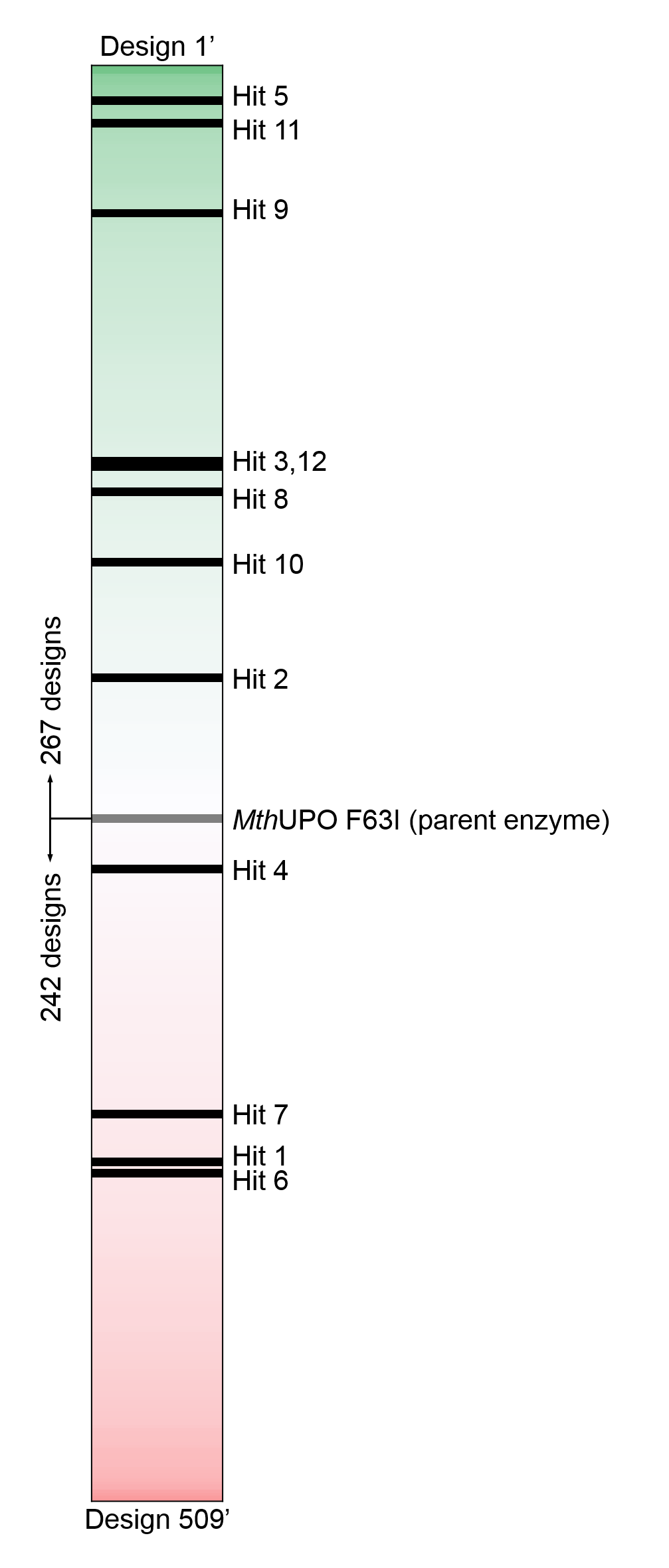

**B)**

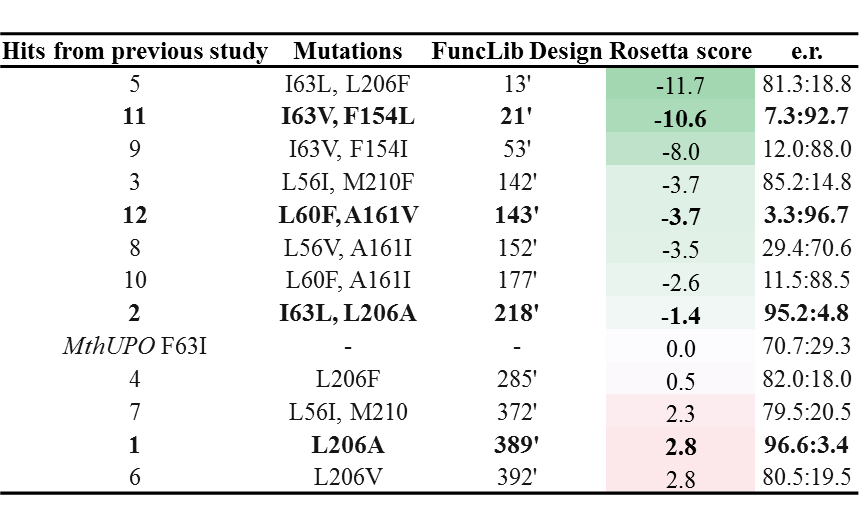

**Fig. S13** Second round of FuncLib evolution toward improvement of the *S*-4-hydroxy-β-ionone formation activity starting from *Mth*UPO L60F, F63I, A161V parent enzyme. See Material and Methods section in the SI for further information about the parameters used for FuncLib. See results discussion in the main text.

**A)** FuncLib evolution round using the L60F, F63I, A161V *Mth*UPO variant as the parent enzyme. The color gradient var indicates the Rosetta score difference between the 197 new FuncLib designs, and the parent enzyme highlighted in gray (green (red) color indicates FuncLib designs that have better (worse) Rosetta scores than the parent enzyme). The hits found in our previous study ^35^ (Münch et al. 2024, Table 2) are also highlighted with black lines.

**B)** Summary table of the hits detected in our previous study ^35^ (Münch et al. 2024, Table 1), the mutations that were introduced, the ordering in the FuncLib run, the FuncLib Rosetta score and the enantiomeric ratio (e.r.) are provided. Entry in bold exhibits the highest enantiometic ratio improvements.

**A)**

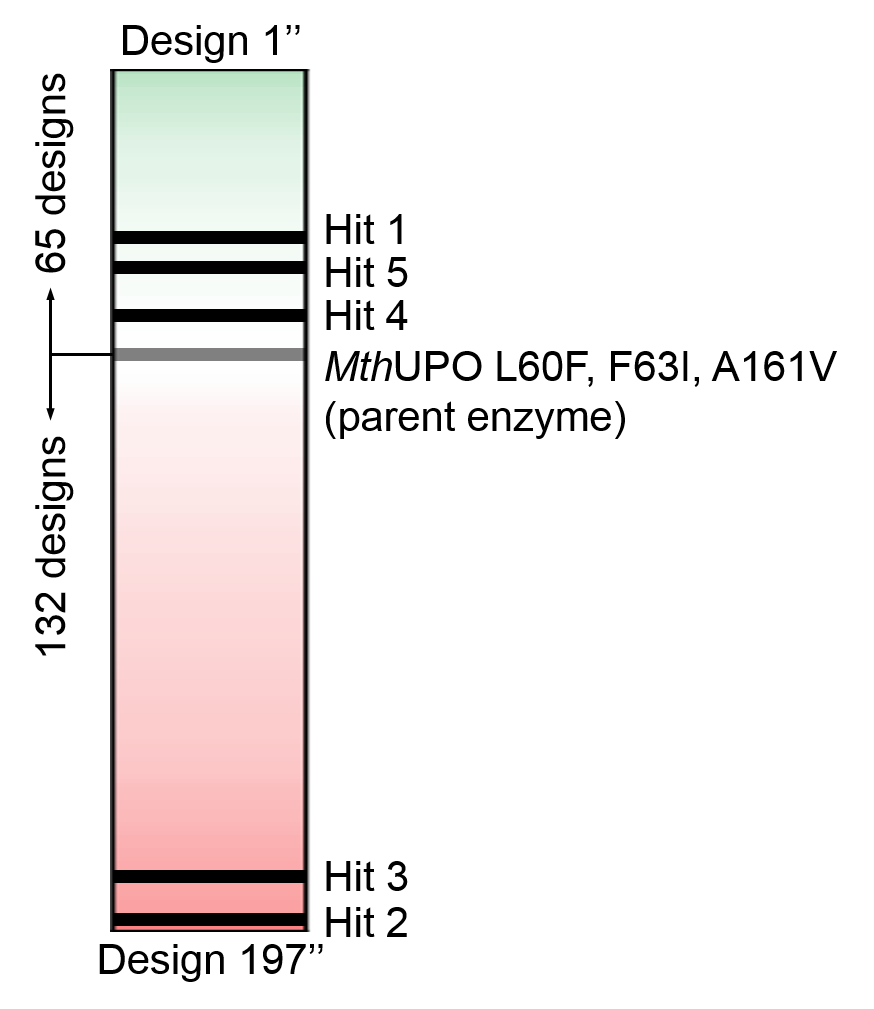

**B)**

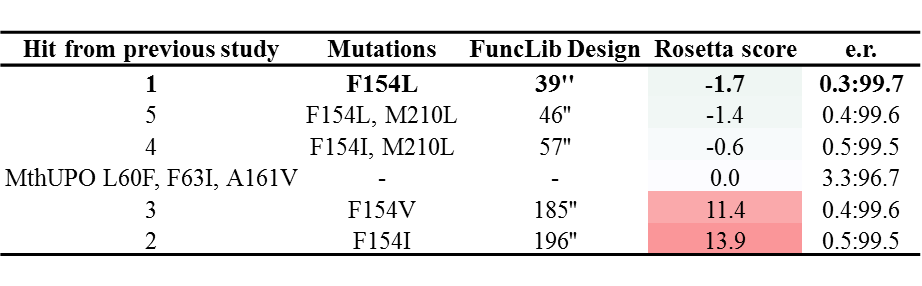

**Fig. S14** Design 18 enzyme helical core modelling with Chai-1 docked with several substrates compared to FuncLib design 18.

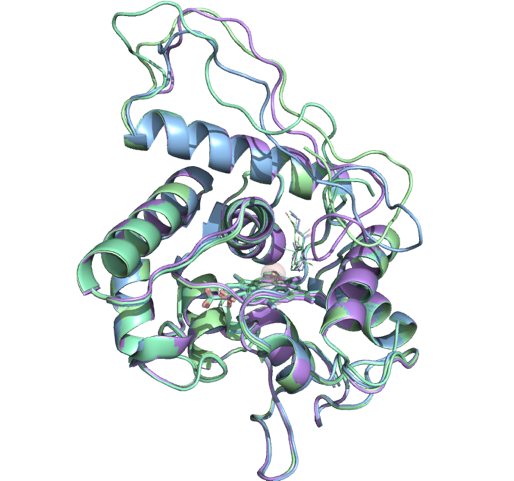

**^1^H-NMR**

Isopiperitenol

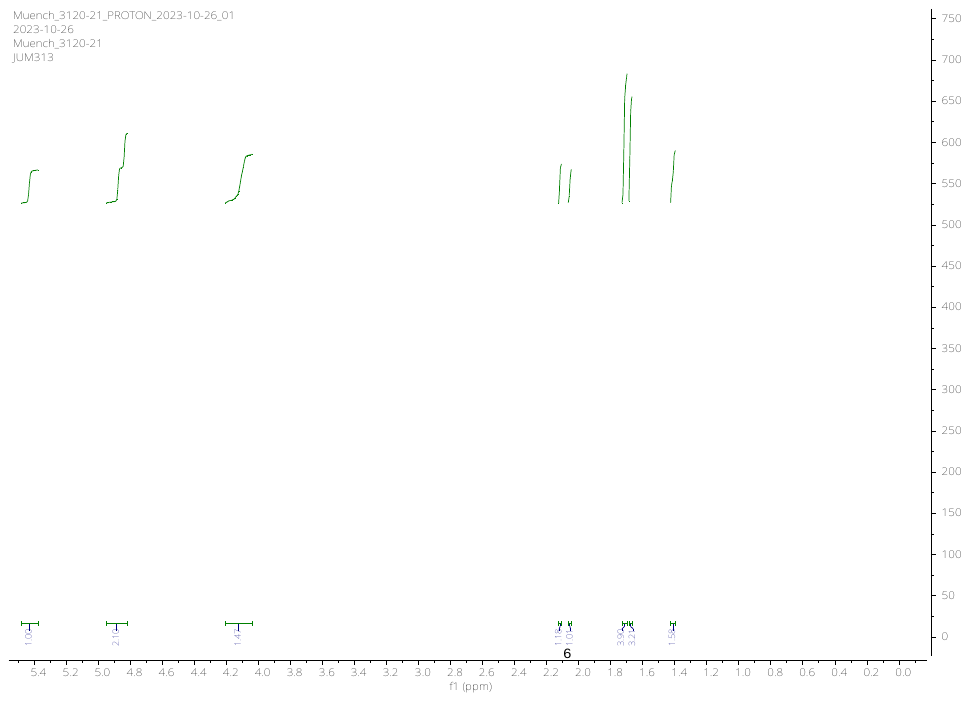

2,3-epoxy nerol

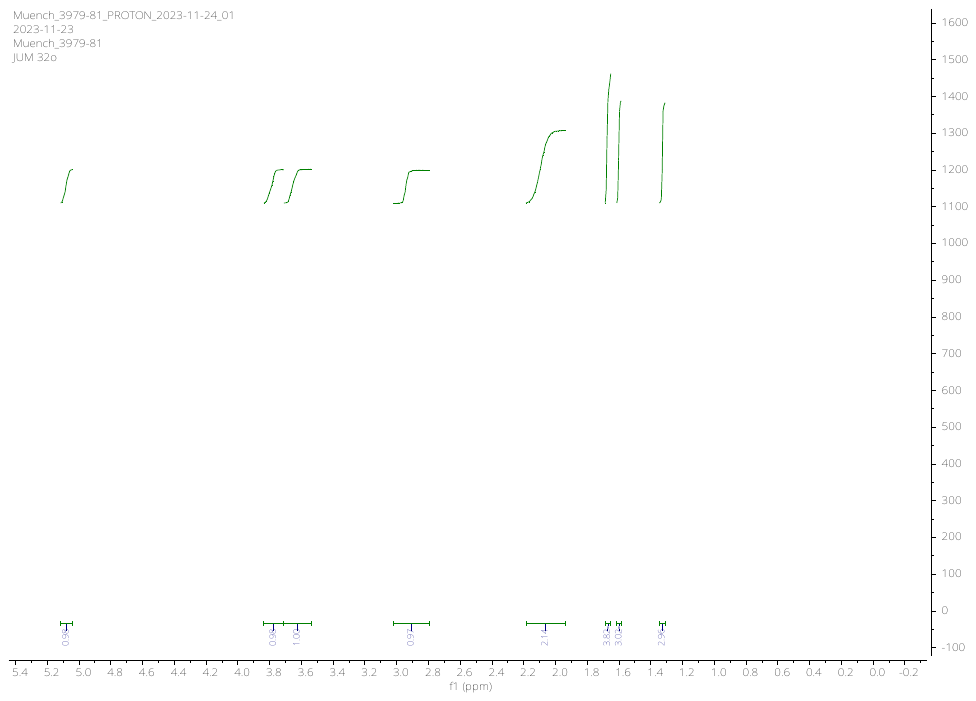
